# Inhibitors of dihydroorotate dehydrogenase synergize with the broad antiviral activity of 4′-fluorouridine

**DOI:** 10.1101/2024.10.05.616778

**Authors:** Leon Schrell, Hannah L. Fuchs, Antje Dickmanns, David Scheibner, Judith Olejnik, Adam J. Hume, Wencke Reineking, Theresa Störk, Martin Müller, Annika Graaf-Rau, Sandra Diederich, Stefan Finke, Wolfgang Baumgärtner, Elke Mühlberger, Anne Balkema-Buschmann, Matthias Dobbelstein

## Abstract

RNA viruses present a constant threat to human health, often with limited options for vaccination or therapy. Notable examples include influenza viruses and coronaviruses, which have pandemic potential. Filo- and henipaviruses cause more limited outbreaks, but with high case fatality rates. All RNA viruses rely on the activity of a virus-encoded RNA-dependent RNA polymerase (RdRp). An antiviral nucleoside analogue, 4′-Fluorouridine (4′-FlU), targets RdRp and diminishes the replication of several RNA viruses, including influenza A virus and SARS-CoV-2, through incorporation into nascent viral RNA and delayed chain termination. However, the effective concentration of 4′-FlU varied among different viruses, raising the need to fortify its efficacy. Here we show that inhibitors of dihydroorotate dehydrogenase (DHODH), an enzyme essential for pyrimidine biosynthesis, can synergistically enhance the antiviral effect of 4′-FlU against influenza A viruses, SARS-CoV-2, henipaviruses, and Ebola virus. Even 4′-FlU-resistant mutant influenza A virus was re-sensitized towards 4′-FlU by DHODH inhibition. The addition of uridine rescued influenza A virus replication, strongly suggesting uridine depletion as a mechanism of this synergy. 4′-FlU was also highly effective against SARS-CoV-2 in a hamster model of COVID. We propose that the impairment of endogenous uridine synthesis by DHODH inhibition enhances the incorporation of 4′-FlU into viral RNAs. This strategy may be broadly applicable to enhance the efficacy of pyrimidine nucleoside analogues for antiviral therapy.

**Graphical Abstract:** 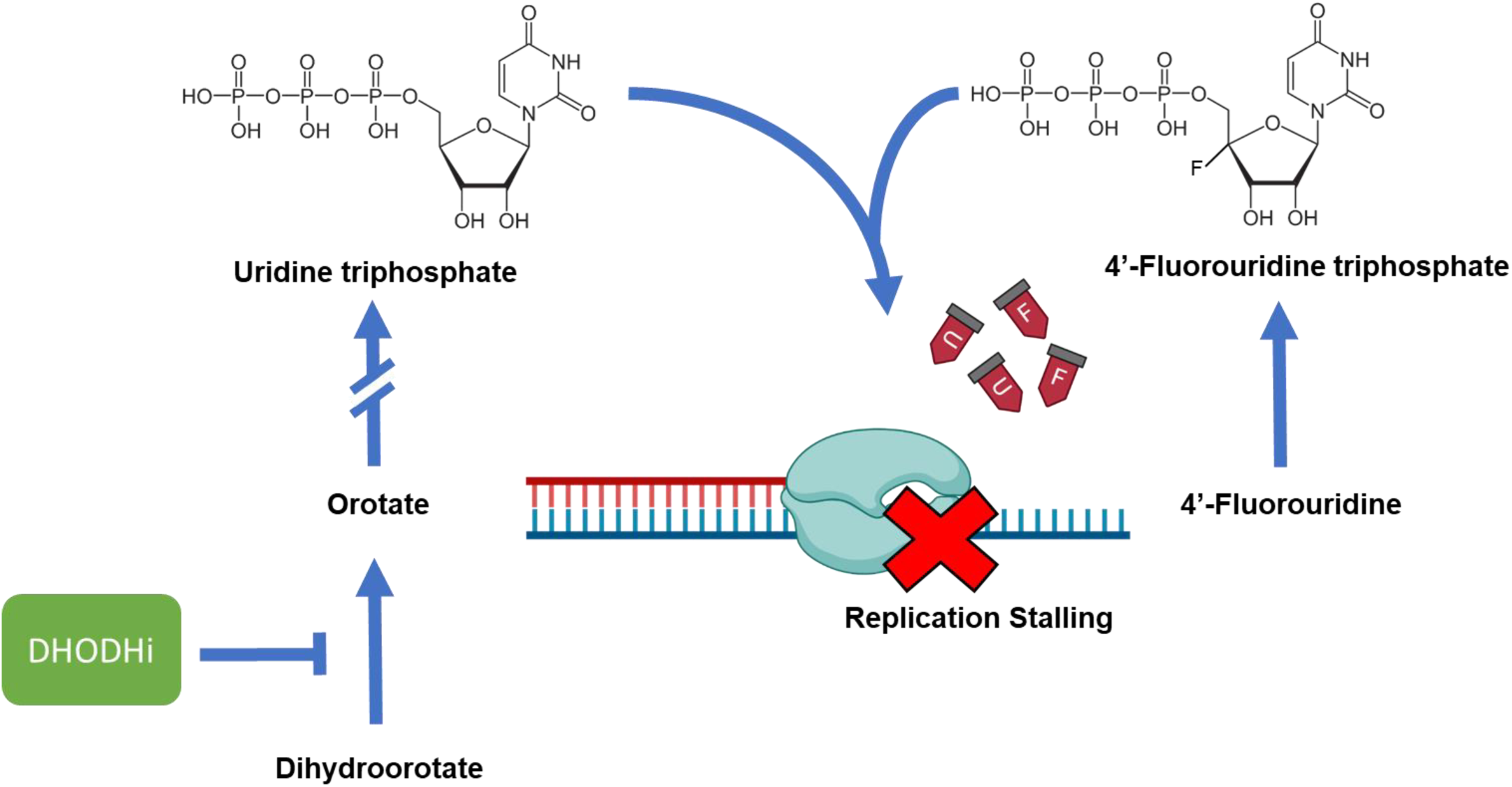

**HIGHLIGHTS:**

- Strong synergy of DHODH inhibitors with 4′-FlU
- Activity of the combination against previously resistant influenza virus
- Broadly active combination against a diverse set of RNA viruses
- Successful targets include highly pathogenic Ebola and Nipah viruses

## INTRODUCTION

RNA viruses are a threat to global health, causing epidemics or even pandemics, with annual death tolls in the range of millions (Garcia-Blanco et al., 2022; Meganck and Baric, 2021). Human influenza A viruses (IAV) are characterized by seasonal spread, with point mutations and antigenic drift occurring each year. Occasionally, the reassortment of genome segments from different virus subtypes gives rise to viruses with novel traits and antigenic shift, causing devastating pandemics (Krammer et al., 2018). Similarly, coronaviruses often contribute to seasonal common cold symptoms but also carry the potential of causing pandemics, such as COVID-19 (Hillary and Ceasar, 2023). The rapid spread of such viruses makes timely vaccine development challenging. Other emerging and re-emerging RNA viruses, including filoviruses such as Ebola virus (EBOV) (Jacob et al., 2020), and henipaviruses, especially the highly pathogenic Nipah virus (NiV) (Li et al., 2023), induce outbreaks with high case fatality rates, and effective vaccines are not available for most of these viruses. The highly pathogenic filo- and henipaviruses are prioritized by the World Health Organization for research and development in emergency contexts (Sweileh, 2017). In these scenarios, broad-spectrum therapeutics could strongly contribute to the management of these infections and to the containment of epidemics.

Antiviral nucleoside analogues are incorporated into the viral genome during virus replication. They were initially developed to antagonize the replication of herpesviruses and human immunodeficiency virus (HIV), inducing chain termination of the nascent DNA strand (De Clercq, 2004). In the meantime, similar nucleosides became available to disrupt RNA virus replication (Geraghty et al., 2021). A prominent nucleoside analogue is molnupiravir, the prodrug of N4-hydrxycytidine (NHC) (Agostini et al., 2019). NHC is active against IAV (Toots et al., 2019) and Severe Acute Respiratory Syndrome Coronavirus, SARS-CoV-2 (Sheahan et al., 2020; Wahl et al., 2021), as well as henipaviruses and EBOV (Lo et al., 2018). Moreover, we (Stegmann et al., 2022) and others (Schultz et al., 2022) reported earlier that the inhibition of endogenous pyrimidine synthesis further increases the efficacy of NHC. This is achievable by inhibitors of dihydroorotate dehydrogenase (DHODH), an enzyme that catalyzes an essential step of pyrimidine synthesis. DHODH inhibitors are already available for clinical use (Zheng et al., 2022), and their combination with NHC synergistically inhibits SARS-CoV-2 replication (Stegmann et al., 2022). Moreover, DHODH inhibitors are active against IAV (Li et al., 2024; Sibille et al., 2022; Xiong et al., 2020) and EBOV (Gong et al., 2021; Luthra et al., 2018; Martin et al., 2018).

NHC has given rise to safety concerns because of its mode of action. Instead of causing chain termination, NHC promiscuously forms base pairs with guanine, but also with adenine (Agostini et al., 2019; Gordon et al., 2021; Kabinger et al., 2021). This incorporation of NHC into viral RNA leads to multiple mutations, which are believed to counteract further virus spread by ‘error catastrophe’. However, treatment using inefficient concentrations and/or application frequencies of Molnupiravir/NHC might induce the accumulation of virus mutants with gains of function, e.g. immune escape. Our previous work corroborates this notion at least in an in vitro setting (Kumar et al., 2024; Zibat et al., 2023). Thus, nucleoside analogues which cause direct or delayed termination might represent a better choice for antiviral therapy.

More recently, the orally available drug candidate 4′-fluorouridine (4′-FlU; EIDD-2749) has demonstrated activity against IAV (Lieber et al., 2023) as well as SARS-CoV-2 (Sourimant et al., 2022), without apparent mutagenicity. Rather, its mechanism of action is delayed chain termination (Lieber and Plemper, 2022). 4′-FlU has also proven effective against a panel of mononegaviruses including respiratory syncytial virus (Sourimant et al., 2022) and a mouse-adapted Heartland virus, which belongs to the order *Bunyavirales* (Westover et al., 2024). However, its efficacy varied between viruses, raising the question of whether this drug could also be fortified by combinatorial treatment with other compounds.

Here we show that the combination of 4′-FlU with DHODH inhibitors strongly synergizes to repress the replication of RNA viruses from four different virus families: IAV, SARS-CoV-2, henipaviruses, and EBOV. Thus, inhibiting pyrimidine biosynthesis strongly augments the efficacy of a broad-range antiviral nucleoside analogue and potentially expands its use against other RNA viruses with epidemic or even pandemic potential.

## RESULTS

### 4′-FlU synergizes with DHODH inhibitors to diminish influenza A virus replication

To test if DHODH inhibitors enhance the impact of 4′-FlU on IAV replication, we infected Madin-Darby canine kidney II cells (termed MDCK cells from here on) with IAV, H1N1 strain A/Puerto Rico/1934 (H1N1/PR/34). One hour post infection (hpi), the cells were treated with 4′-FlU and/or the DHODH inhibitors teriflunomide (Greene et al., 1995), brequinar (Chen et al., 1986), or BAY 2402234 (Christian et al., 2019) in different concentrations for 48 h. We quantified the amount of viral RNA in the supernatant of the infected cells by qRT-PCR. Strikingly, combinations of 4′-FlU with DHODH inhibitors reduced the amount of released viral RNA up to 100-fold more profoundly than each drug alone (Figures 1A; S1, S2). This was reflected by synergy scores of more than 20, where scores >10 are indicative of synergistic drug activity, i.e., more than additive effects. The decrease in RNA levels observed with the drug combination was also mirrored in the overall reduction of viral titers. While single drug treatment resulted in a reduction of viral titers of less than two logs, the combination suppressed virus propagation far more strongly, approaching nearly undetectable levels (Figure 1B). The drugs did not display detectable cytotoxicity, as determined by the release of lactate dehydrogenase (Figure 1C). Synergistic effects were also observed when performing the experiments in Calu-3 cells (Figures 1D, S3), which are derived from neoplastic lung cells and represent a model system for the infected respiratory tract (Harcourt et al., 2011). These data suggest that 4′-FlU synergizes with DHODH inhibitors to efficiently inhibit IAV replication.

**Figure 1.**
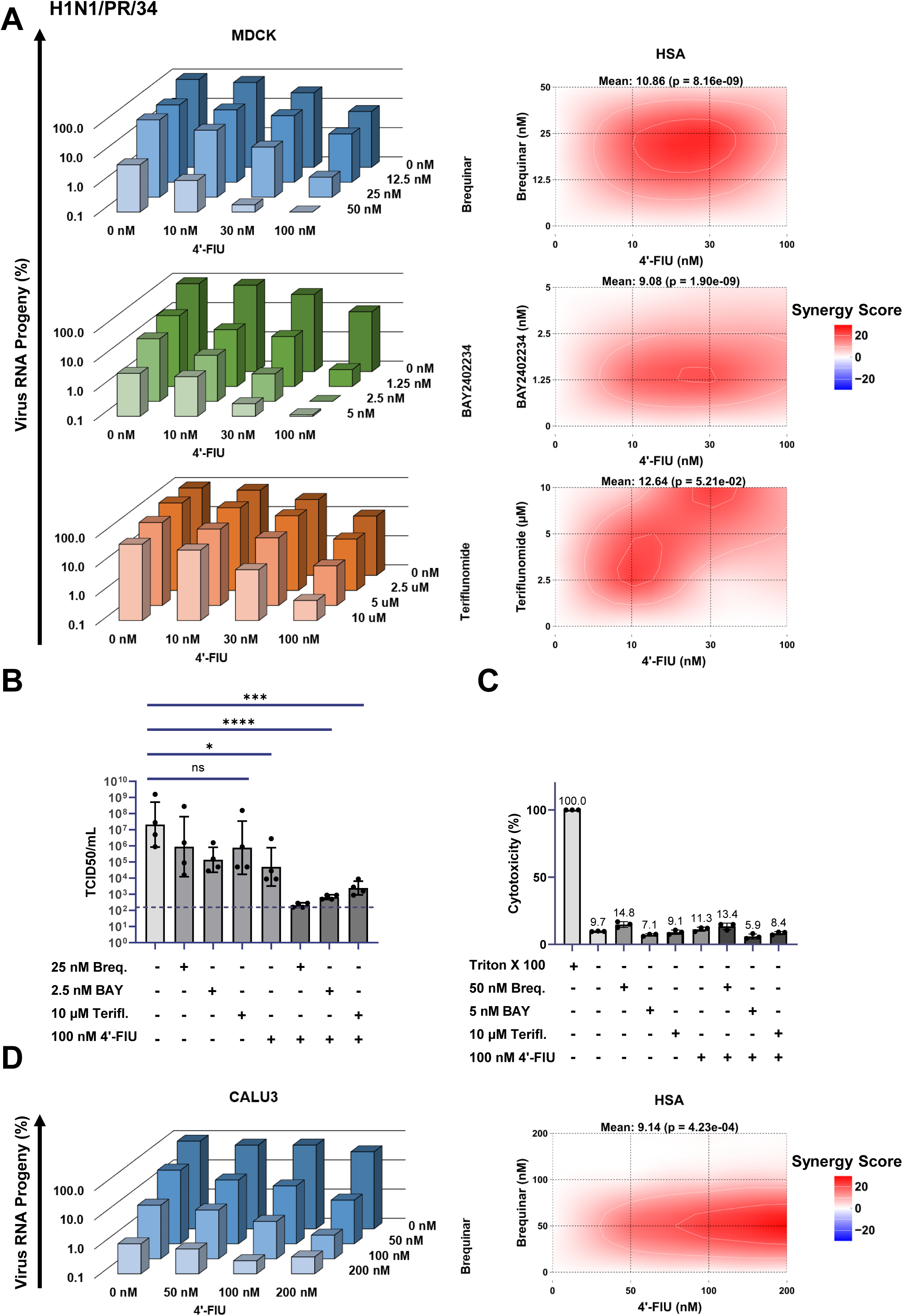
Synergistic effect of 4′-FlU and DHODH inhibitors on influenza A virus replication. (A) Synergistic antiviral activity of 4′-FlU combined with different DHODH inhibitors, observed on IAV (H1N1/PR/34) replication. MDCK cells were infected and treated 1 h post-infection with varying concentrations of 4′-FlU, alone or in combination with the DHODH inhibitors brequinar, BAY2402234, and teriflunomide. After 48 h, viral RNA was extracted from the supernatant and quantified using qRT-PCR. Undisturbed virus growth was set to 100%. Synergy scores were calculated using Synergyfinder.org utilizing the HSA model. Each data point represents the mean of 3 biological replicates. A mean p-value < 0.05 indicates a significant global synergistic effect. Synergy score values > 10 indicate synergism while < -10 indicates an antagonistic effect; scores between -10 and 10 reflect an additive effect of the drugs. Cf. Figures S1, S2. (B) The combination of 4′-FlU and DHODH inhibitors significantly reduces viral titers. Virus infection and treatment were performed as in (A). After 48 h, the supernatant containing the virus was harvested and titrated on MDCK cells to determine the TCID50/mL (mean with SD, n=3). (C) Evaluation of cytotoxicity induced by single drugs or combination. Uninfected MDCK cells were cultured for 48 h in the presence of 4′-FlU alone or in combination with DHODH inhibitors. Then, the supernatant was harvested to quantify the amount of lactate dehydrogenase (LDH) released from the cells for each condition, serving as an indicator of cytotoxicity. Total LDH release was measured by lysing the cells with Triton X100 (mean with SD, n=3). (D) Comparable synergy is observed in Calu-3 cells. The proposed drug combination also exhibits strong reduction of viral replication in a human lung cancer cell line (mean, n=3). Cf. Figure S3.

### DHODH inhibition re-sensitizes 4′-FlU-resistant IAV to 4′-FlU

IAV can develop resistance towards 4′-FlU upon prolonged exposure to the drug, especially in suboptimal concentrations, mostly due to mutations within the viral RNA-dependent RNA polymerase (RdRP) complex. The most prominent resistance-conferring mutation was identified as the exchange Y488C within the polymerase basic protein 2 (PB2) (Lieber et al., 2024). We recapitulated this in an in vitro selection scheme and found the same mutation quickly arising (Figures 2A, S4). Strikingly, the combination of 4′-FlU and DHODH inhibitors continued to suppress the replication of this mutant virus, even at concentrations where the single drugs had no detectable effect on virus replication (Figures 2B, S4). Hence, DHODH inhibition might be able to re-sensitize 4′-FlU-induced escape-mutants or even prevent their occurrence during treatment.

**Figure 2.**
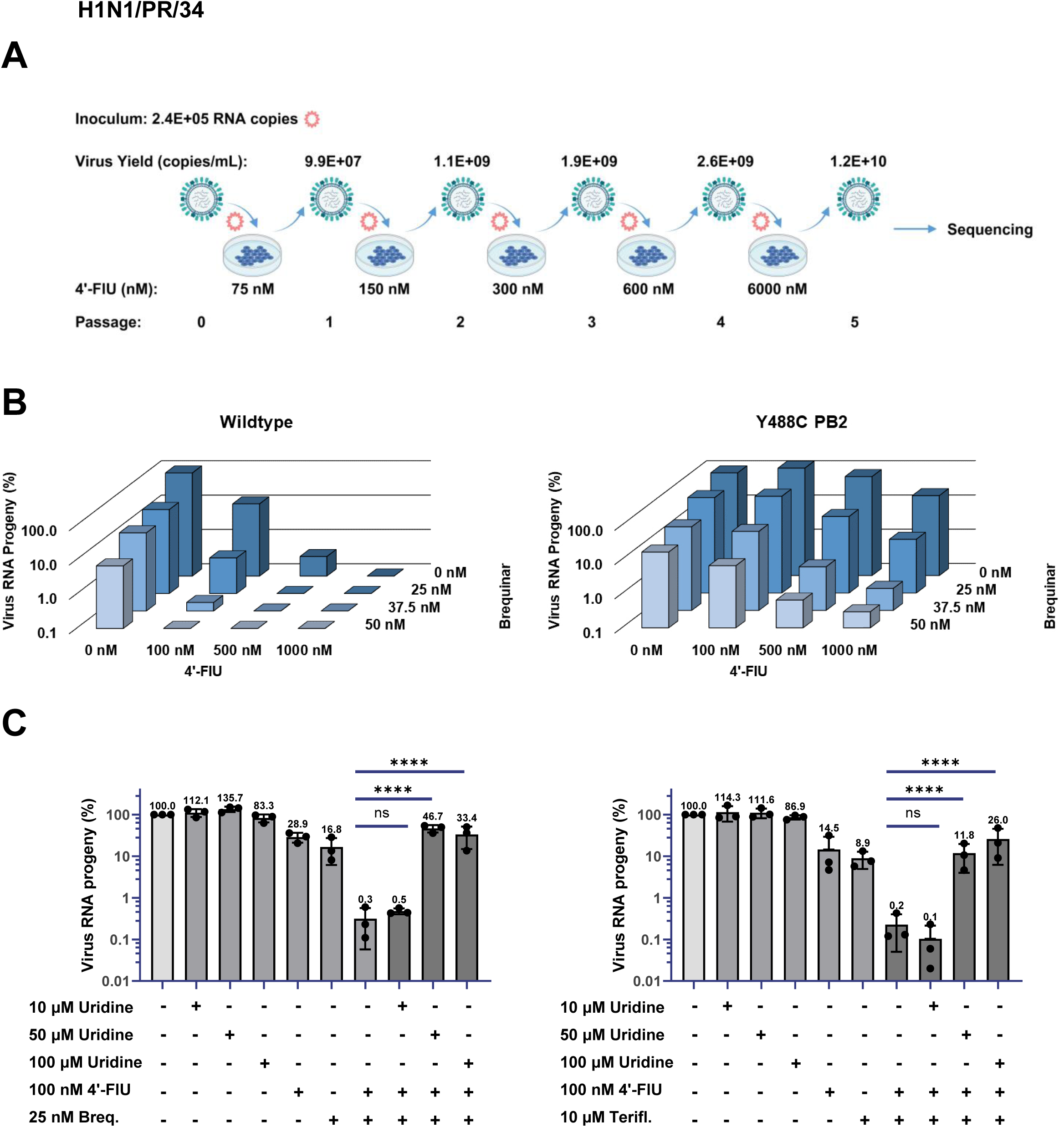
DHODH inhibition rescues 4′-FlU efficacy to interfere with the replication of resistant influenza A viruses. (A) Selection of 4′-FlU resistant IAV. IAV (H1N1/PR/34) was passaged five times in the presence or absence of increasing levels of 4′-FlU on MDCK cells. For each passage, viral supernatant was collected 48 h post-infection, with each subsequent infection maintaining a controlled inoculum concentration. Selection with the drug resulted in the mutation Y488C in the PB2 subunit of the IAV RNA polymerase, cf. Figure S4. (B) 4′-FlU efficacy against resistant IAV can be rescued by simultaneous DHODH inhibition. MDCK cells were infected with either Wildtype (WT) or Y488C PB2 H1N1/PR/34 IAV and subsequently treated with increasing concentrations of 4′-FlU and/or brequinar. Viral RNA was harvested from the supernatant 32 h post infection and quantified using qRT-PCR (mean, n=3). Cf. Figure S4. (C) The synergistic effect of 4′-FlU and pyrimidine depletion is reversed upon the addition of uridine. MDCK cells were infected and treated as in (A), with or without increasing concentrations of uridine. Viral RNA from the supernatant was harvested and quantified as in (B) (mean with SD, n=3).

To clarify the mechanism by which DHODH inhibitors fortify the antiviral effect of 4′-FlU, we treated the infected cells with 4′-FlU in combination with either brequinar or teriflunomide. The addition of uridine to the media largely restored viral replication in a dose-dependent manner (Figure 2C). This strongly suggests that the depletion of uridine is central to the observed drug synergy. We hypothesize that upon inhibition of DHODH, endogenous uridine synthesis drops, thereby enhancing the incorporation of 4′-FlU into nascent viral RNAs.

### 4′-FlU and DHODH inhibitors synergize to inhibit various IAV subtypes

To assess how broadly 4′-FlU and DHODH inhibitors interfere with IAV replication, we treated cells that were infected with distinct IAV strains, i.e. the H3N2 virus strain A/Panama/2007/1999 (H3N2/PN/99) and the H1N1 strain A/Wilson-Smith/1933 (H1N1/WSN/33). In all cases, the drug combinations profoundly diminished the extent of viral replication, far more than the single drugs, as determined by qRT-PCR (Figures 3A, S5). In addition, the drug combination led to a considerable reduction of the infection rates, as shown by immunofluorescence analysis using antibodies directed against the IAV nucleoprotein (NP) (Figure 3B). Likewise, the levels of IAV NP were strongly diminished by the drug combination, whereas single drugs had a much lesser effect (Figure 3C). This indicates that the synergistic efficacy of 4′-FlU and inhibitors of pyrimidine synthesis against IAV is applicable to diverse IAV strains.

**Figure 3.**
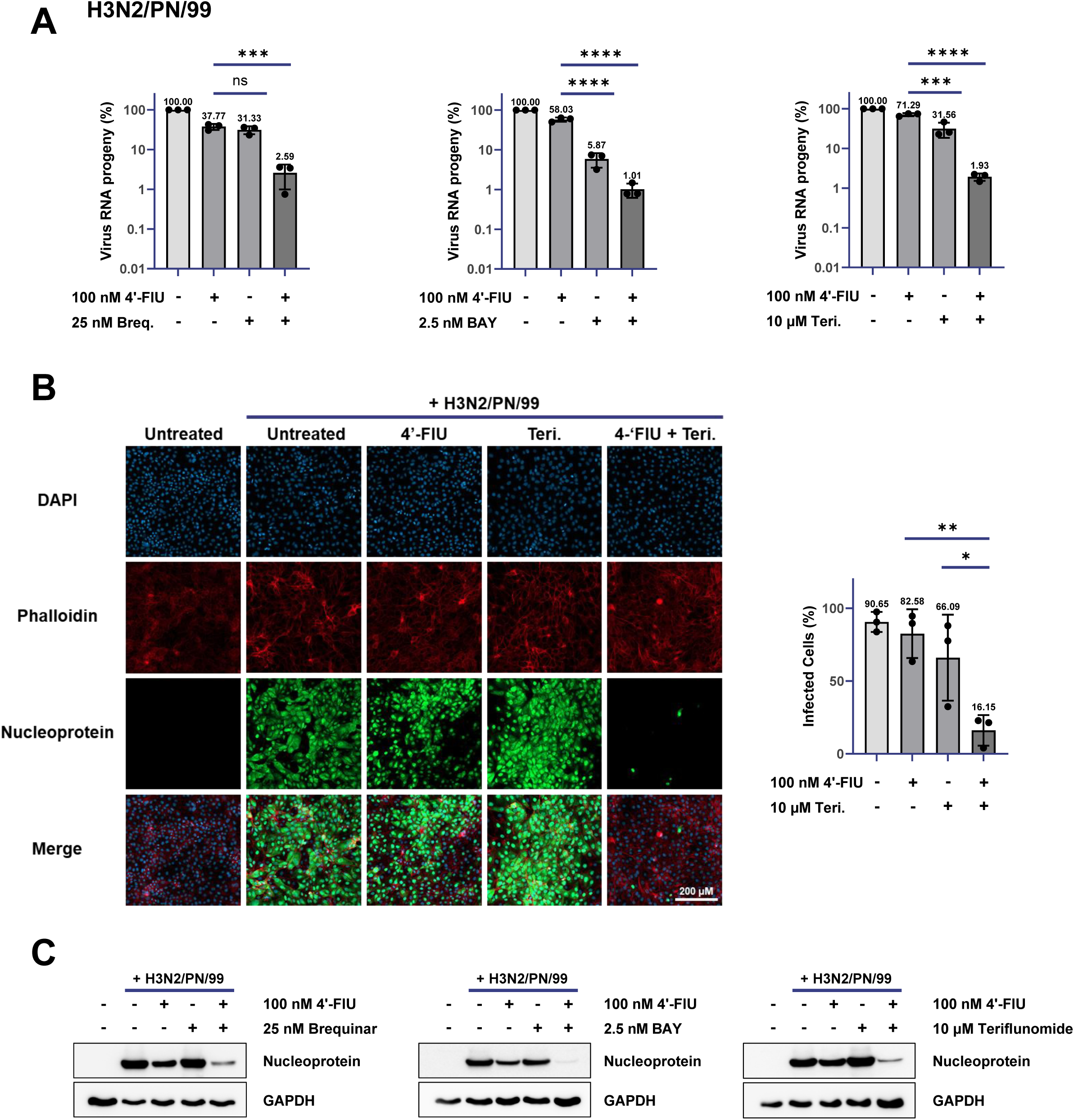
Effects of 4′-FlU and DHODH inhibitor combination on H3N2/PN/99 influenza A virus proliferation. (A) Synergistic reduction of IAV H3N2/PN/99 replication by combining 4′-FlU and DHODH inhibitors. Infected MDCK cells were treated with 4′-FlU and the indicated DHODH inhibitors after 1 h. Viral RNA was harvested 24 h post-infection from the supernatant, and viral RNA amounts were quantified by qRT-PCR (mean with SD, n=3). For an analogous experiment using IAV H1N1/WSN/33, cf. Figure S5. (B) Strong inhibition of viral infection spread by combining 4′-FlU and DHODH inhibitor. MDCK cells, infected and treated as described in (A), were fixed with 4% paraformaldehyde at 24 h post infection. Virus infection rates were assessed by quantitative immunofluorescence (mean with SD, n=3). Left, representative images. Right, quantification. (C) Suppressed viral protein production by the drug combination. MDCK cells were infected with IAV (H3N2/PN/99) at an MOI of 0.01 in the presence of no drug, 4′-FlU, the indicated DHODH inhibitors, or both 4′-FlU and DHODH inhibitor. Protein lysates from infected and treated MDCK cells were collected 24 h post infection and analyzed by Western blot using antibodies against IAV nucleoprotein, with cellular GAPDH serving as a loading control.

### The synergy of 4′-FlU and DHODH inhibitors is observed on zoonotic influenza A viruses

IAVs from birds or non-human mammals represent a constant pandemic threat, as a result of spill-over events between animals and the human population (Simon et al., 2014) and the possibility of forming new reassortant viruses displaying antigenic shift (Krammer et al., 2018). To assess the efficacy of the drug combination against zoonotic viruses, we tested two porcine and two avian IAV strains. A/swine/Germany/SIR32/2022 (H1avN2) and A/swine/Germany/SIR33/2022 (H1pdmN2) represent viruses circulating in European pig farms (Simon et al., 2014). H1pdmN2 is a closely related descendant of the pandemic virus from the year 2009. H1avN2 is derived from birds and became endemic in pig populations years ago (Henritzi et al., 2020). Sporadic transmissions of these viruses to humans were observed (Durrwald et al., 2020; Parys et al., 2021), underlining their zoonotic potential. Moreover, two highly pathogenic avian IAVs, A/chicken/Germany-NI/AI04286/2022 (H5N1) and A/chicken/Germany-NW/AI03705/2021 (H5N8), were tested regarding their drug susceptibilities. H5N1 viruses are currently spreading globally, infecting a large variety of mammalian hosts (Burrough et al., 2024). Since 2016, H5N8 viruses have caused devastating outbreaks in domestic poultry, and sporadic infections of humans have been reported (Calle-Hernandez et al., 2023) (https://www.ecdc.europa.eu/en/publications-data/threat-assessment-first-human-cases-avian-influenza-h5n8). As shown in Figures 4, S6 and S7, these virus strains were also highly susceptible to the synergy of 4′-FlU and the DHODH inhibitor teriflunomide, as revealed by qRT-PCR analysis of the virus-containing cell supernatants. Thus, the drug combination might potentially help to confine outbreaks caused by reassortant IAVs.

**Figure 4.**
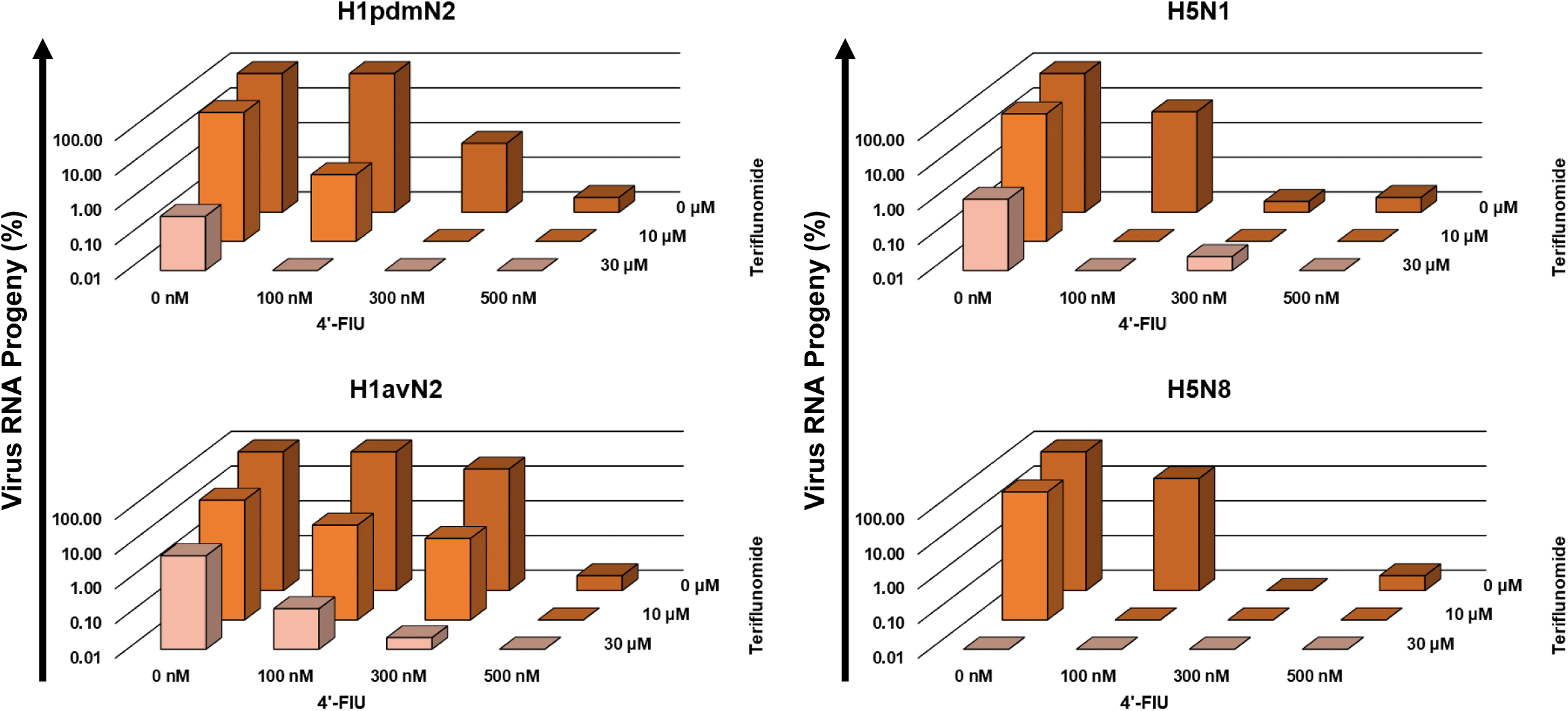
Broadly effective synergism of 4′-FlU and DHODH inhibitors against recent IAV strains. MDCK cells were infected with a selection of recent IAV strains isolated from deceased animals (birds and pigs). A/swine/Germany/SIR32/2022 (H1avN2) and A/swine/Germany/SIR33/2022 (H1pdmN2) are porcine viruses (Simon et al., 2014). H1pdmN2 is a closely related descendant of the pandemic virus from the year 2009. H1avN2 is derived from birds and became endemic in pig populations years ago (Henritzi et al., 2020). Moreover, two highly pathogenic avian IAVs, A/chicken/Germany-NI/AI04286/2022 (H5N1) and A/chicken/Germany-NW/AI03705/2021 (H5N8), were investigated regarding their drug susceptibilities. Following infection with MOI 0.01, cells were treated with the indicated drug concentrations alone or in combination and incubated for 48 h at 37°C. Supernatant was harvested and virus RNA progeny was determined using qRT-PCR targeting the M1 gene of IAVs. All values were normalized to the untreated control, which was set to 100%. While exhibiting slight variations in overall response, all four tested IAV strains displayed a notably enhanced inhibition of replication upon administration of the drug combination. For standard deviations and analyses of significance, cf. Figures S6, S7.

### Combining 4′-FlU with inhibitors of DHODH synergistically diminishes the replication of SARS-CoV-2

Next, we asked whether the previously reported antiviral effect of 4′-FlU on SARS-CoV-2 (Sourimant et al., 2022) could also be enhanced by inhibiting DHODH. This was plausible since we (Stegmann et al., 2022) and others (Schultz et al., 2022) have previously observed strong synergy of DHODH inhibitors with another pyrimidine analogue, NHC, to antagonize SARS-CoV-2 replication. We combined 4′-FlU with the DHODH inhibitors BAY 2402234 and brequinar on cells infected with a Wuhan-like SARS-CoV-2 strain that we had previously isolated (Stegmann et al., 2021). As a result, we found strongly diminished cytopathic effect (CPE) when combining the drugs (Figure 5A) without detectable cytotoxicity of the drugs themselves (Figure 5B). Production of viral RNA was synergistically reduced (Figures 5C, D, S8, S9), and diminished virus propagation was also found for the SARS-CoV-2 Omicron variant B.1.1.529 (Figures 5E, S10). Similar synergies were observed when quantifying infectious particles released to the supernatant (Figure 5F), assessing infection rates (Figures 5G, H, S11), or detecting viral proteins (Figure 5I). In conclusion, the combination of 4′-FlU with a DHODH inhibitor strongly and synergistically compromises the replication of SARS-CoV-2.

**Figure 5.**
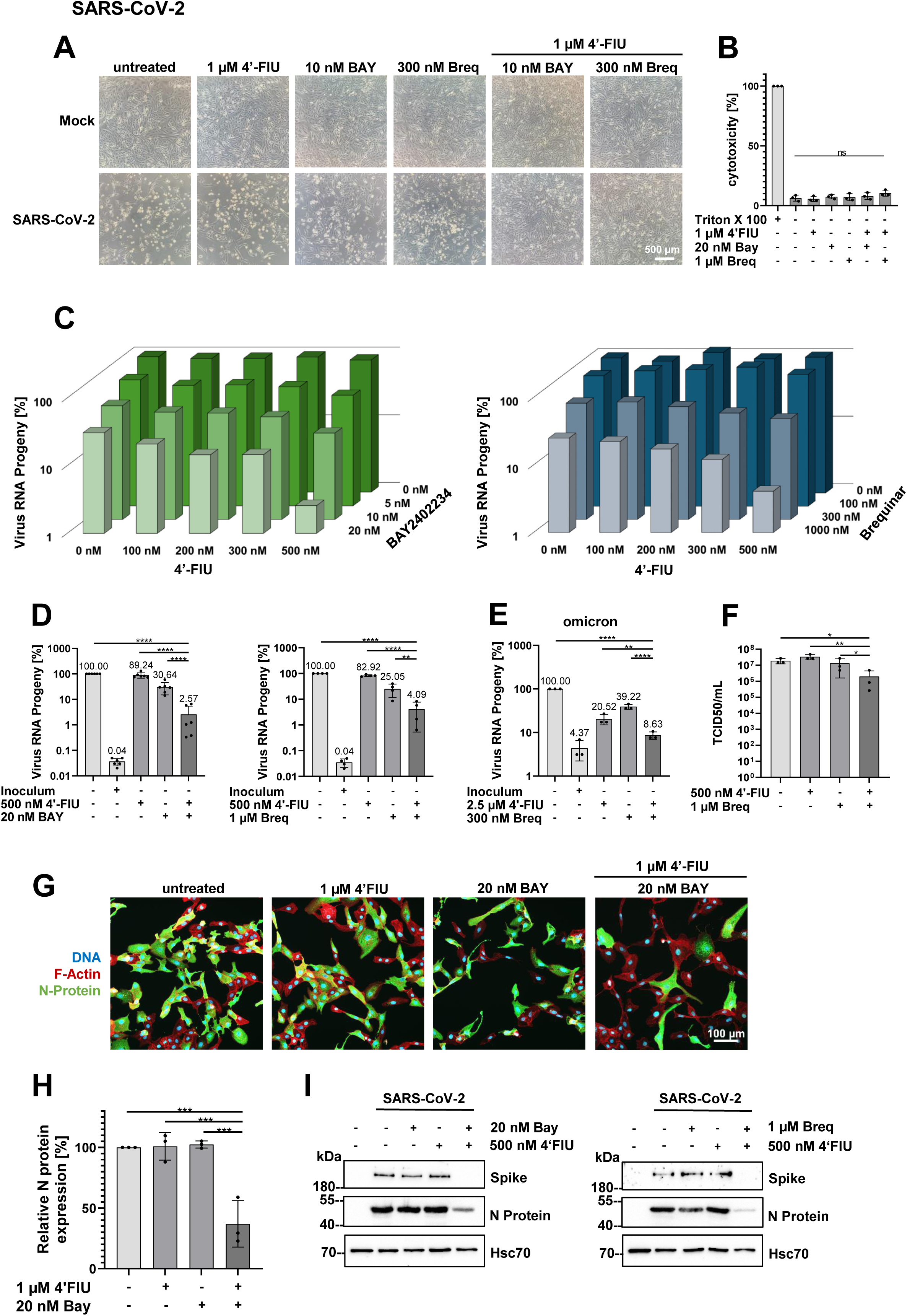
Synergistic effect of 4′-FlU and DHODH inhibitors on SARS-CoV-2 replication. (A) Combination treatment with 4′-FlU and the DHODH inhibitors BAY2402234 or brequinar reduces the cytopathic effect (CPE) caused by SARS-CoV-2. Vero E6 cells were infected with SARS-CoV-2, Wuhan-like Gottingen isolate (Stegmann et al., 2021) (MOI 0.1) or left uninfected (Mock). 1 h post infection (hpi), the cells were treated with the indicated concentrations of the drugs and incubated for 48 h. Cell morphology was observed by phase contrast microscopy (scale bar, 500 µm). (B) Combination treatment by 4′-FlU and DHODH inhibitors does not cause cytotoxic effect in Vero E6 cells. The cells were treated with 4′-FlU and DHODH inhibitors for 48 h, and the cell supernatants were tested for lactate dehydrogenase (LDH) release as an indicator of cell damage. The percentage of LDH release was calculated compared to total LDH release of Triton X 100-treated cells (mean with SD; n = 3). (C) 4′-FlU and the DHODH inhibitors BAY2402234 and brequinar synergistically reduce the amount of SARS-CoV-2 RNA in the supernatants of infected cells. Vero E6 cells were infected as in (A). The RNA was isolated from cell supernatants and quantified by qRT-PCR. The amounts of viral RNA are shown in percent of untreated control (mean; n = 6 for 4′-FlU and BAY2402234; n = 4 for 4′-FlU and brequinar; logarithmic scale). Cf. Figures S8, S9 for standard deviations and significances. (D) Selected drug concentrations analyzed in (C) displayed with all data points and standard deviations. The inoculum sample was isolated immediately after infection and indicates the background signal obtained without virus replication. The untreated control was set to 100% virus RNA progeny and the other values were normalized to it. In comparison to single treatment, the combination treatment reduced the virus RNA progeny with high significance (mean with SD; n = 6 / n = 4 respectively; logarithmic scale). (E) 4′-FlU and the DHODH inhibitor brequinar reduce the RNA progeny of SARS-CoV-2 Omicron variant. Vero E6 cells were infected with SARS-CoV-2 variant omicron (MOI 0.03), treated with the indicated drugs at 1 hpi and incubated for 48 h. Viral RNA in the supernatant was quantified by qRT-PCR. 100% virus RNA progeny was assigned to the untreated control, and the RNA levels obtained upon treatment were normalized accordingly (mean with SD; n = 3; logarithmic scale). Cf. Figure S10. (F) The combination of 4′-FlU and the DHODH inhibitor brequinar reduces SARS-CoV-2 titers. After infection and treatment of Vero E6 cells as in (A), virus-containing supernatant was titrated by tissue culture infectious dose (TCID50) assay (mean with SD; n = 3; logarithmic scale). (G) 4′-FlU and the DHODH inhibitor BAY2402234 in combination reduce SARS-CoV-2 infection rates. Vero E6 cells, seeded on slides, were infected and treated as in (A). At 48 hpi, the cells were fixed with 4% PFA for 1 h and stained with an antibody directed against the viral nucleocapsid protein (N protein) to visualize viral infection. 4,6-Diamidino-2-phenylindole (DAPI) and phalloidin were used to stain cellular DNA and F-actin, respectively. Images were taken with Zeiss Celldiscoverer 7 automated fluorescence microscope (scale bar, 100 µm). For separate color channels, cf. Figure S11. (H) The percentage of SARS-CoV-2-infected cells was calculated from whole well scans as in (G), using the Gene- and Protein Expression tool provided by Zen, Zeiss (mean with SD; n = 3). (I) Spike and N protein levels are reduced in SARS-CoV-2-infected cells following treatment with 4′-FlU and a DHODH inhibitor. Vero E6 cells were infected and treated as described for (A), and protein lysates were used for Western Blot analysis. Spike and N proteins were stained with respective antibodies, and the cellular HSC70 protein was detected as loading control.

### 4′-FlU diminishes SARS-CoV-2 replication in a hamster model

To assess the antiviral efficacy of 4′-FlU and DHODH inhibition in animals, hamsters were infected with SARS-CoV-2 and treated either with 4′-FlU and/or teriflunomide, or vehicle as untreated control (Figures 6A). We measured oral and nasal virus shedding in the animals, quantifying both viral RNA and infectious particles in oral swabs and nasal lavage samples (Figure S12A, B). Treatment with teriflunomide alone resulted in a minor change in virus shedding. In contrast, treatment with 4′-FIU, alone or in combination with teriflunomide, strongly diminished and mostly abolished the shedding of replication competent virus at all measured time points.

**Figure 6:**
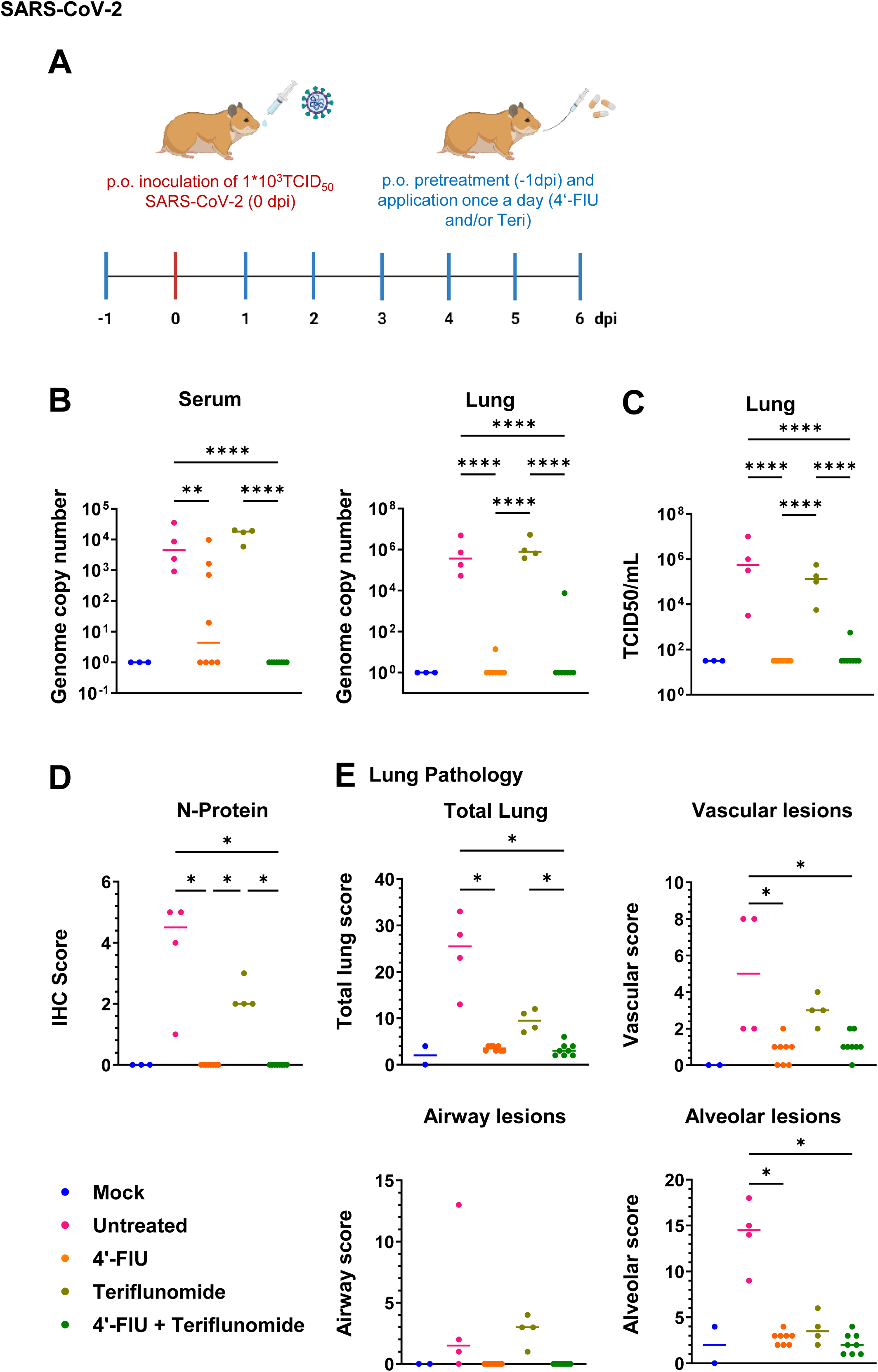
Drug efficacy in a hamster model of SARS-CoV-2 infection. (A) To test the efficacy of 4′-FlU and the DHODH inhibitor teriflunomide in vivo, hamsters were treated and/or infected with SARS-CoV-2 as outlined in the scheme. Virus RNA and infectious particles were measured in oral swabs and nasal lavage samples, cf. Figure S12A, B. (B) At the end of the experiment, postmortem analyses of virus RNA in blood (serum) and lungs were performed, by qRT-PCR. Teriflunomide alone did not significantly diminish virus load, whereas 4′-FlU as well as its combination with teriflunomide were highly efficient to eliminate detectable virus. Cf. Figure S12C for virus RNA in nasal conchae and tracheae. (C) Postmortem analysis of infectious particles in the lungs by measurement of TCID50. The results correspond to the RNA levels determined in (B). Cf. Figure S12C for infectious particles in nasal conchae and tracheae. (D) Virus-infected cells were detected by immunohistochemistry (IHC) staining of lung tissue post mortem. Here again, 4′-FlU, alone or in combination, efficiently eliminated detectable virus. Examples of the IHC results are shown in Figure S13. (E) Pathomorphological analyses of the lung tissues obtained from infected and/or treated animals. Scores were calculated as previously described (Aksu et al., 2024; Armando et al., 2022) to reflect the COVID-like pathological alterations in total lungs, blood vessels (vascular), airways, and alveoli. Most alterations were largely suppressed by 4′-FlU, alone or together with teriflunomide. Scatter plots of Kruskal-Wallis multi-comparison test with subsequent pairwise Mann-Whitney-U-tests and Benjamini-Hochberg correction for multiple comparisons (False Detection Rate: 0.05, asterisk: discovery, q<0.05). Representative images are shown in Figures S14 and S15.

At 6 days post infection (dpi), we detected viral genomic RNA in the sera, the lungs, the nasal conchae and the tracheae of infected but otherwise untreated animals (Figures 6B, S12C), with teriflunomide treatment alone making little difference in this regard. In contrast, treatment with 4′-FlU eliminated most detectable viral RNA. This therapeutic effect was so strong that it was hardly enhanced by the addition of teriflunomide. Similar observations were made when analyzing the amount of infectious virus in the lungs, nasal conchae and tracheae of the animals (Figures 6C, S12C). Moreover, we detected the SARS-CoV-2 N protein in the lungs of the animals by immunohistochemistry (Figures 6D, S13). Moderate to strong labeling of the SARS-CoV-2 nucleoprotein was present in the untreated and infected group (median score 4.5; range 1 – 5) and to a somewhat lesser extent in the teriflunomide-treated group (median score: 2; range 2 - 3), while there was no immunolabeling in 4′-FlU-treated and combination-treated groups (total lung IHC score: 0). Thus, the treatment with 4′-FlU, with or without additional teriflunomide, strongly suppressed the propagation of SARS-CoV-2 in hamsters.

The untreated, SARS-CoV-2-infected hamsters developed pathomorphological changes that recapitulate COVID-19, consisting of alveolar and airway inflammation accompanied by prominent vascular alterations (Figures 6E, S14, S15). The median total lung score in this group was 25.5 (range 13 - 33). Teriflunomide alone reduced this score to about 9.5 (range 7 - 12); here, signs of vasculitis remained but alveolar lesions were decreased. When treated with 4′-FlU, or with the combination of teriflunomide and 4′-FlU, only minimal pathological changes remained within all examined animals. The remaining alterations were characterized by alveolar and perivascular edema and hemorrhage, most likely related to euthanasia. The median total lung score of the 4′-FlU group was 3.5 (range 3 - 4). In the combined teriflunomide/4′-FlU group, the median total lung score was 3 (range 2 - 4).

In summary, no COVID-related lesions were detected in both treatment groups that included 4′-FlU; here, the levels of replication competent virus in the respiratory tract tissues were significantly reduced, virus shedding was completely abolished, and pathomorphological alterations were minimal. Due to the strong effects already seen in the 4′-FlU-treated animals, a synergistic effect of the combined therapy with 4′-FlU and teriflunomide was not detectable. However, the combination treatment was at least as effective as 4′-FlU monotherapy.

### 4′-FlU inhibits Cedar and Nipah virus replication, and DHODH inhibitors synergistically enhance its efficacy

Henipaviruses cause infections with some of the highest case fatality rate in humans. As a model system for the genus *Henipavirus*, we employed recombinant Cedar virus (CedV). CedV represents a low pathogenic member of the genus, enabling investigation under BSL2 conditions and serving as a model for the highly virulent NiV and Hendra virus (Amaya et al., 2021; Laing et al., 2018; Marsh et al., 2012).

To investigate how a henipavirus responds to 4′-FlU, and DHODH inhibition, CedV was propagated in the presence of the drugs. 4′-FlU potently blocked the replication of CedV, and this antiviral effect was strongly fortified by the addition of the DHODH inhibitor teriflunomide (Figure 7A, B). Next, we analyzed the effect on the highly pathogenic NiV. Indeed, similar patterns of drug efficiency could be observed, moderately reducing virus replication after single treatment with 4′-FlU, and strongly in combination with teriflunomide (Figure 7C, D). Thus, henipaviruses are susceptible to treatment with 4′-FlU, and DHODH inhibition further boosts the antiviral activity of 4′-FlU.

**Figure 7.**
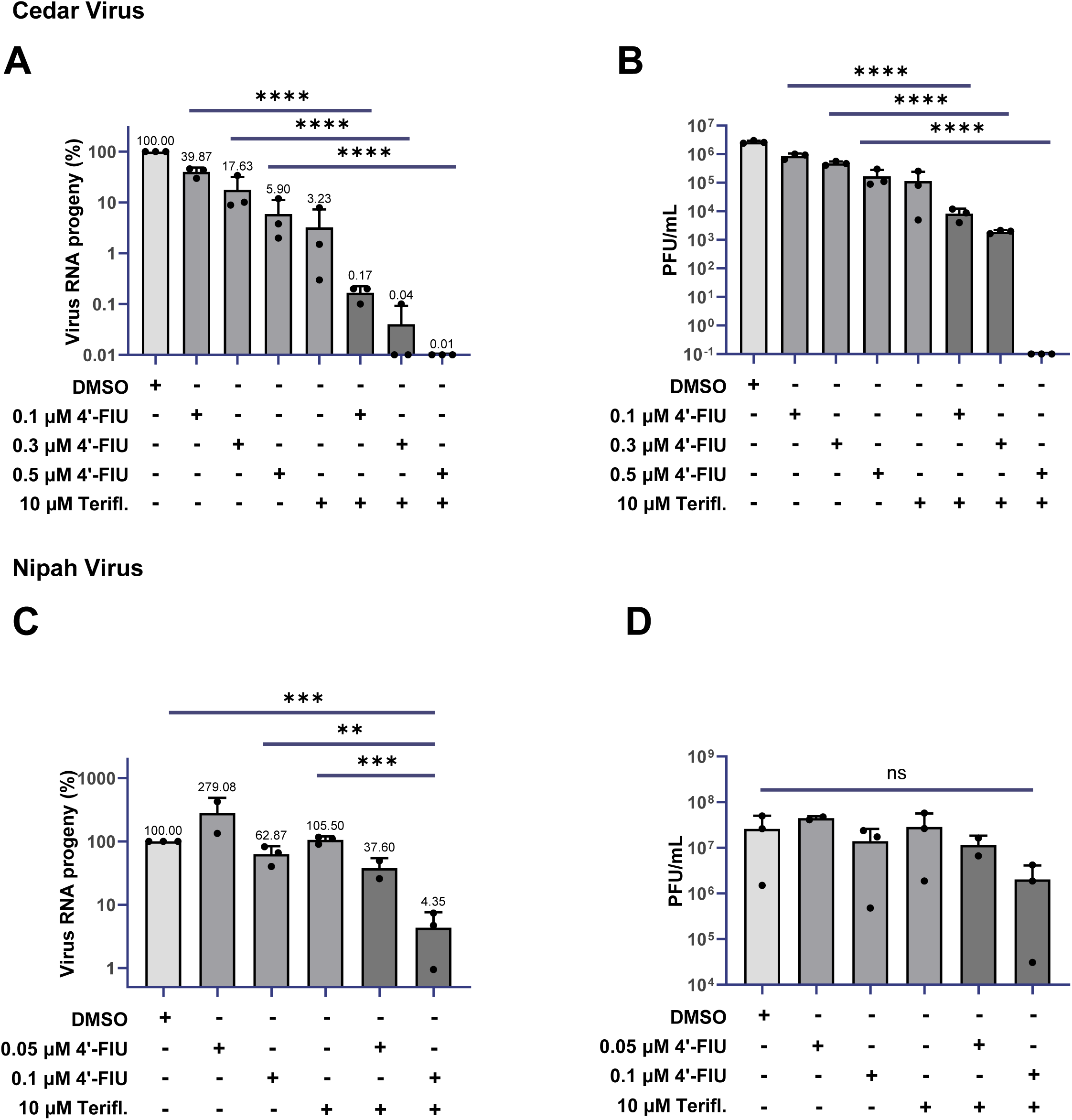
Diminished replication of Cedar and Nipah virus in the presence of 4′-FlU and DHODH inhibitors. (A) Cedar virus (MOI 0.001) was used to infect Vero76 cells for 1 h before treatment with 4′-FlU and the DHODH inhibitor teriflunomide (Teri) at the indicated concentrations. The viral RNA contained in the supernatant was quantified by qRT-PCR. Virus RNA progeny is plotted in percent and all values were normalized to the untreated control which was set to 100% (mean; n = 3; logarithmic scale). (B) Titration of infectious virus in the supernatants of infected cells revealed a reduction of plaque forming units (PFU) following treatment as in (A). (C) Vero76 cells were infected with Nipah virus (MOI=0.02) for 1 h and further incubated for 24 h while being treated with 4′-FlU and/ or teriflunomide at the indicated concentrations. The supernatant was harvested, and virus RNA progeny was determined via qRT-PCR targeting the N gene. Virus RNA progeny, normalized to the untreated control which was set to 100% (mean; n = 3; logarithmic scale). (D) After performing the experiment described in (C), Nipah virus was titrated in duplicates using plaque assays.

### 4′-FlU and DHODH inhibitors synergize to counteract Ebola virus infection

Finally, we tested the antiviral activity of 4′-FlU and the effect of combination treatment on the highly pathogenic EBOV. Like NiV, EBOV belongs to the group of nonsegmented negative sense RNA viruses and causes outbreaks with high case fatality rates. The currently approved therapeutic options against EBOV are restricted to antibody-based therapies, whereas there is a lack of licensed small molecule compounds (Almeida-Pinto et al., 2024). We therefore addressed two questions: a) whether 4′-FlU shows antiviral activity against EBOV and b) whether 4′-FlU-mediated inhibition could be fortified by DHODH inhibitors. We used recombinant EBOV expressing ZsGreen (EBOV-ZsGreen) to facilitate the readout of infection under BSL4 conditions (Hume et al., 2022). We observed potent activity of 4′-FlU against EBOV in Huh7 (human liver carcinoma) cells, as demonstrated by a dose-dependent decrease of EBOV-ZsGreen infection rates upon 4′-FlU treatment (Figures 8A and B, S16 and S17). Moreover, when combining 4′-FlU with the DHODH inhibitor brequinar, we noted a synergistic reduction of EBOV-ZsGreen replication, with synergy scores of greater than 20 when using 30 or 100 nM brequinar, despite mild inhibition by brequinar alone (Figures 8B, C, S17). This strongly suggests that 4′-FlU might be a promising therapeutic candidate for the treatment of EBOV disease and that DHODH inhibitors might further amplify the therapeutic effect.

**Figure 8.**
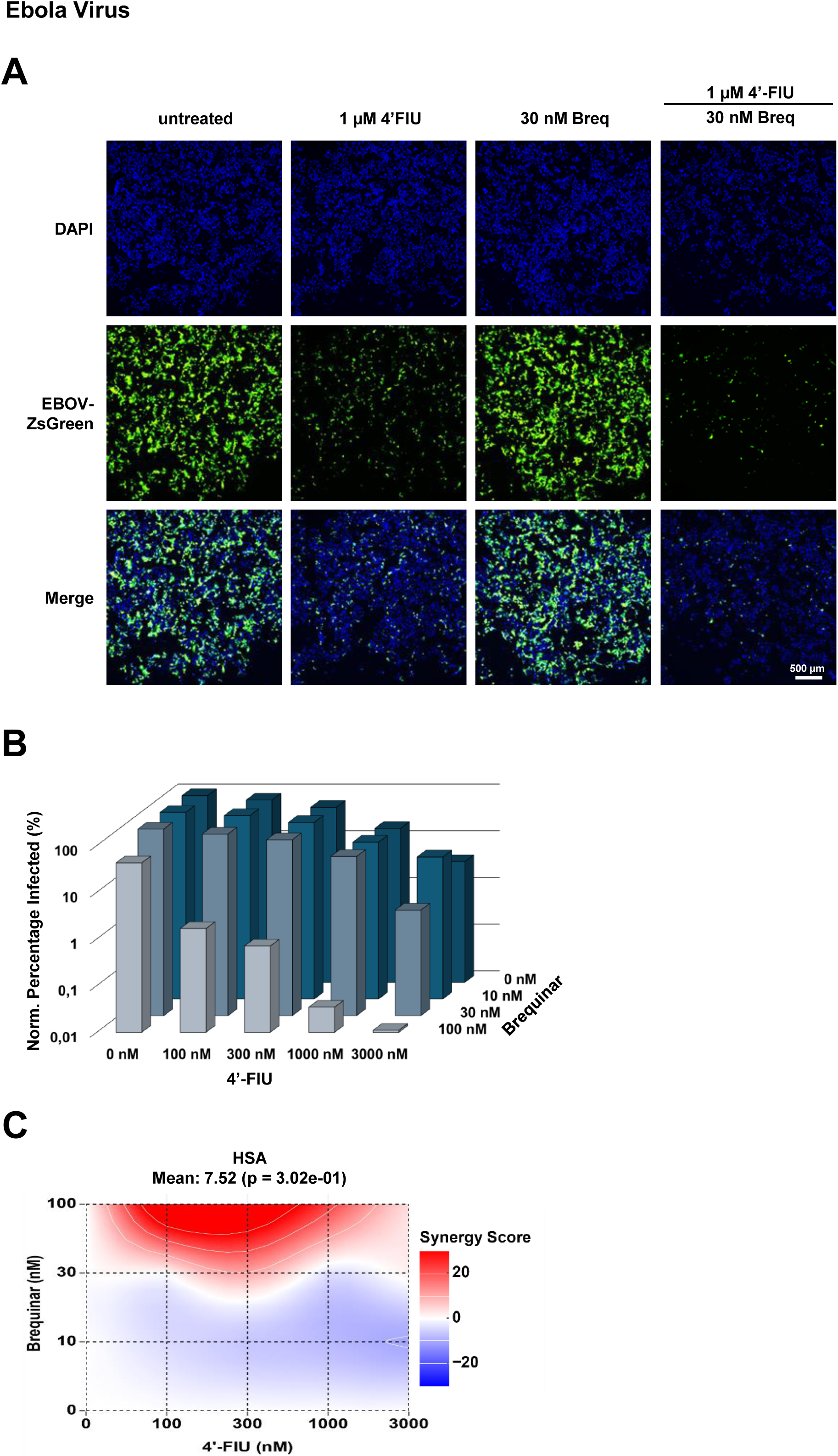
Combination treatment of 4′-FlU and brequinar inhibits Ebola virus replication. (A) Combination treatment with 4′-FlU and the DHODH inhibitor brequinar reduced EBOV replication synergistically. Huh7 cells were pretreated for 1 h with the indicated drug concentrations before infection with EBOV-ZsGreen (MOI 0.1). 24 hpi, 4′-FlU was replenished in the respective wells. At 48 hpi, the cells were fixed with 10% formalin and cellular DNA was stained with DAPI. Scale bar, 500 μm. Cf. Figure S16 for additional images upon treatment with different drug concentrations. (B) The percentage of infected cells in (A) was determined using the QuPath software. Values were normalized to the EBOV-ZsGreen-infected, untreated control which was set to 100% (mean; n = 3; logarithmic scale). Cf. Figure S17 for single data points, standard deviations and significances. (C) Synergy of 4′-FlU and brequinar treatment. Synergy scores were calculated as described in the legend to figure 1. Synergy score values >10 indicate synergism. .

## DISCUSSION

Our results indicate that the inhibition of pyrimidine synthesis synergistically cooperates with 4′-FlU to effectively counteract the replication of a range of RNA viruses, including positive sense RNA viruses (SARS-CoV-2), segmented negative sense RNA viruses (IAV), and nonsegmented negative sense RNA viruses (CedV, NiV, EBOV). Mechanistically, we propose that DHODH inhibition depletes endogenous uridine, thereby increasing the incorporation of 4′-FlU into nascent viral RNA. Previous work demonstrated antiviral activity of 4′-FlU against influenza viruses, including IAV, influenza B virus, and SARS-CoV-2 (Lieber et al., 2023; Lieber and Plemper, 2022; Sourimant et al., 2022). These viruses are also susceptible to DHODH inhibition (Hahn et al., 2020; Sibille et al., 2022; Stegmann et al., 2022; Zheng et al., 2022). However, the synergistic effect of the drug combination has not been shown before. A moderate response of henipaviruses to 4′-FlU was previously observed (Sourimant et al., 2022), but here again, we newly describe the addition of DHODH inhibitors to improve this antiviral activity against CedV and NiV (Figure 7). In the case of EBOV, both the efficacy of 4′-FlU and its synergy with DHODH inhibitors are reported for the first time (Figure 8, S16 and S17). These results further indicate the broadly applicable potential of 4′-FlU as an antiviral drug, and the increase of its potency by interfering with cellular pyrimidine synthesis.

4′-FlU treatment can give rise to the selection of resistant IAV mutants, even though these mutants appeared attenuated in an animal model (Lieber et al., 2024). The addition of DHODH inhibitors partially re-sensitized the mutant virus to 4′-FlU treatment (Figures 2B, S4), raising the perspective that DHODH inhibition might prevent the replication of 4′-FlU-resistant mutants in the case of other viruses, too. Moreover, since DHODH inhibitors target a cellular metabolic pathway, viruses are less likely to develop specific resistance mechanisms towards these drugs. Taken together, the combination of 4′-FlU and DHODH inhibitors holds the potential for broad efficacy against a number of pathogenic viruses with a low potential for the development of resistance.

The inhibition of DHODH can dramatically fortify the efficacy of an antiviral pyrimidine analogue, as shown here for 4′-FlU and previously for N4-hydroxycytidine/molnupiravir (Schultz et al., 2022; Stegmann et al., 2022). Other pyrimidine analogues that might be improved by similar combinations include lumicitabine, for treating infections with respiratory syncytial virus (Patel et al., 2019) but also NiV (Lo et al., 2020); sofosbuvir for hepatitis C virus (Lawitz et al., 2015), brivudine for varizella zoster virus (De Clercq, 2023), and zidovudine for HIV (Yarchoan et al., 1986), among others (Geraghty et al., 2021). In general, multiple opportunities exist to achieve synergy between inhibitors of nucleotide biosynthesis and antiviral nucleoside analogues (de Mariz, 2023).

In addition to the viruses investigated here, 4′-FlU is also active against respiratory syncytial virus (RSV), measles virus, human parainfluenza virus type 3, and rabies virus (Sourimant et al., 2022), as well as various other highly pathogenic avian influenza viruses (Lieber et al., 2023). Both 4′-FlU and DHODH inhibitors are orally available, increasing their convenience and applicability in the clinics. Combining 4′-FlU with a DHODH inhibitor might thus be effective against a broad range of RNA viruses and applicable in clinical settings, for quick and sustainable suppression of virus replication.

### Limitations of the study and its implications

The impact of 4’-FlU in the hamster model of COVID-19 (Figures 6, S12, S13, S14, S15) was unexpectedly strong, making it difficult to observe additional improvements by DHODH inhibition. Trying a spectrum of 4’-FlU doses and/or the use of additional infection models is expected to clarify this in future studies. Another potential limitation of these approaches is the availability of uridine in the blood. The levels of serum uridine are estimated to be around 5 µM (Yamamoto et al., 2011), possibly enough to compete with 4′-FlU despite the inhibition of endogenous pyrimidine synthesis through DHODH inhibitors. Thus, potential clinical applications may require prolonged DHODH inhibition to deplete serum uridine in order to observe a more pronounced synergistic effect with 4′-FlU. Moreover, the effectiveness of this treatment might depend on the particular virus infection being treated. At least in the case of IAV, we observed significant rescue of replication only with 50 µM uridine, which is far beyond physiological levels (Figure 2C). This would argue that IAV infections might be particularly amenable to the combination treatment in vivo.

Another caveat with regard to the potential therapeutic use of DHODH inhibitors during viral infections is their effect on the host immune response. These inhibitors are mostly used as immunosuppressants in clinical settings (Leban and Vitt, 2011), acting at least in part through their interference with the proliferation of T cells. Specifically, DHODH inhibition diminishes oxidative phosphorylation (OXPHOS) and aerobic glycolysis in activated T cells through inhibition of complex III of the respiratory chain (Klotz et al., 2019). This would generally be undesirable in the context of virus infections, since it might delay the development of specific immunity and T-cell-mediated elimination of infected cells. On the other hand, excessive immune and inflammatory responses contribute to the pathogenesis of many virus infections, as exemplified most impressively by COVID-19, which is therefore often treated with glucocorticoids (Amati et al., 2023; Mourad et al., 2023). In such cases, the suppressive effect of DHODH inhibitors might be beneficial. We propose that the combined treatment with 4′-FlU and DHODH inhibitor should be administered long enough to ensure complete virus elimination and avoid rebound effects. In this way, the treatment would reduce both virus particle formation and inflammation-related pathogenesis.

## ACKNOWLEDGEMENTS

We thank Stefan Pohlmann (German Primate Center Gottingen, Germany) and Timm Harder (Friedrich-Loeffler-Institut, Greifswald, Germany) for influenza virus strains and expert advice, Karen Linnemannstons for support in advanced microscopy and Carolin Rudiger for support with the Nipah virus infections.

This work was supported by the Volkswagenstiftung VW-9B785 (M.D.), the Coronavirus Forschungsnetzwerk Niedersachsen (CoFoNi, M.D.), the National Institute of Allergy and Infectious Diseases of the National Institutes of Health grant R01AI133486 (E.M.), and the Emerging Pathogens Initiative of the Howard Hughes Medical Institute grant Agmt 9/16/22 (Lead Investigator Anna Pyle; E.M.).

H.L.F. was supported by the Gottingen Graduate School for Neurosciences, Biophysics, and Molecular Biosciences (GGNB). L.S. was supported by the International Max Planck Research School for Genome Science (IMPRS-GS) and the GGNB.

## AUTHOR CONTRIBUTIONS

Conceptualization, M.D.; methodology, L.S., H.L.F., A.D., D.S., J.O., A.J.H., S.F., W.R., E.M.,

S.D.; validation, L.S., D.S.; investigation, L.S., H.L.F., A.D., D.S., J.O., A.J.H., A. G.-R., S.D.,

S.F., M.M., W.R., T.S., A.B.-B.; writing – original draft, M.D.; writing – review and editing, all authors; supervision, A.G.-R., S.D., W.B., E.M., A.B.-B., M.D.

## MATERIALS AND METHODS

RAW DATA for all Figure that display quantitative data are provided in the Supplemental Table.

The following MATERIALS were used.

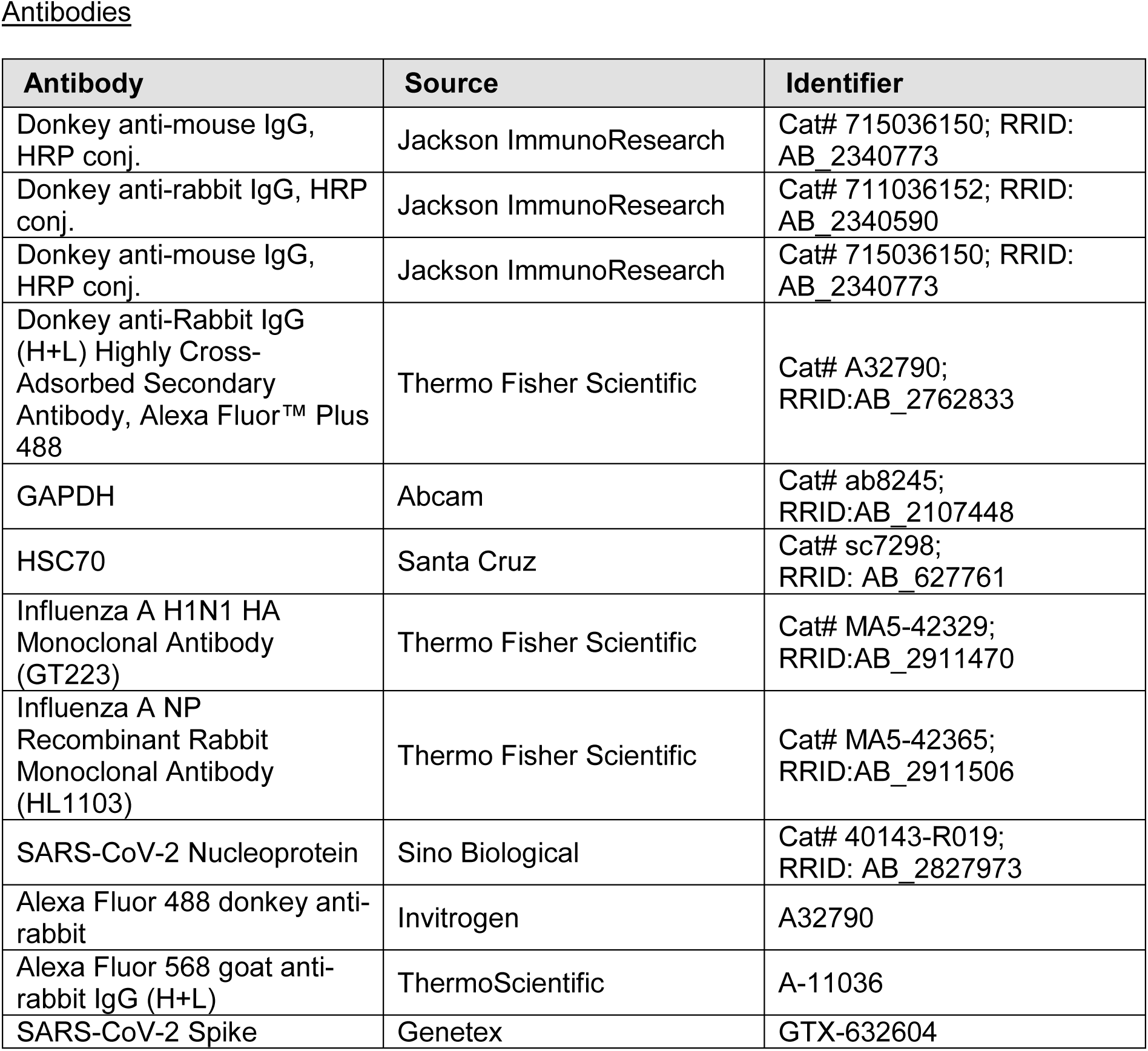
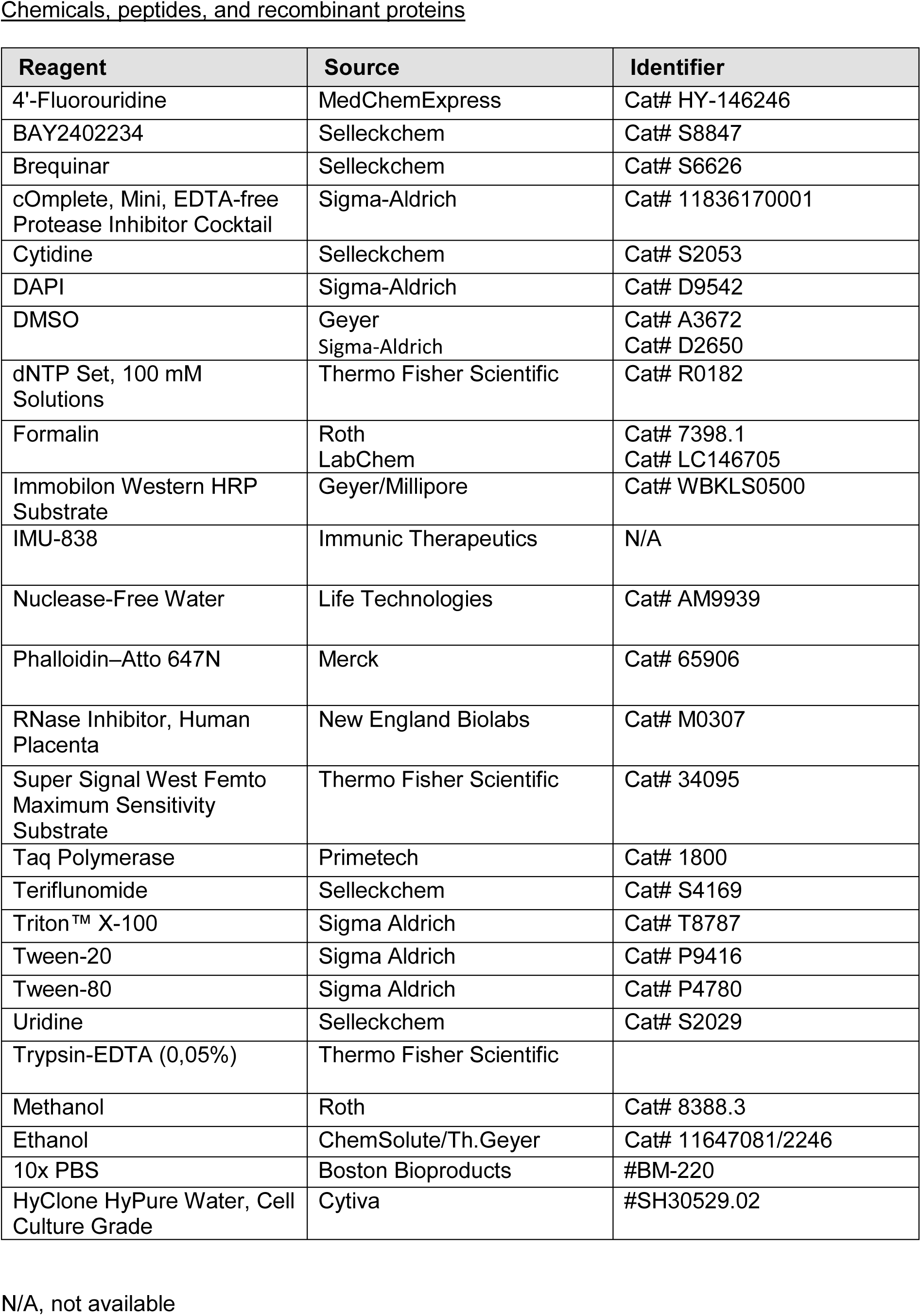
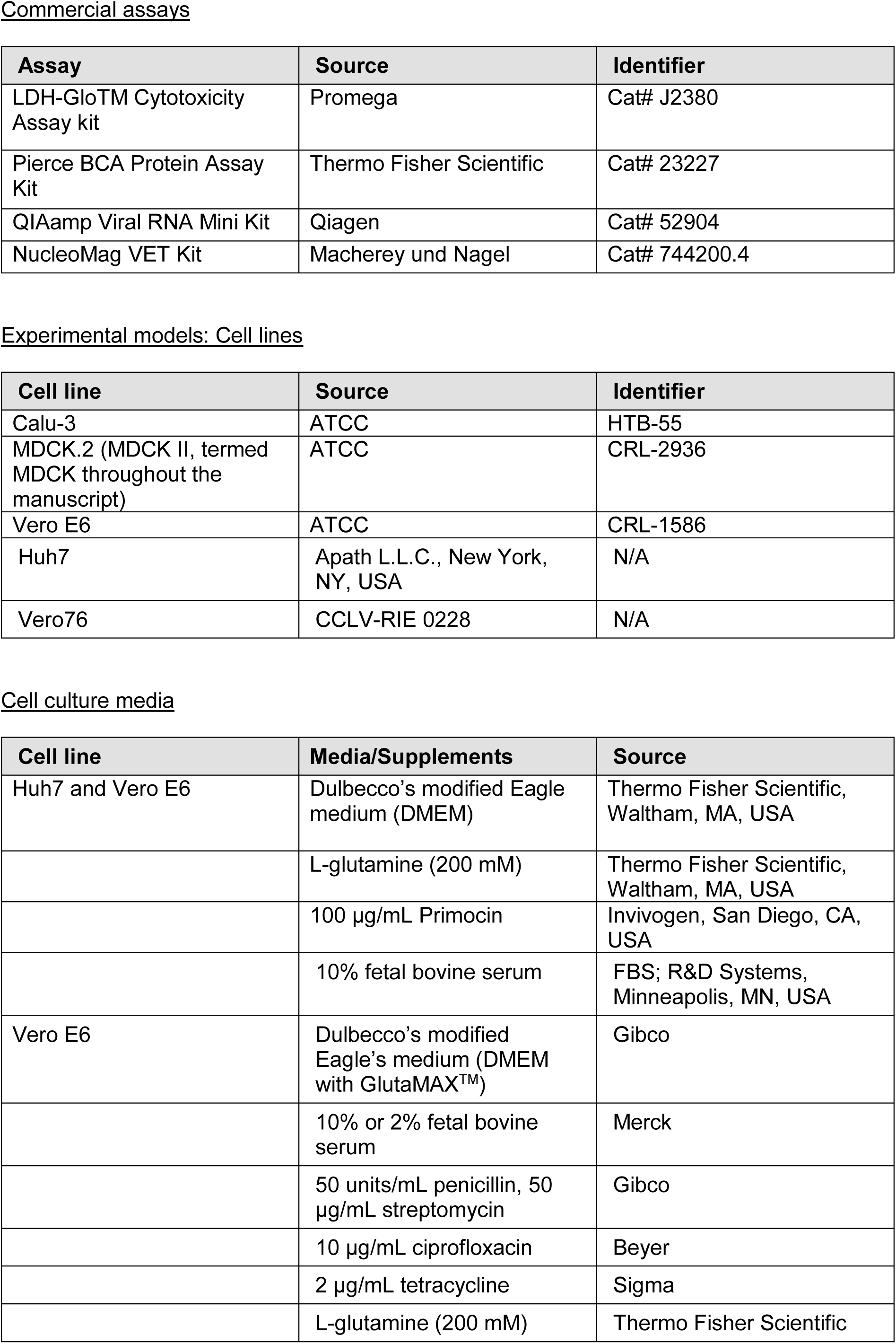
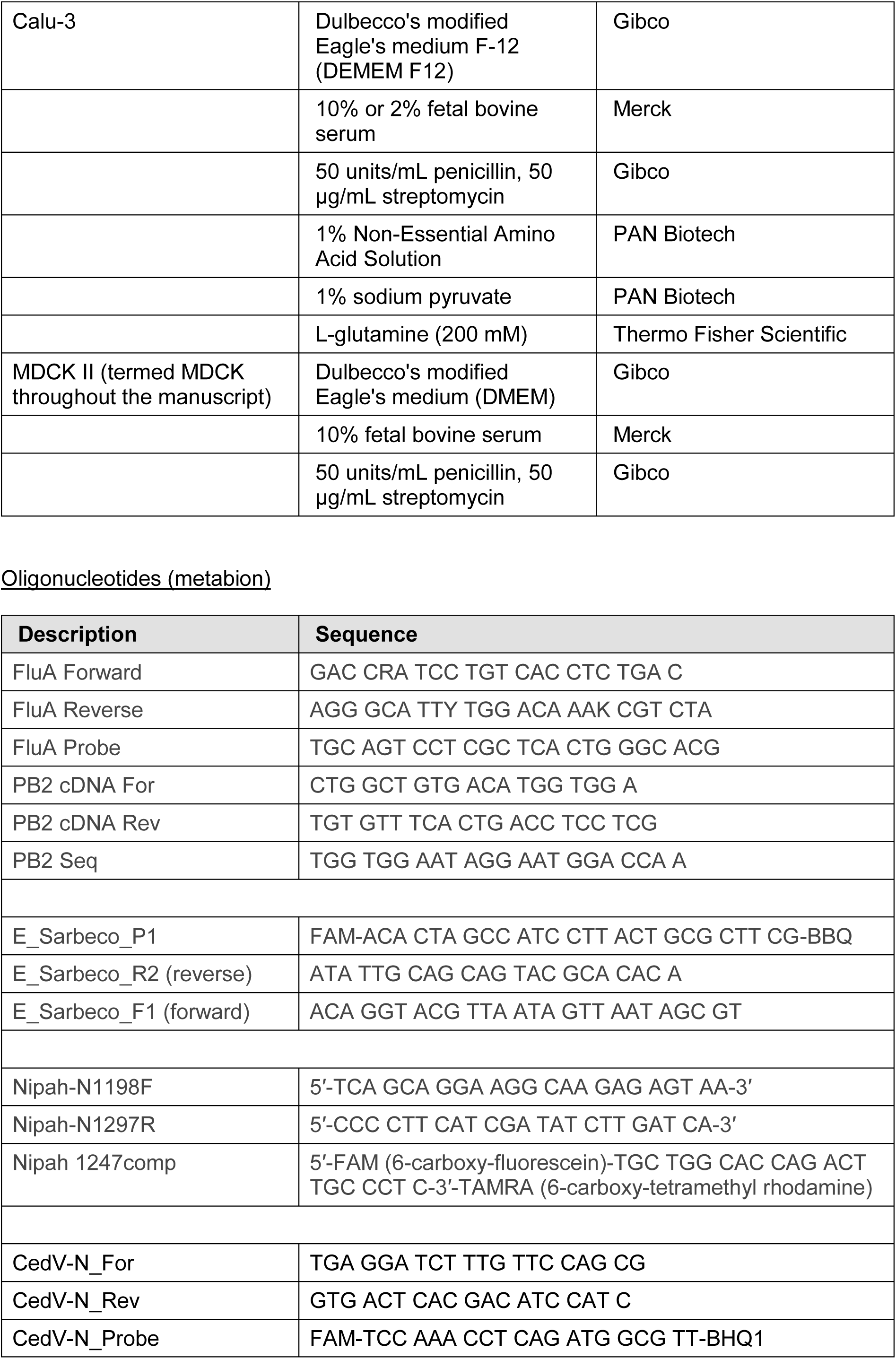
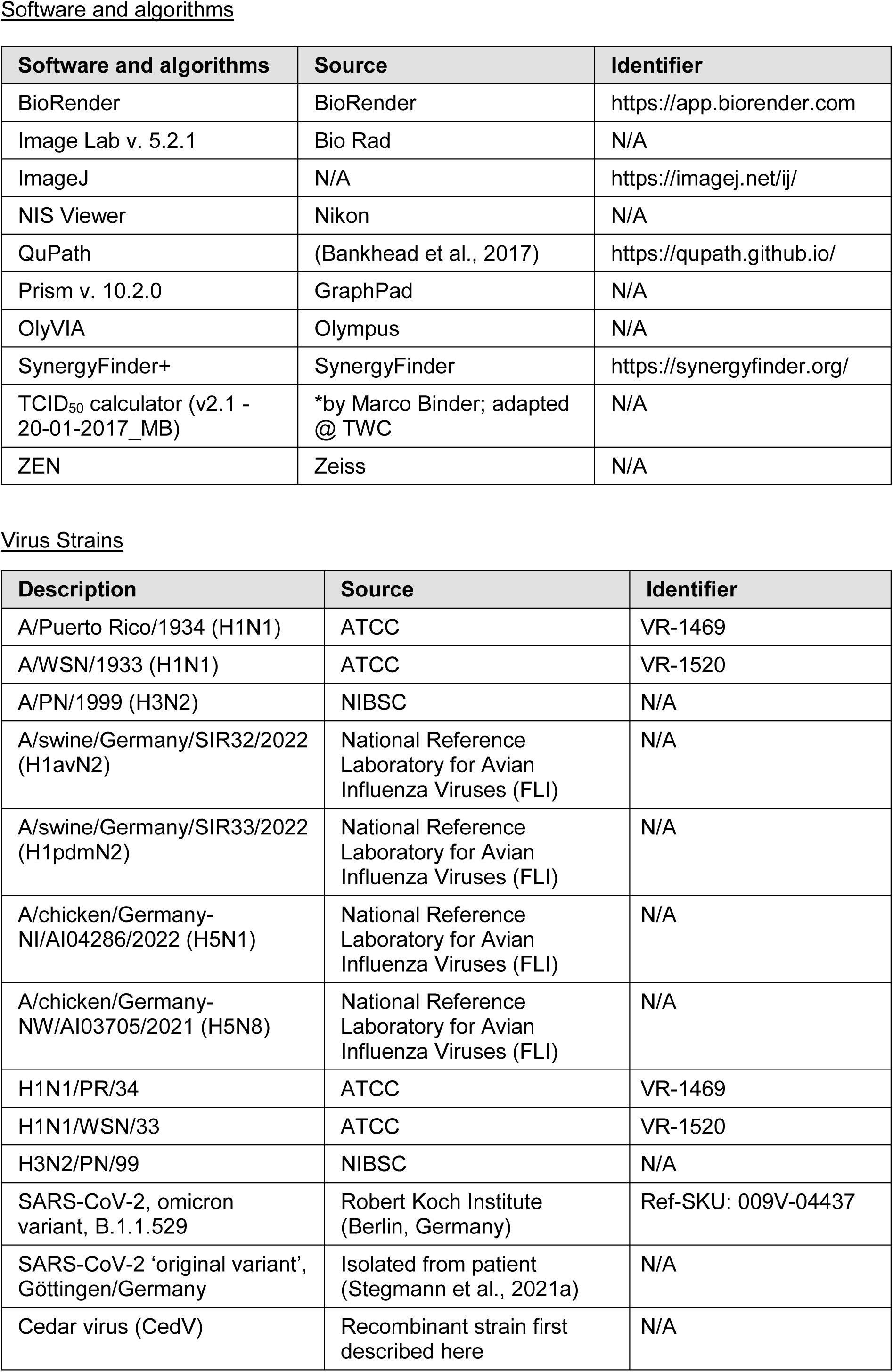
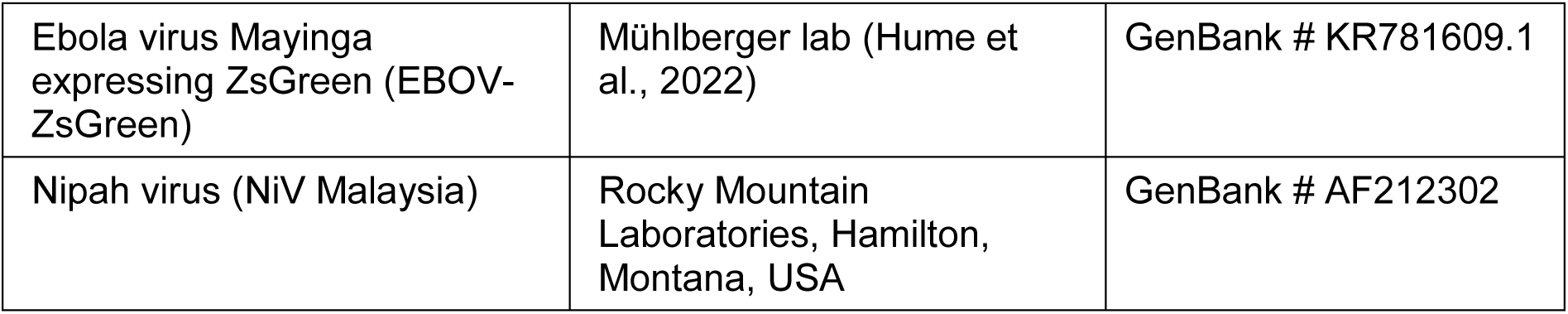

METHODS are provided separately for each virus investigated.

### Influenza A virus

#### Virus propagation

IAVs H1N1/WSN/33, H1N1/PR/34, and H3N2/PN/99 were kindly provided by Stefan Pohlmann (German Primate Center). IAVs H1avN2, H1pdmN2, H5N1, and H5N8 were isolated under safety precautions from deceased animals and kindly provided by Timm Harder (Friedrich-Loeffler-Institut). Virus stocks were grown and titrated on MDCK cells as described previously (Graaf et al., 2022). Virus infections were conducted in DMEM containing 0.1% FBS supplemented with 2 µg/mL TPCK trypsin (Sigma).

#### Cells

Madin-Darby Canine Kidney II (here referred to as MDCK; ATCC) cells were cultured in Dulbecco’s modified Eagle’s medium (DMEM; Gibco) supplemented with 10% fetal bovine serum (FBS; Merck), 50 units/mL penicillin (Gibco) and 50 µg/mL streptomycin (Gibco). Calu-3 cells were cultured in DMEM-F-12 Nutrient Mixture supplemented with 10% FBS, 50 units/mL penicillin, 50 µg/mL streptomycin, 200 mM L-glutamine (Thermo Fisher Scientific), 1% sodium pyruvate (PAN Biotech) and 1% Non-Essential Amino Acid Solution (PAN Biotech). Cells were cultured in a humidified atmosphere at 37°C with 5% CO2.

#### LDH Assay

Cytotoxicity of the compounds was evaluated using the LDH-GLO Cytotoxicity Assay Kit (Promega). In brief, 25,000 MDCK cells were seeded into a 24-well plate. The following day, cells were subjected to either single or combinatorial treatment with DHODH inhibitors and/or 4′-FlU. After 48 h of treatment, supernatant samples were collected. To determine the maximum possible LDH release, untreated cells were exposed to 10% Triton X-100 in phosphate-buffered saline (PBS) for 15 minutes. The luminescence signal was measured using a Berthold Centro LB963 plate reader.

#### Quantitative RT-PCR

To assess drug efficacy, 50,000 MDCK or Calu-3 cells were seeded into a 24-well plate and incubated overnight. Prior to infection, the medium was aspirated and replaced with DMEM containing 0.1% FBS. Cells were then infected with the corresponding virus strains at MOI 0.01 for 1 h. After the inoculation period, the virus inoculum was removed and cells were washed using PBS. Following infection, cells were exposed to increasing concentrations of DHODH inhibitors and/or 4′-FlU. After reaching the indicated time points, viral RNA was harvested from the supernatant using the QIAamp Viral RNA Mini Kit. Briefly, 140 µL of viral supernatant was lysed with 560 µL AVL-Lysis Buffer containing carrier RNA. Next, 630 µL of 99% ethanol was added and the mixture was vigorously vortexed at 1,400 rpm for 1 min. Now, 630 µL sample was loaded onto a column and centrifuged at 8,000 rpm for 1 min. This step was repeated until all the sample was processed. The column was washed with 500 µL Buffer 1 and centrifuged at 8,000 rpm for 1 min, followed by a wash using 500 µL Buffer 2 and centrifugation at 14,000 rpm for 3 min. Viral RNA was eluted using 60 uL RNAse free water. Thermal cycling was performed as described previously (Drosten, 2019). Primers used to amplify viral RNA were FluA Forward, FluA Reverse and FluA Probe, which target a highly conserved region in the IAV RNA encoding the Matrix protein.

For avian and porcine influenza viruses, the RNA extraction was performed using NucleoMagVet RNA/DNA isolation kit (Macherey & Nagel GmbH, Germany) in a KingFisher Flex Purification System (Thermo Fisher Scientific, USA). Virus RNA progeny was determined using qRT-PCR targeting the M1-gene of IAVs as described previously (Hassan et al., 2022; Spackman et al., 2003). Briefly, 1.25 μl RNAse free water, 6.25 μl 2x qRT-PCR Buffer and 0.5 μl RT-PCR Enzyme Mix (AgPath-ID One-Step qRT-PCR Reagents, Ambion-Applied Biosystems) was mixed. 1 μl -FAM-probe was added to target M1 and 1 μl EGFP-Mix4 (5)- HEX served as internal control.

#### Immunofluorescence

To quantify the percentage of infected cells, 50,000 MDCK cells were seeded into each well of an 8 well µ-slide (Ibidi) and infected with the respective virus at MOI 0.01 for 1 h. Following washing, DMEM containing the indicated drug concentrations was added to the cells. Upon reaching the designated time point, the virus-containing supernatant was removed, and the cells were fixed with 4% paraformaldehyde (PFA) for 1 h at room temperature (RT). After rinsing with PBS, cells were permeabilized in a 0.5% Triton X-100-PBS solution for 30 minutes and then rinsed twice with PBS. Non-specific binding sites were blocked by incubating cells in 10% FBS-PBS solution for 10 min. An unconjugated primary antibody against the viral nucleoprotein (Thermo Fisher Scientific, 1:2,500) was applied overnight at 4°C. After three PBS washes, slides were incubated with secondary Alexa Fluor 488 donkey anti-rabbit IgG (Thermo Fisher Scientific, 1:500) for 1 h at RT. F-actin was visualized using Phalloidin-Atto 647N (Sigma-Aldrich #65906, 1:1,000), and nuclei were stained with 4′,6-diamidino-2-phenylindole (DAPI) for 10 min at RT. Whole slides were imaged using the Zeiss Celldiscoverer 7 automated microscope. The percentage of infected cells was determined using the Zeiss Gene- and Protein-Expression module.

#### Tissue Culture Infectious Dose

To determine the Tissue Culture Infectious Dose (TCID50/mL), 10,000 MDCK cells per well were seeded into a 96-well plate and incubated overnight. Viral supernatant was generated as described in the previous sections. The following day, growth medium was replaced with fresh DMEM, and virus-containing supernatant was added in a 10-fold serial dilution, followed by 48 h incubation. Each sample was titrated in quadruplicate. Subsequently, plates were fixed with 4% PFA and infection was visualized by immunofluorescence staining as described above. Fluorescence signals were acquired using a Nexcelom Celigo Microplate Reader. Viral titers were quantified according to the Spearmann and Karber method (Karber, 1931).

#### Western Blot analysis

To investigate viral protein synthesis, 300,000 MDCK cells per well were seeded into 6-well plates and incubated overnight. The following day, the cells were infected with the virus at MOI 0.01 for 1 h. After washing with PBS, medium containing the specified drug concentrations was added and incubated for 24 h. The cells were then washed once with PBS and harvested using radioimmunoprecipitation assay (RIPA) lysis buffer (20 mM TRIS-HCl pH 7.5, 150 mM NaCl, 10 mM EDTA, 1% Triton X-100, 1% deoxycholate salt, 0.1% SDS, 2 M urea) supplemented with protease inhibitors. After sonication, the samples were mixed with Laemmli buffer and heated to 95°C for 5 min. Equal amounts of each sample were separated by electrophoresis on a 10% SDS page and transferred to a nitrocellulose membrane. Non-specific binding sites were blocked by incubating the membrane in 5% (w/v) nonfat milk in TBS containing 0.1% Tween-20 for 1 h. Subsequently, the membrane was incubated with primary antibodies at 4°C overnight. To visualize the binding, the membranes were incubated with HRP-conjugated secondary antibodies (donkey anti-rabbit IgG; Jackson ImmunoResearch) at RT for 1 h, followed by detection using Immobilon Western HRP Substrate.

#### Synergy Score

For synergy calculations, 100% viability was established as the baseline, representing Influenza A RNA generated without treatment. All other conditions were normalized and expressed as a percentage of the control. Highest Single Agent (HSA) (Berenbaum, 1989) scores were calculated using the https://synergyfinder.org/ web application. Each represented score resulted from three independent biological replicates. Synergy score values > 10 indicate synergism, while scores < -10 indicate an antagonistic effect; scores between -10 and 10 reflect additive effects of the drugs.

#### Selecting 4′-FlU resistant virus

For virus adaptation, MDCK cells were exposed to H1N1/PR/34 at MOI 0.1 for 1 h. Subsequently, the cells were washed with PBS, and the initial selection was started with 75 nM 4′-FlU. The viral populations were harvested 48 h post-infection and used for the next passage. Following each passage, the quantity of virus RNA copies/mL was assessed using qRT-PCR. Subsequent infections were performed using RNA copies equivalent to an MOI of 0.1. The concentration of 4′-FlU was doubled after each passage, culminating in a dose escalation test reaching a concentration of 6000 nM in the final passage. As a control, the same virus strain was passaged under identical conditions but without 4′-FlU.

#### Sanger Sequencing

To investigate the potential presence of resistance-conferring mutations, as previously outlined (Lieber et al., 2024), cDNA was synthesized for all regions of interest in resistant and wild-type virus populations. Subsequently, the viral cDNA products underwent Sanger sequencing for mutation analysis.

#### Statistical Analysis

Data analysis was performed using GraphPad Prism 10 for Windows (GraphPad Software LLC. San Diego, CA, USA). For experiments characterized by exponential growth kinetics (e.g., titration and qRT-PCR), ordinary one-way ANOVA test was performed using the geometric mean, followed by Dunnett’s multiple comparisons test. Conversely, experiments with more linear kinetics (e.g., immunofluorescence and LDH-Assay) utilized the arithmetic mean for analysis. Ns, not significant; *, P≤0.05; **, P≤0.01; ***, P≤0.001; ****, P≤0.0001.

### SARS-CoV-2

#### Cell culture

Vero E6 cells were cultivated in Dulbecco’s modified Eagle’s medium (DMEM with GlutaMAX^TM^, Gibco) that was supplemented with 10% fetal bovine serum (Merck), 50 units/mL penicillin, 50 μg/mL streptomycin (Gibco), 200 mM L-glutamine (Thermo Fisher Scientific), 2 μg/mL tetracycline (Sigma) and 10 μg/mL ciprofloxacin (Bayer). The cells were maintained at 37°C in a humidified atmosphere with 5% CO2.

#### Treatment and infection

Cells were seeded into the cell culture dish as outlined below.

| Cell culture dish | Cell amounts |
| --- | --- |
| 24-well-plates (Corning) | 20,000 cells per well |
| 8-well chamber slides (ibidi) | 5,000 cells per well |
| 96-well plates (Greiner) | 3,000 cells per well |

Following 24 h incubation at 37°C, media was exchanged to Dulbecco’s modified Eagle’s medium (DMEM with GlutaMAX^TM^, Gibco) supplemented with only 2% fetal bovine serum (Merck), 200 mM L-glutamine (Thermo Fisher Scientific), 50 units/mL penicillin, 50 μg/mL streptomycin (Gibco), 2 μg/mL tetracycline (Sigma) and 10 μg/mL ciprofloxacin (Bayer), and cells were infected with SARS-CoV-2 Wuhan-like original strain (Stegmann et al., 2021a) at MOI 0.1. After 1 h, brequinar (Selleckchem S6626), BAY2402234 (Selleckchem S8847) or 4’-FlU (Hycultec HY-146246) were added at varying concentrations. Cells were incubated for 48 h at 37°C.

#### Quantitative RT-PCR

In order to quantify the viral RNA, samples of the cell culture supernatant were harvested, and total RNA was isolated using the QIAamp Viral RNA Kit (Qiagen, 52904). qRT-PCR was performed, using a TaqMan probe (Corman et al., 2020) purchased from Eurofins that binds to the amplified region corresponding to the viral gene that encodes the envelope protein (position 26,141-26,253 in the SARS-CoV-2 genome). The resulting Ct values were used to calculate the virus RNA progeny. The amount of viral RNA found in the supernatant of SARS-CoV-2-infected but untreated cells was defined as 100% and used to normalize the amounts obtained after each treatment.

#### Synergy Score

Synergy of the Virus RNA progeny, calculated from the Ct values obtained by qRT-PCR, was determined using the online tool SynergyFinder https://synergyfinder.fimm.fi/ (Ianevski et al., 2020).

#### Tissue Culture Infectious Dose 50

Virus-containing supernatant of infected and treated cells was titrated onto Vero E6 cells in a 96-well plate (end point dilution assay). This setup was used to determine the Tissue Culture Infectious Dose 50 (TCID50) per mL. Immunofluorescent staining of the viral nucleocapsid protein and the cell nucleus was performed as indicated below. The cells were then scanned using a Celfgo® S Imaging Cytometer (Nexcelom Bioscience).

#### Immunofluorescence analyses

Vero E6 cells, seeded in 8-well chamber slides, were infected with SARS-CoV-2 (MOI 0.1) and treated as above. Following 48 h incubation at 37°C, cells were washed with PBS and fixed for 1 h at room temperature with 4% formaldehyde in PBS. Fixed cells were permeabilized for 30 min with 0.5% Triton X-100, washed twice with PBS and blocked with 10% FCS in PBS. Primary and secondary antibodies were diluted in blocking solution. Cells were incubated for 1 h with a primary antibody against Nucleoprotein (N; Sino Biological #40143-R019, 1:8000), washed twice with PBS and incubated for 1 h with a corresponding secondary antibody, DAPI (Sigma; 1:3,000) and Phalloidin–Atto 647N (Sigma; 1:500). After washing, the cells were kept in PBS until microscopy. Scans of the entire well were taken with the CellDiscoverer7 (Zeiss) using 5x magnification, and they were used to quantify virus-infected cells in an automated fashion. Nuclei were counted for total cell number and cells expressing NP were counted providing the amount of virus-infected cells (Bioapp: Gene- and Protein expression, Zen desk, Zeiss).

#### Immunoblot analysis

Protein samples were harvested by washing cells with PBS before lysis in radioimmunoprecipitation assay (RIPA) lysis buffer with additional protease inhibitors. Pierce BCA Protein assay kit (Thermo Fisher Scientific) was used to determine the total protein concentrations, in order to match protein amounts between all samples. After denaturation for 5 min at 95°C in Laemmli buffer, the proteins were separated according to size using sodium dodecyl sulfate–polyacrylamide gel electrophoresis (SDS-PAGE). Further on, the proteins were transferred to a nitrocellulose membrane. The membrane was blocked in 5% skim milk in TBS with 0.1% Tween 20 for 1 h before incubation in primary antibodies over night at 4°C. After washing three times, membranes were incubated in secondary antibodies at room temperature for 1 h (donkey anti-rabbit IgG or donkey anti-mouse IgG, Jackson Immunoresearch). Super Signal West Femto Maximum Sensitivity Substrate (Thermo Fisher) and Immobilon Western Substrate (Millipore) were used to detect the respective proteins in the ChemiDoc Imaging SystemTM MP (BioRad).

#### Quantification of LDH release to determine cytotoxicity

Vero E6 cells were seeded in a 24-well plate and treated as stated above. After 48 h incubation at 37°C, the cell supernatant was used to determine the LDH release of the cells using the LDH-Glo™ Cytotoxicity Assay kit (Promega). The bioluminescence corresponding to the LDH content was quantified using a Luminometer (Berthold Celtro). A maximum LDH release control was produced by incubating untreated cells with 2 µL of 10% Trition X-100 for 10 minutes and set to 100% cytotoxicity. Cytotoxicity was calculated with following formula:

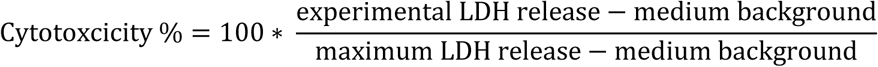

#### Statistical Analysis

Described in the section ‘Statistical Analysis’ for Influenza virus.

### SARS-CoV-2 infection, hamster model

#### Ethics statement

Hamster experiments were carried out according to the German Regulations for Animal Welfare after obtaining the necessary approval from the authorized ethics committee of the State Office of Agriculture, Food Safety and Fishery in Mecklenburg-Western Pomerania (LALLF MV) under permission number 7221.3-1-049/20 and approval of the commissioner for animal welfare at the Friedrich Loeffler Institute (FLI), representing the Institutional Animal Care and Use Committee (IACUC).

#### Infection

Male Golden Syrian Hamsters (*Mesocricetus auratus*; RjHan:AURA; 80–100 g; Janvier Labs, France) were orotracheally inoculated with 10^3^ TCID50 SARS-CoV-2 strain Germany/BavPat1/2020 (BavPat1), GISAID accession EPI_ISL_406862, in a volume of 100 µL under brief isoflurane anesthesia as described previously (Stegmann et al., 2022). Oral swab samples were collected at -1, 1, 3 and 5 days post infection (dpi) and were collected in 500 µl DMEM+P/S, while nasal washes (200 µl sterile PBS) were collected at 2, 4 and 6 dpi under brief isoflurane anesthesia. Here, PBS was carefully flushed through the nose and collected in a 1.5 ml tube. At 7 dpi, animals were sacrificed and samples of the respiratory tract were collected for virological and histological analysis.

#### Determination of virus RNA and infectious particles in tissues

Total RNA extraction as well as qRT-PCR of swab samples, nasal washes and tissue samples and determination of TCID50 virus titer were performed as previously published (Blaurock et ^a^l., 202^2^).

#### Pathology

Paraffin embedded lung lobes and tracheal sections were submitted to the Department of Pathology, University of Veterinary Medicine Hannover, Hannover, Germany. Tissue blocks were trimmed and stained with hematoxylin-eosin according to routine protocols. Histopathological assessment of respiratory lesions was performed by two veterinary pathologists (T.S., W.B.) semiquantitatively according to routine protocols with minor modifications (Armando et al., 2022).

In summary, alveolar, vascular and conductive airway lesions were scored in the lungs. Multiplication of their extent and severity was used to generate combined scores before summing up the scores for all parameters.

GraphPad Prism software version 10 for Windows™ (GraphPad software, San Diego, California, USA) was used for statistical analysis and graph preparation. Total scores of all groups were compared by Kruskal-Wallis multi-comparison test with subsequent pairwise Mann-Whitney-U-tests and Benjamini-Hochberg correction for multiple comparisons (False Detection Rate: 0.05). Exact p values ≤ 0.05 were assumed statistically significant.

SARS-CoV-2 nucleoprotein immunohistochemistry was performed according to established protocols (Armando et al., 2022). After deparaffinization and rehydration, sections were incubated with a mouse monoclonal antibody against SARS-CoV-2 nucleoprotein (Sino Biological, Peking, China-40143-MM05). For negative controls, the antibody was replaced by the respective protein concentration of ascitic fluid from non-immunized BALB/cJ mice. Immunolabeling was visualized using the EnVision+ polymer system (Dako Agilent Pathology Solutions) and 3,3’-diaminobenzidine tetrahydrochloride as chromogen (Sigma Aldrich).

All slides were evaluated with a Zeiss Axioscope (Zeiss, Gottingen, Germany; field of view at 400x magnification: 0.16 mm2). Whole slides were scanned with the Olympus VS200 slide scanner (Olympus Deutschland GmbH, Hamburg, Germany) and images of animals representing the group median score were exported with the respective OlyVIA software.

### Henipaviruses: Cedar and Nipah virus

All work with live Nipah virus was performed in the BSL4 laboratory at the Friedrich-Loeffler-Institut. Cedar virus (CedV) was handled at BSL2.

#### Generation of recombinant Cedar virus

Recombinant CedV was generated from a plasmid containing the complete genome sequence of the CedV isolate CG1a (Genbank NC_025351.1f), described previously (Marsh et al., 2012). The full-length cDNA was obtained by insertion of synthetically generated DNA sequences (Eurofins Genomics, Germany) in a pBluescript II plasmid (Stratagene) derivative comprising the T7-RNA-polymerase promoter and downstream hepatitis delta virus ribozyme and T7-transcription terminator sequences as described before (Schnell et al., 1994). Support plasmids for the expression of CedV N, P, and L proteins were generated by insertion of synthetic gene sequences in the expression vector pCAGGS (Niwa et al., 1991). For the rescue of CedV, 10^6^ BSR-T7/5 cells (Buchholz et al., 1999) were transfected with 3.5 µg full-length cDNA plasmid with pCAGGS CedV-N (1.25 µg), -P (0.8 µg) and -L (0.4 µg) by Lipofectamin 2000 (Thermo Fisher Scientific). After 24 h incubation at 37°C and 5 % CO2, the cells were scraped off the 6-well cell culture plate and transferred to a T-25 tissue flask. Appearance of cell syncytia as a sign of virus replication and spread was microscopically monitored for 4 to 6 days. When 70-90 % of the cells formed syncytia, the supernatant was harvested and used for the production of virus stocks on BSR-T7/5 cells.

#### Drug treatment and infection

Vero 76 cells were seeded in 24-well plates for viral infection experiments. The next day, the cells were washed once with PBS to remove any residual media or debris. Following this, the cells were infected with the virus at a multiplicity of infection (MOI) of 0.001 (CedV) or MOI 0.2 (NiV) for 1 h. Then, the virus inoculum was removed and the cells were washed again with PBS to remove unbound viruses. Fresh medium containing the specified inhibitors in the indicated concentrations was then added in ZB5 supplemented with 2.5% FCS. After 24 h (NiV) or 48 h (CedV) incubation, supernatant was harvested and stored at -80°C until further processing. Experiments were performed in duplicates over three independent rounds.

#### Quantitative RT-PCR

Viral RNA was prepared using the QIAamp Viral RNA Kit (Qiagen). For both RT-qPCRs, we used the QuantiTect-Probe RT-PCR Kit (Qiagen). NiV RNA was analyzed as described (Mungall et al., 2006) using the primers Nipah-N1198F and Nipah-N1297R and the fluorogenic probe (TaqMan) Nipah 1247comp (cf. Materials). For the detection of CedV RNA, we used the following set of primers and probe targeting a region in the viral nucleoprotein (N) gene: CedV-N_For and CedV-N_Rev, as well as a probe with a 5′ 6Carboxyfluoresceine (FAM) reporter dye and a 3′ Black Hole Quencher (BHQ1) CedV-N_Probe (cf. Materials).

#### Plaque Assay

For titration, Vero 76 cells were seeded into 12-well plates and grown until fully confluent. Viral supernatant was titrated in tenfold dilutions in duplicates. Cells were infected for 1 h, then virus inoculum was removed, and cells were overlayed with 1% carboymethylcellulose (Sigma-Aldrich, Germany) in DMEM with 2.5 % FCS and incubated for 4 days. After the incubation period, the overlay was removed, and the cells were fixed with 10% formalin for 1 h. Following fixation, the cells were stained with 0.5% crystal violet in ethanol for 30 min to visualize and quantify the viral plaques.

### Ebola virus

#### Biosafety statement

All work with EBOV was performed in the BSL-4 facility of Boston University’s National Emerging Infectious Diseases Laboratories (NEIDL) following approved SOPs in compliance with local and federal regulations pertaining to handling BSL-4 pathogens and Select Agents.

#### Cell lines

Huh7 cells (kindly provided by Apath L.L.C., New York, NY, USA) and Vero E6 cells (ATCC, Manassas, VA, USA; CRL-1586) were maintained in Dulbecco’s modified Eagle medium (DMEM; Thermo Fisher Scientific, Waltham, MA, USA) supplemented with L-glutamine (200 mM; Thermo Fisher Scientific, Waltham, MA, USA), 100 µg/mL Primocin (Invivogen, San Diego, CA, USA), and 10% fetal bovine serum (FBS; R&D Systems, Minneapolis, MN, USA). Cells were grown at 37°C and 5% CO2.

#### Virus propagation

Recombinant Ebola virus (strain Mayinga) expressing ZsGreen (EBOV-ZsGreen; GenBank number KR781609.1) has been described before (Hume et al., 2022). EBOV-ZsGreen was propagated in Vero E6 cells in cell culture medium supplemented with 2% FBS. Virus titers were determined in Vero E6 cells by TCID50 assay using the Spearman and Karber algorithm (Karber, 1931; Ramakrishnan, 2016).

#### Drug treatment and infection

1x10^4^ Huh7 cells were seeded per well in 96-well plates. One day later, the cells were pre-treated with the indicated compounds (150 µl/well) for 1 h at 37°C and 5% CO2. 4′-FIU was resuspended in H2O and serially diluted to indicated concentrations in cell culture medium supplemented with 2% FBS. Brequinar (SelleckChem, Houston, TX) was resuspended in DMSO and diluted in cell culture medium supplemented with 2% FBS or 4′-FIU-containing cell culture medium supplemented with 2% FBS to prepare compound mixtures. After 1 h pretreatment, the cells were infected with EBOV-ZsGreen at an MOI of 0.1 without removing the compound-containing cell culture medium. The cells were incubated for 2 days and fixed for further analysis. 4′-FIU was replenished by adding fresh 4′-FIU to the compound-containing cell culture media at 1 dpi. The experiment was performed 3 times independently with each experimental condition setup in triplicate wells.

#### Immunofluorescence and quantification

EBOV-ZsGreen-infected cells were fixed 2 dpi with 10% formalin for at least 6 h. To stain the cell nuclei, the cells were washed four times in PBS and incubated with 4’,6-diamidino-2-phenylindole (DAPI; Sigma-Aldrich) in 200 ng/mL PBS for 15 minutes at room temperature. Images were acquired using a Nikon Eclipse Ti2 microscope with Photometrics Prime BSI camera and NIS Elements AR software. Infection rates (% ZsGreen expressing cells of total cells as stained by DAPI) were determined using QuPath software (https://qupath.github.io/).

#### Synergy Score

Synergy was calculated for the values obtained while quantifying the infected cells, using the online tool SynergyFinder https://synergyfinder.fimm.fi/ (Ianevski et al., 2020).

#### Statistical Analysis

These analyses were performed as described in the section ‘Statistical Analysis’ for Influenza.

---

**S1.**
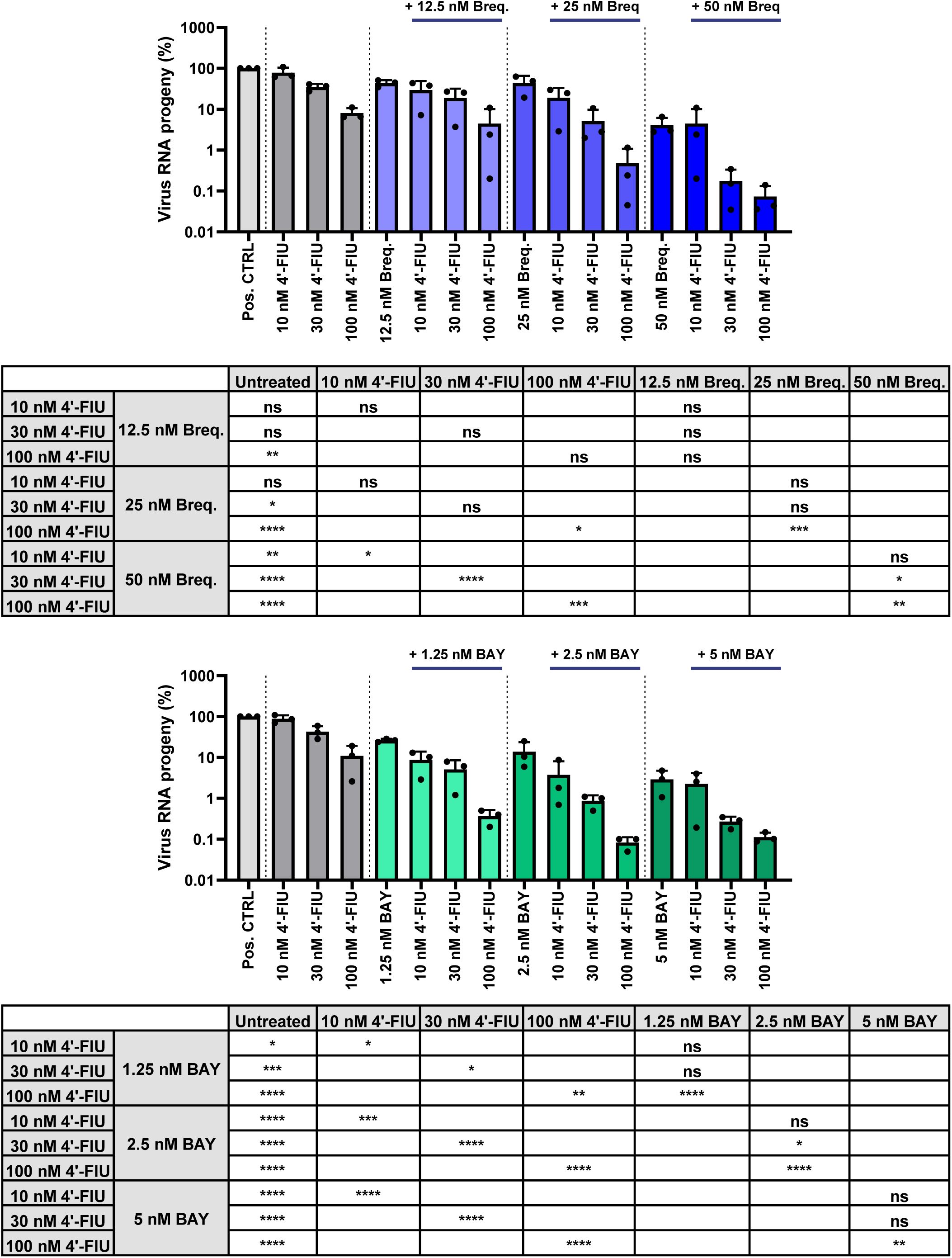
display the quantifications of virus RNA that are displayed more concisely in the indicated corresponding main figures.

**S2.**
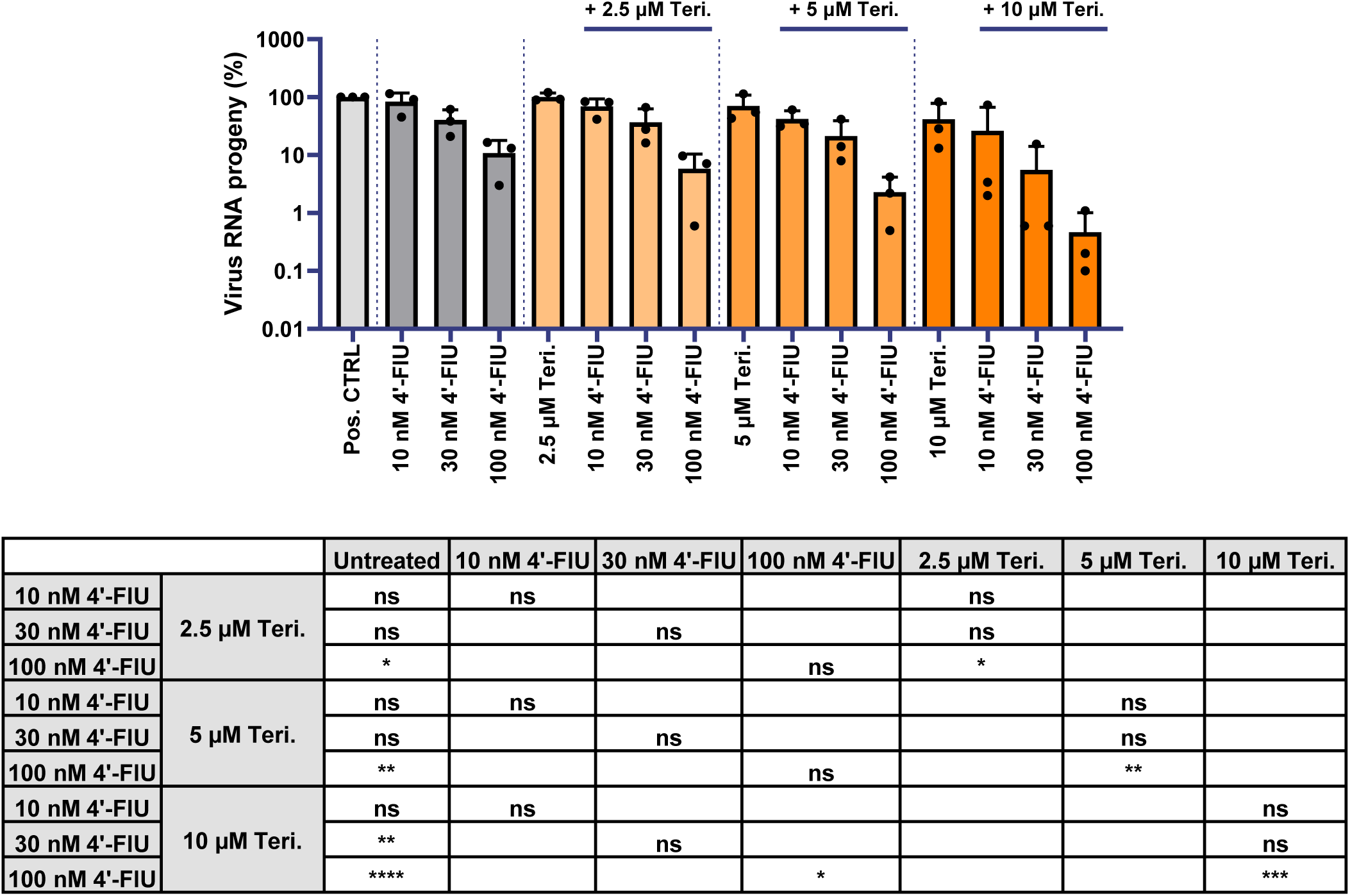
display the quantifications of virus RNA that are displayed more concisely in the indicated corresponding main figures.

**S3.**
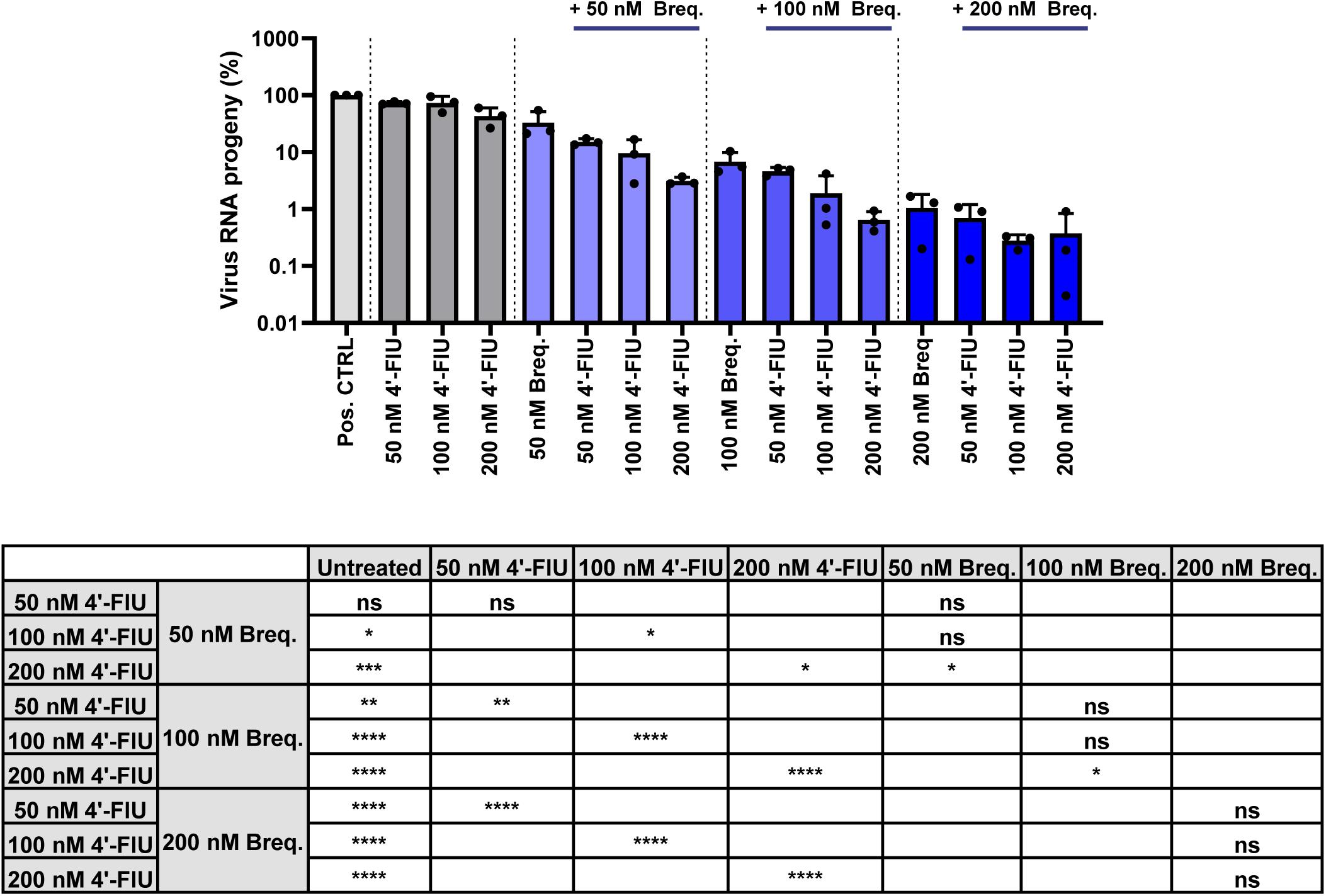
display the quantifications of virus RNA that are displayed more concisely in the indicated corresponding main figures.

**S4.**
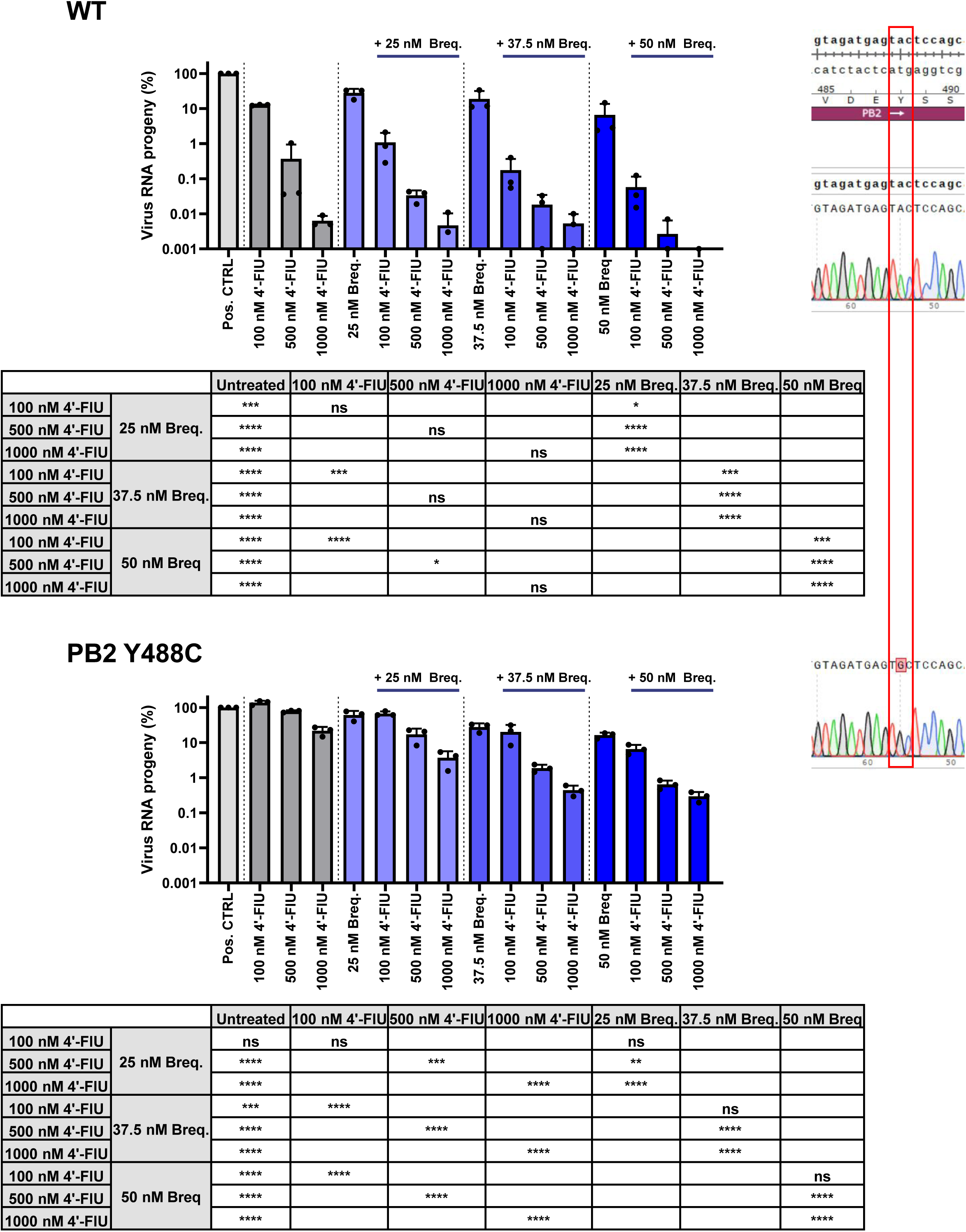
display the quantifications of virus RNA that are displayed more concisely in the indicated corresponding main figures.

**S5.**
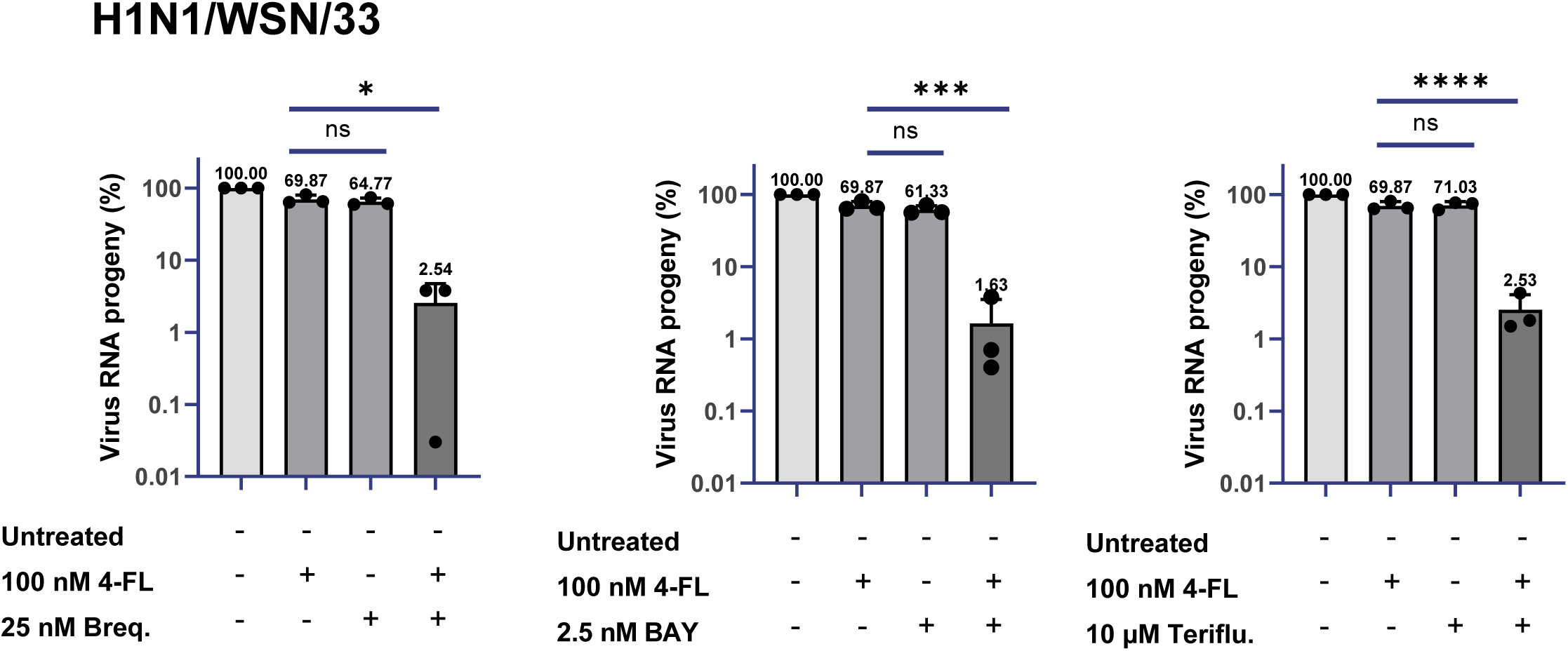
display the quantifications of virus RNA that are displayed more concisely in the indicated corresponding main figures.

**S6.**
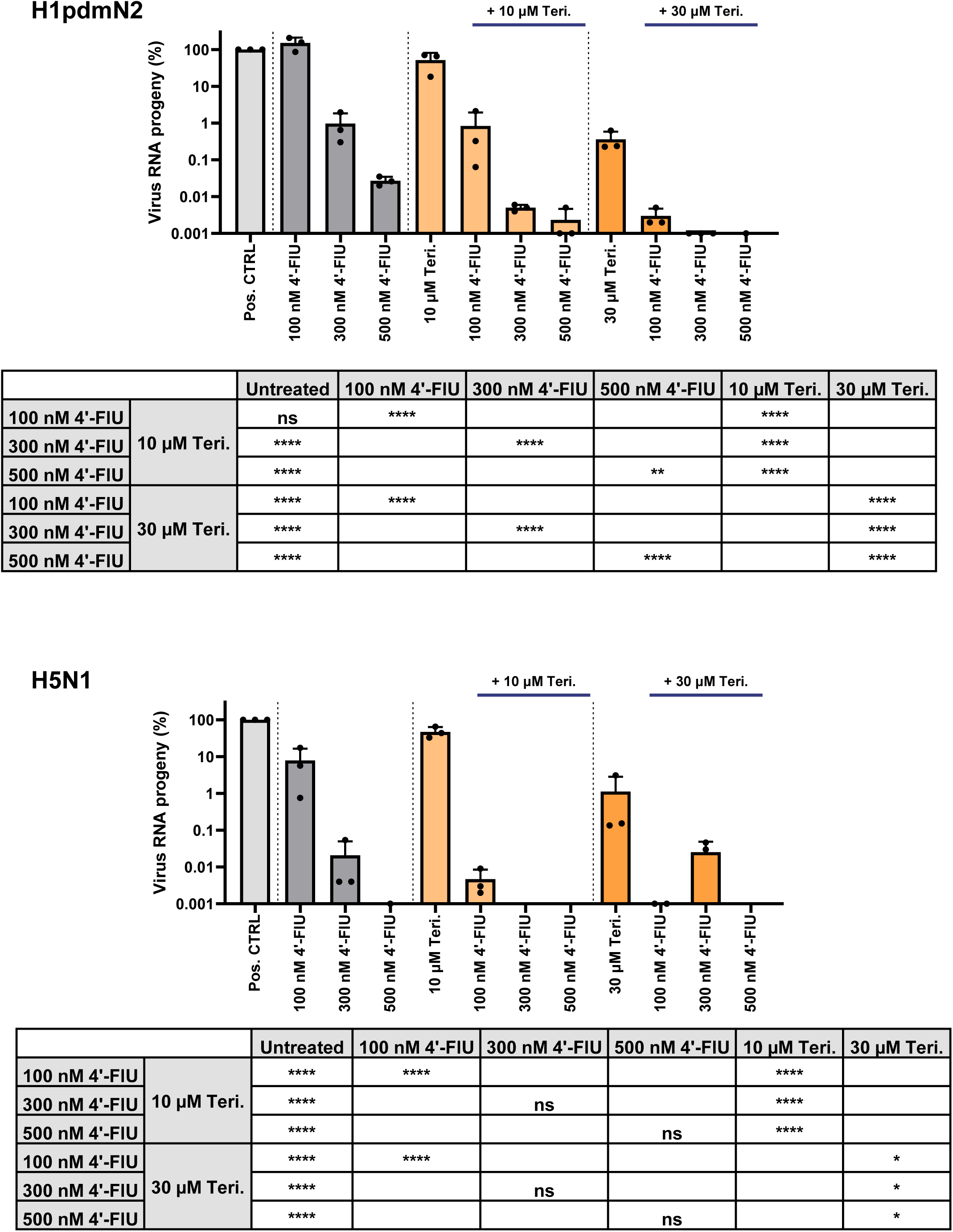
display the quantifications of virus RNA that are displayed more concisely in the indicated corresponding main figures.

**S7.**
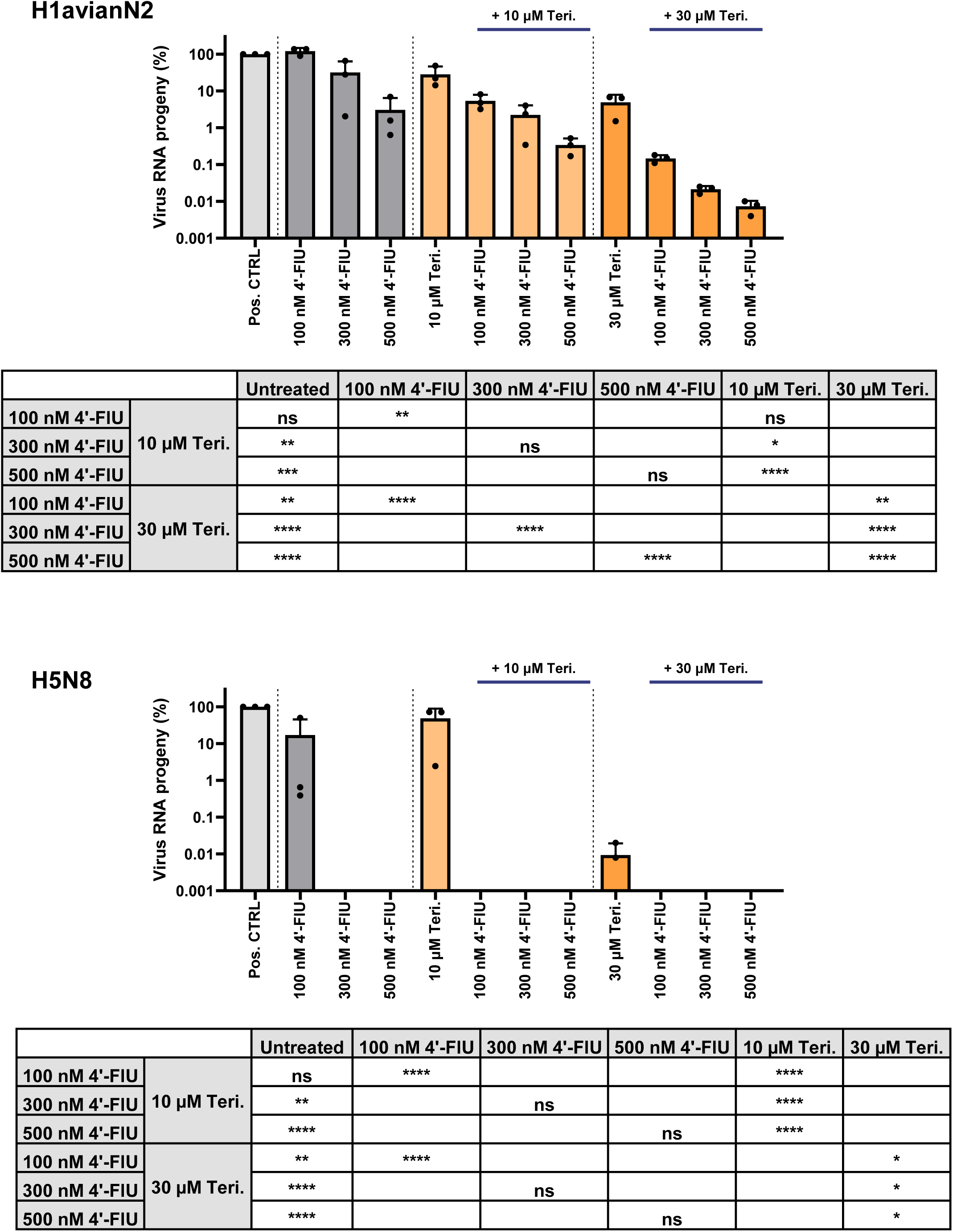
display the quantifications of virus RNA that are displayed more concisely in the indicated corresponding main figures.

**S8.**
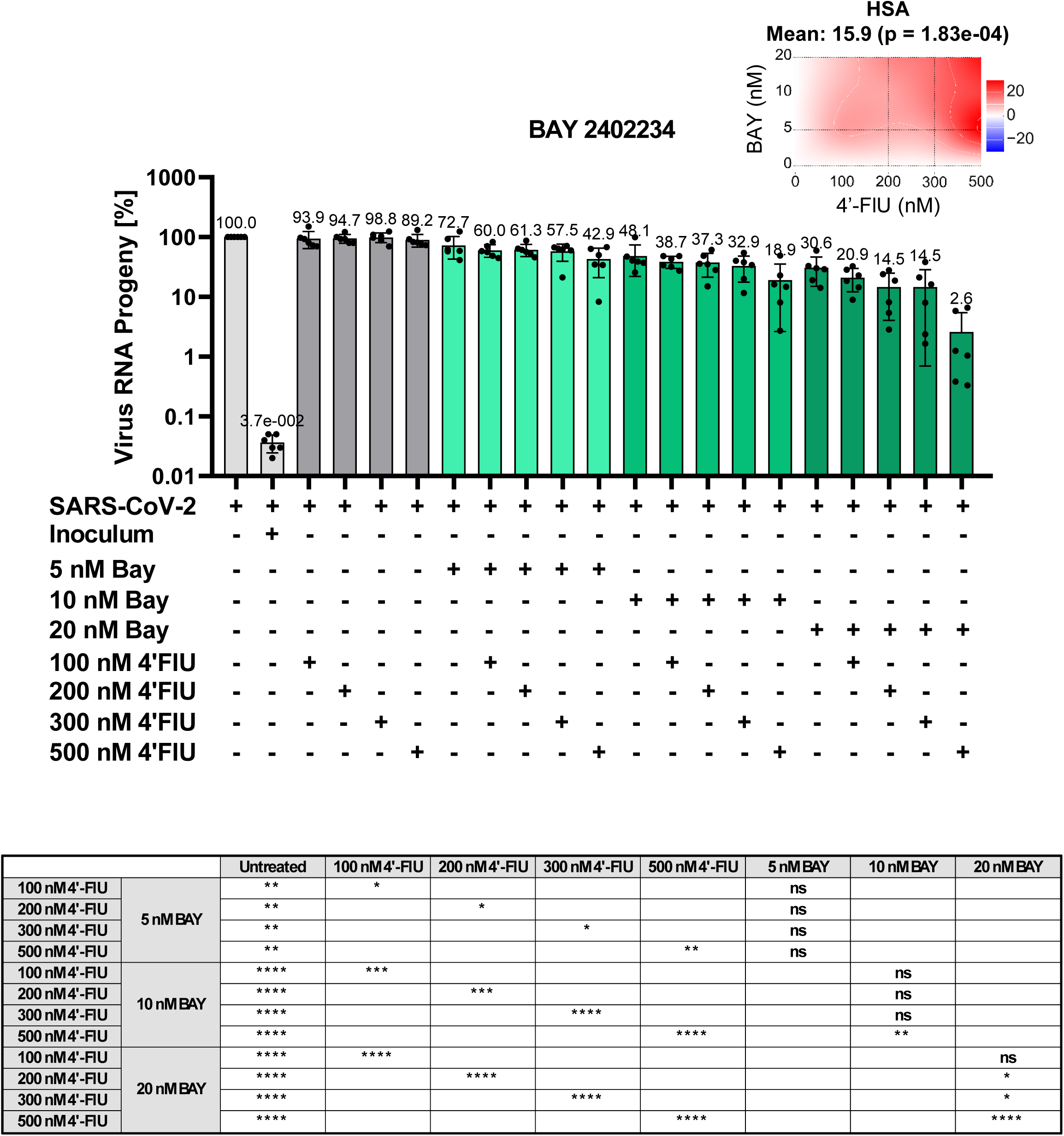
display the quantifications of virus RNA that are displayed more concisely in the indicated corresponding main figures.

**S9.**
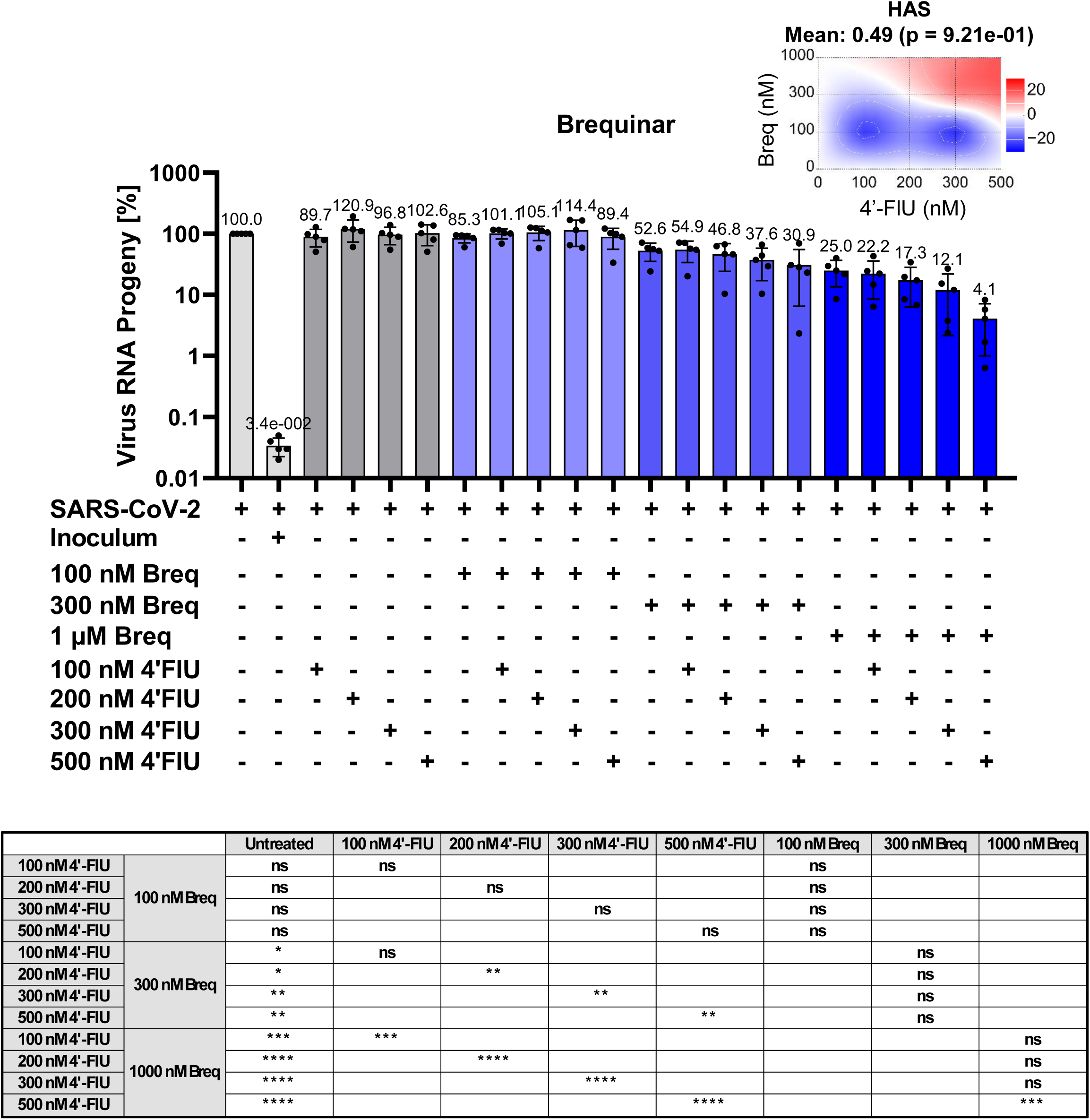
display the quantifications of virus RNA that are displayed more concisely in the indicated corresponding main figures.

**S10.**
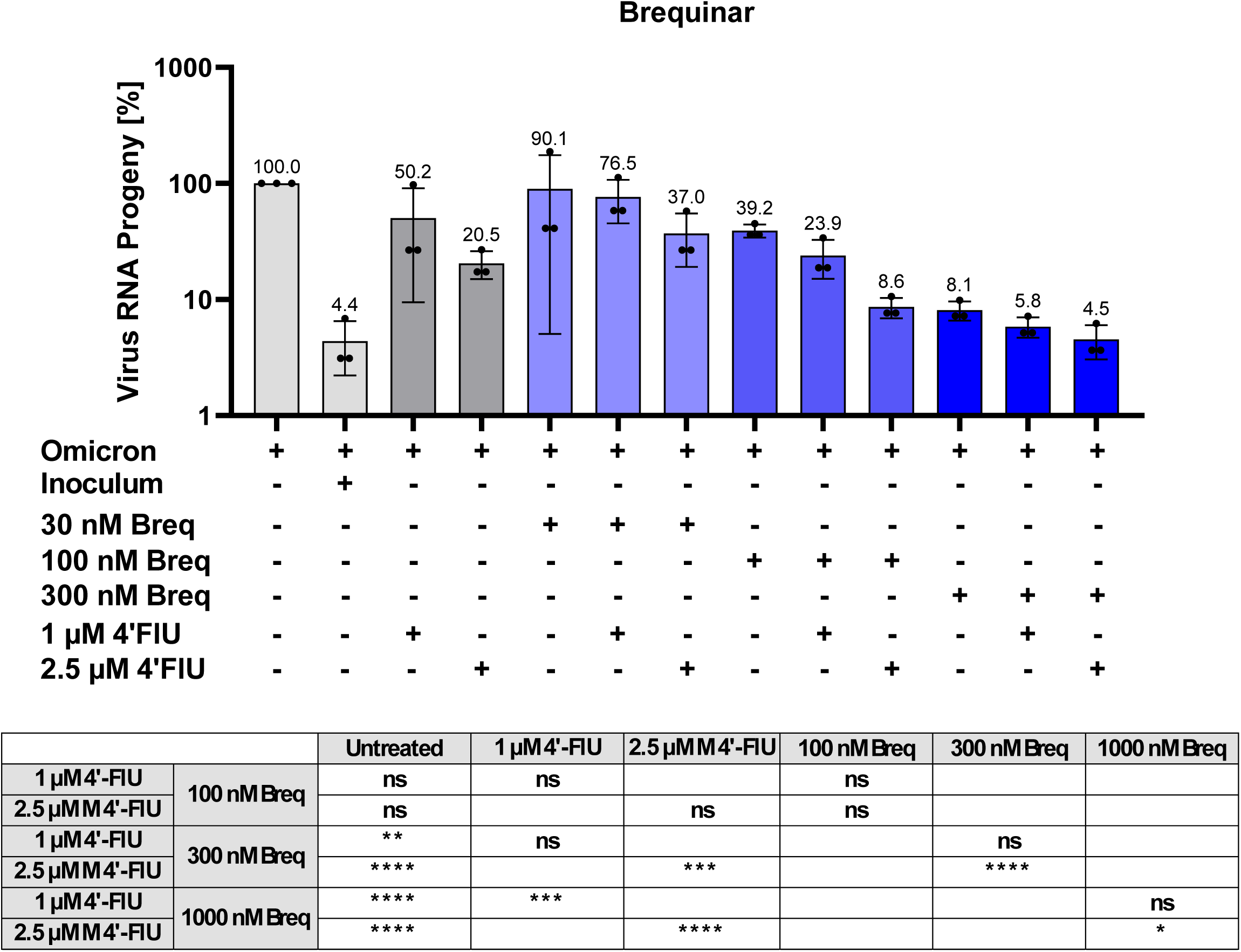
display the quantifications of virus RNA that are displayed more concisely in the indicated corresponding main figures.

**S11.**
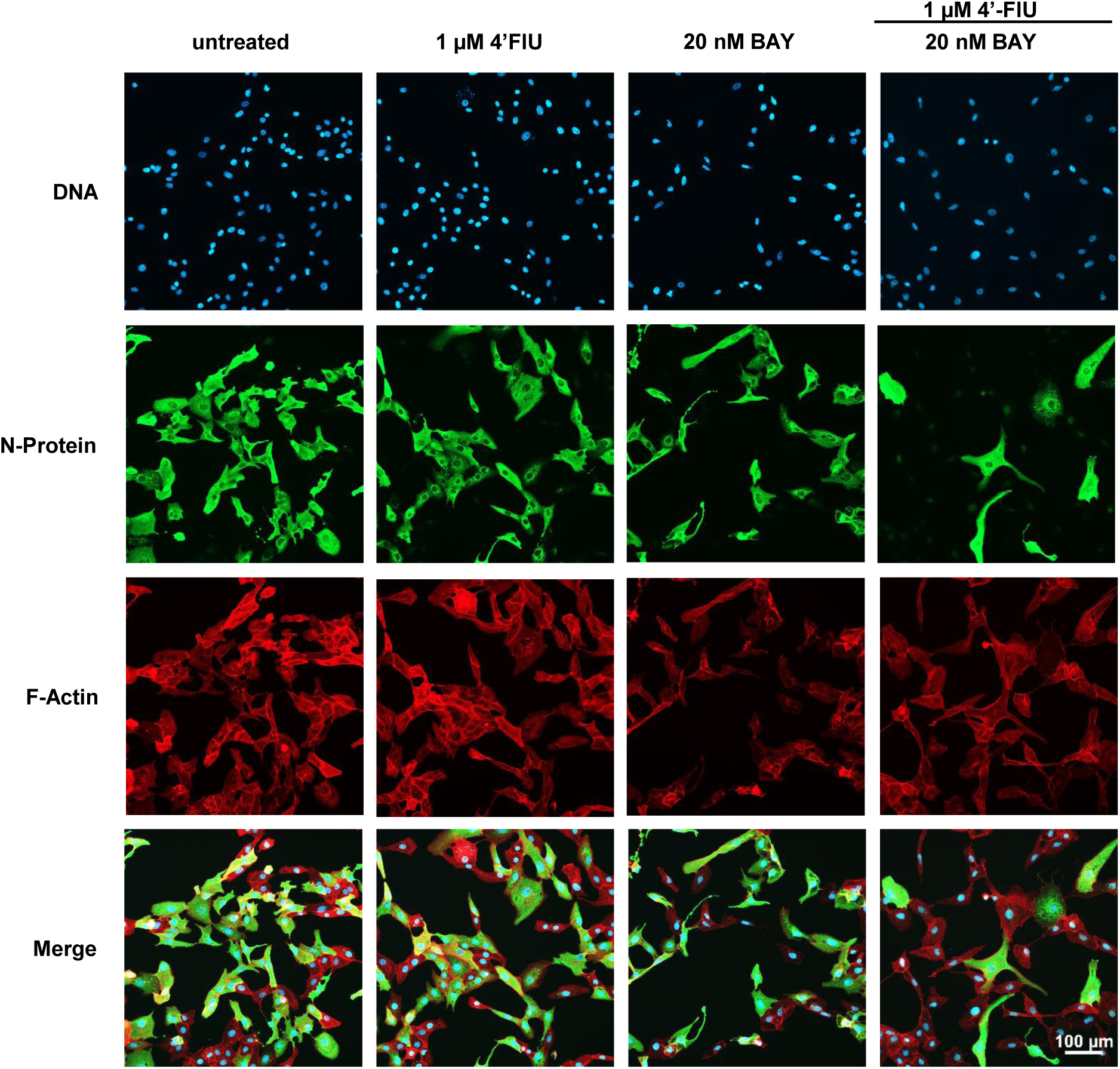
provides the single-color images corresponding to the overlays shown in Figure 5G.

**S12.**
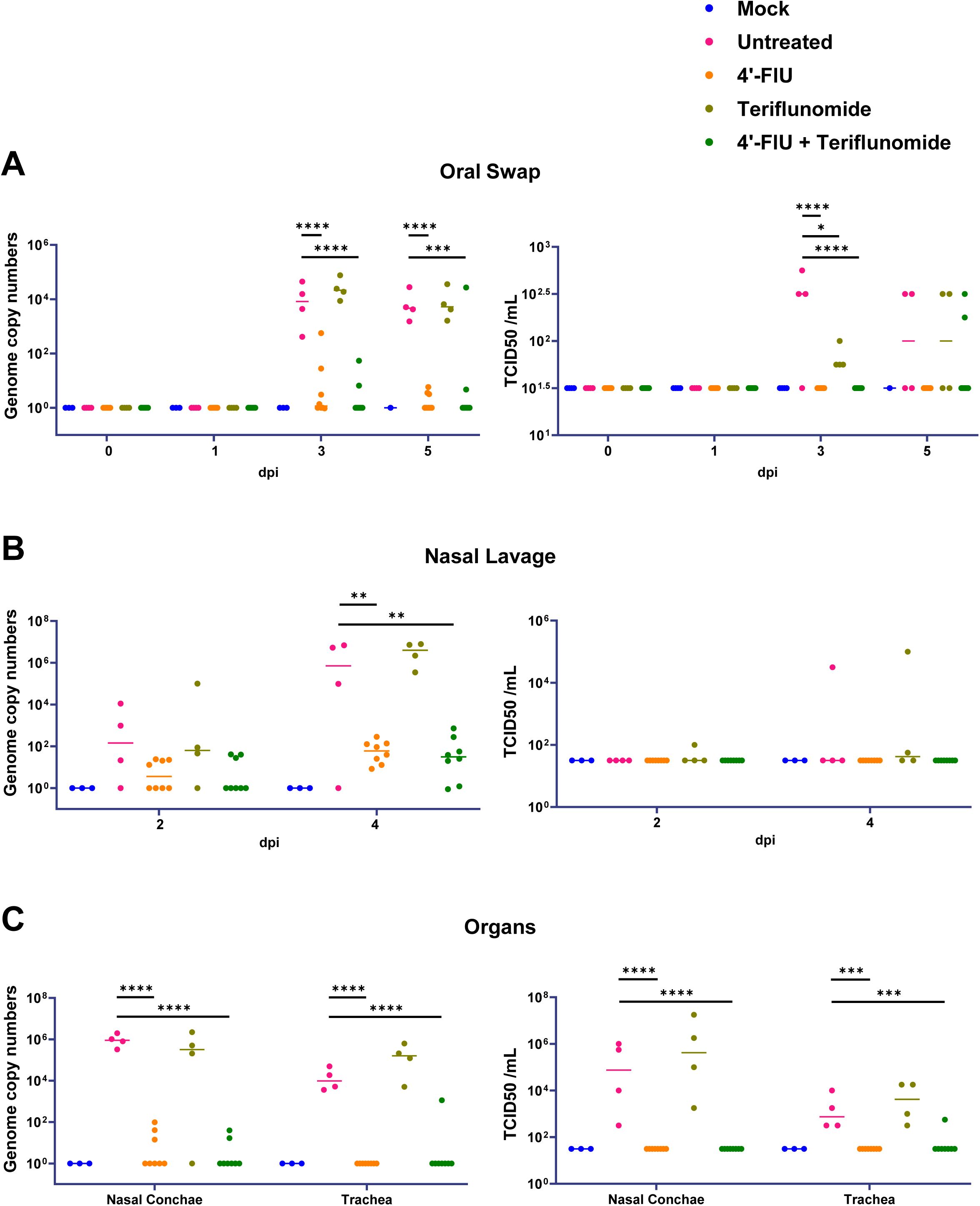
Virus genome copy numbers, determined by qRT-PCR (left), and infectious particles (TCID50, right) in oral swaps (A) and nasal lavage samples (B) obtained at the indicated days post infection (dpi). (C), Post mortem determinations of SARS-CoV-2 RNA and infectious particles in nasal conchae and tracheae (cf. Figure 6B, C).

**S13.**
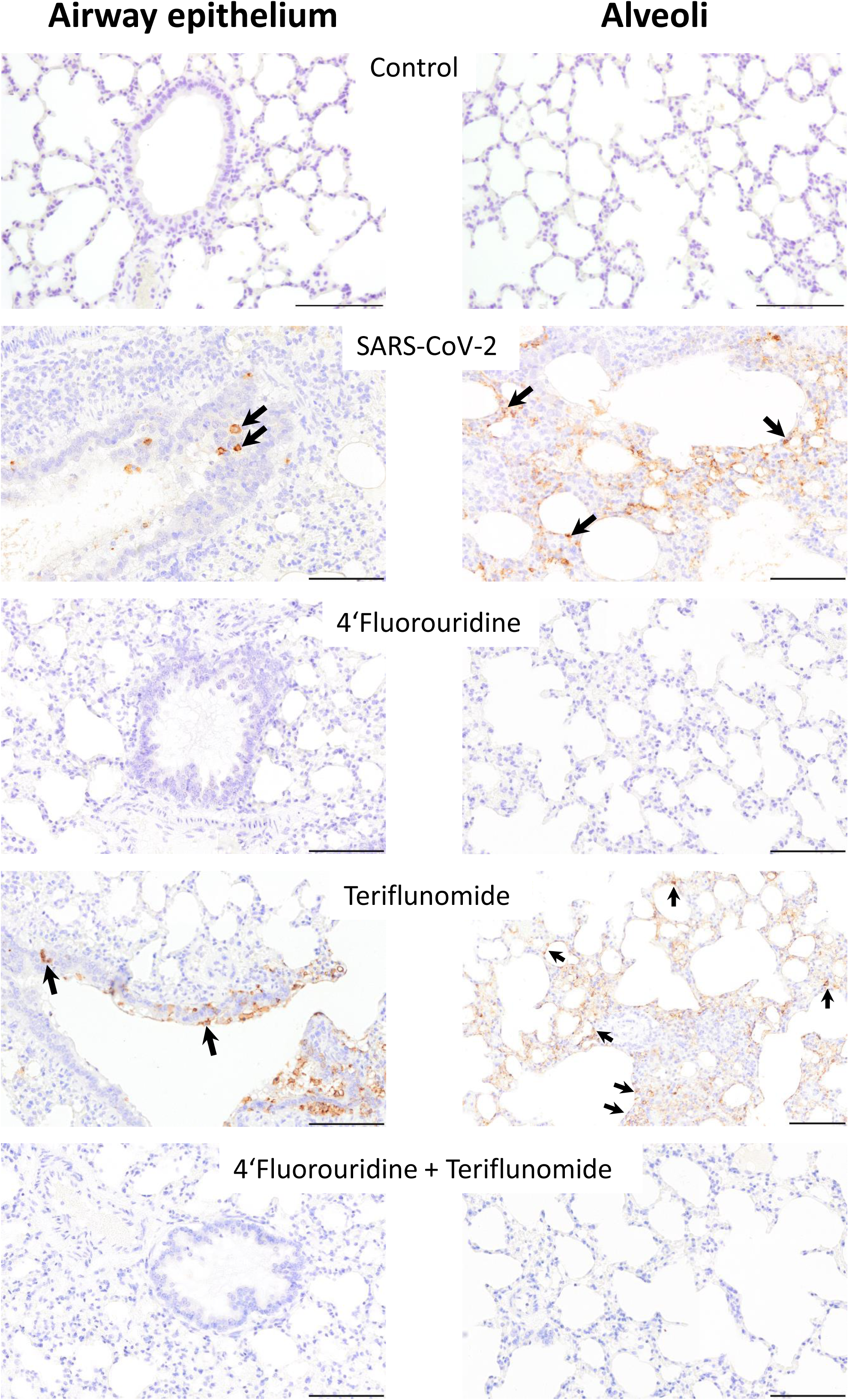
IHC images that correspond to Figure 6D. Comparison of SARS-CoV-2 nucleoprotein (NP) immunoreactivity: In the untreated and teriflunomide treated SARS-CoV-2 hamsters, strong immunolabeling (arrows) of airway epithelial cells and alveolar lining cells, including type II pneumocytes, was present. No immunoreactivity was noted in the 4′-FlU and combination treatment group. Chromogen: 3,3’-diaminobenzidine, hematoxylin counterstain. Bar=100µm.

**S14.**
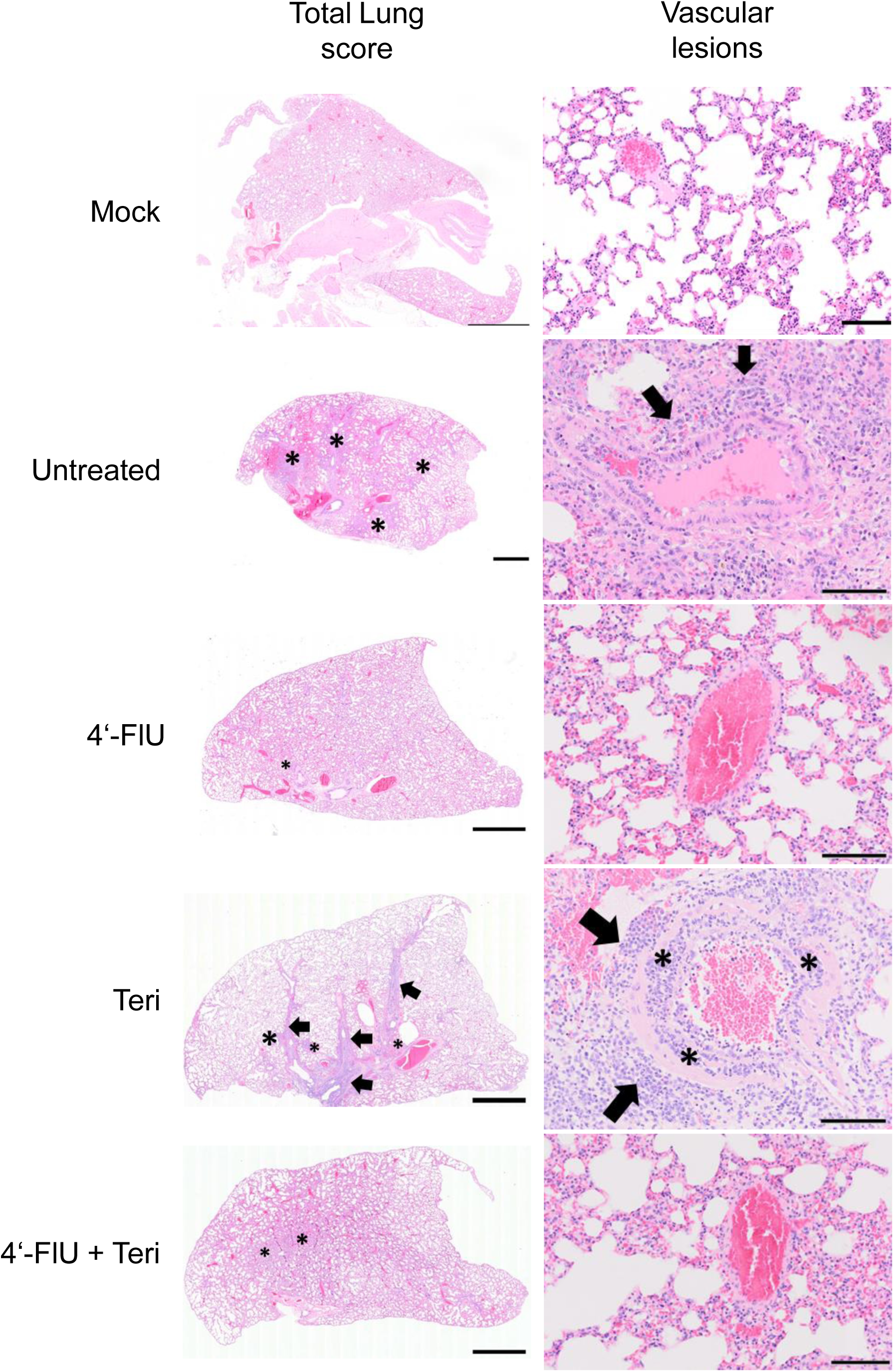
displays the pathomorphological analyses summarized in Figure 6E. Comparison of total lung scores (including alveolar, vascular and airway scores): Lungs of the non-infected control hamsters showed minimal edema and/or bleeding. In contrast, multifocal consolidation, alveolitis and atelectasis (asterisk) were present in the SARS-CoV-2 infected hamsters without treatment. Lesions were nearly completely resolved in both 4′-FlU treated groups, while teriflunomide alone showed some improvement of alveolar and vascular pathology but pronounced airway lesions (arrows). Bar = 2mm Comparison of vascular lesions: In SARS-CoV-2 infected untreated and teriflunomide-treated hamsters, marked vascular lesions were present. Lesions consisted of mononuclear infiltration (black arrows) and perivascular edema. In the only teriflunomide-treated hamsters, pronounced intramural infiltration of mononuclear cells and heterophils (asterisk) was detectable. In the 4′-FlU-treated and combination-treated hamsters, vascular lesions were nearly absent. There were no lesions in the non-infected and untreated control group. Hematoxylin-eosin. Bar = 100µm

**S15.**
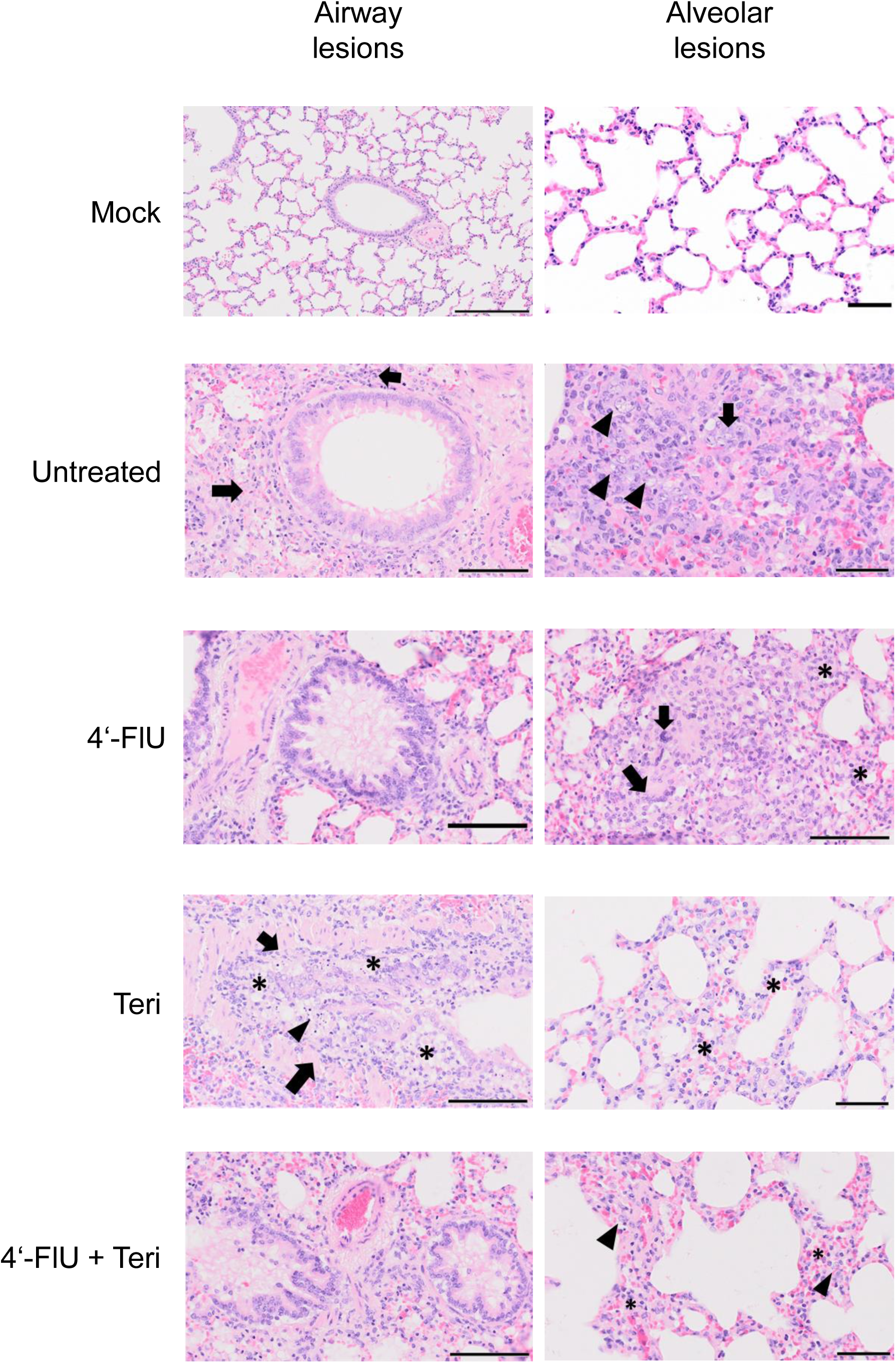
displays additional pathomorphological analyses, as summarized in Figure 6E. Comparison of airway lesions: In SARS-CoV-2 infected untreated and teriflunomide-treated hamsters, pronounced airway lesions were noted. Lesions consisted of peribronchial and peribronchiolar mononuclear infiltration (arrows). In addition, varying degrees of epithelial hyperplasia were present. Lesions were enhanced in the teriflunomide-treated hamsters that showed subepithelial edema (asterisks) and multifocal epithelial necrosis (arrowhead). Intraluminally, viable and degenerated heterophils as well as cellular debris were present. Lesions were nearly completely resolved in the 4′-FlU-treated and combination-treated hamsters. Hematoxylin-eosin. Bar = 100µm Comparison of alveolar lesions: In SARS-CoV-2 infected untreated hamsters, multifocal atelectasis was characterized by pronounced proliferation of atypical pneumocytes type II characterized by karyomegaly and anisokaryosis (arrowheads). In addition, there were multinucleated cells (arrows). Similar lesions were found in the 4′-FlU-treated hamsters though the total extent of lesions was markedly reduced. Teriflunomid-treated and combination-treated hamsters showed minimal lesions mostly consisting of mild alveolar edema and hemorrhage as well as scattered heterophils within the alveolar septae (asterisk). Occasionally, atypical cells were detected in the combination-treated group. Hematoxylin-eosin. Bar = 50µm

**S16.**
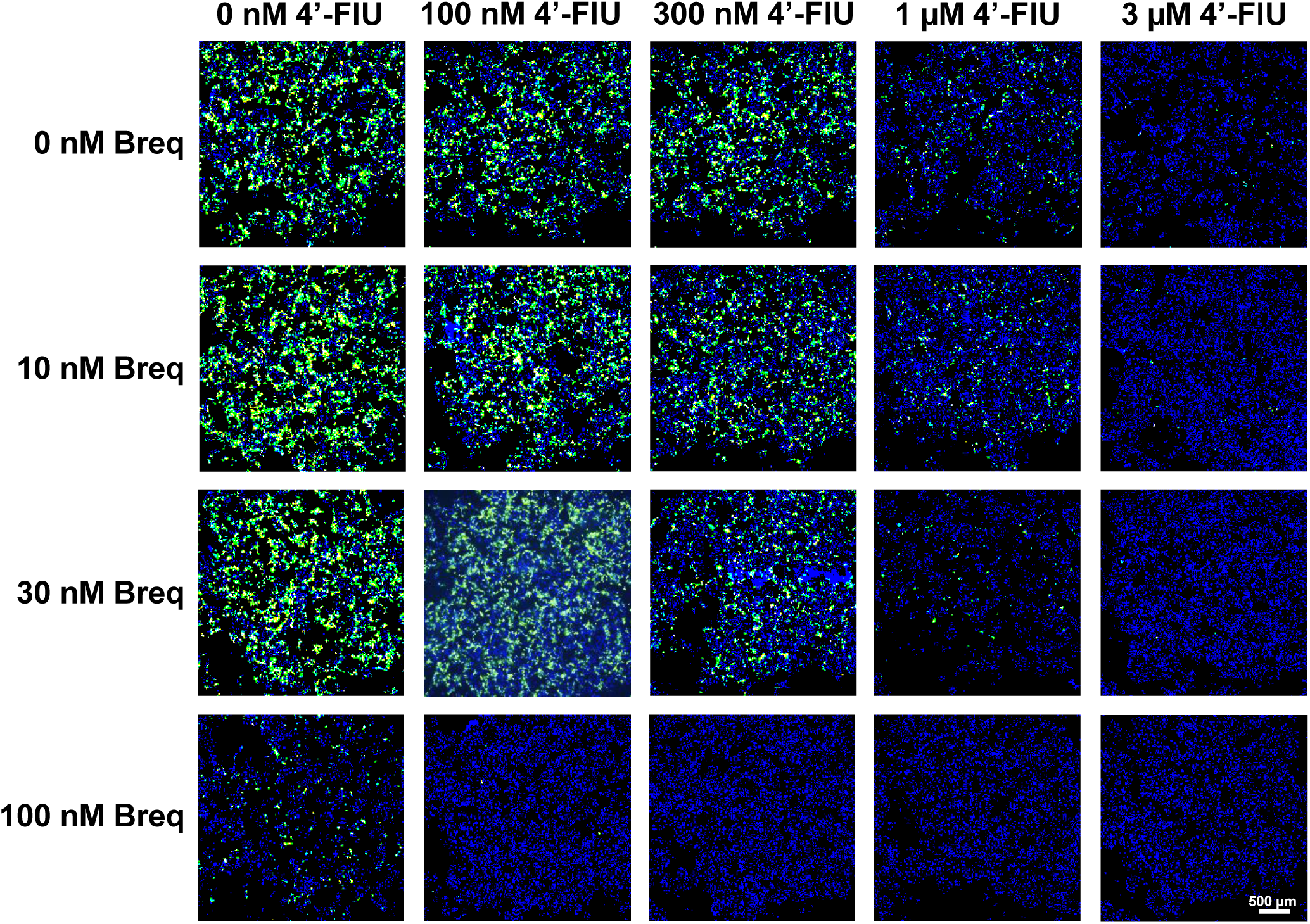
shows the green fluorescence of cells infected with recombinant EBOV and treated with 4′-FlU and brequinar, as in Figure 8A.

**S17.**
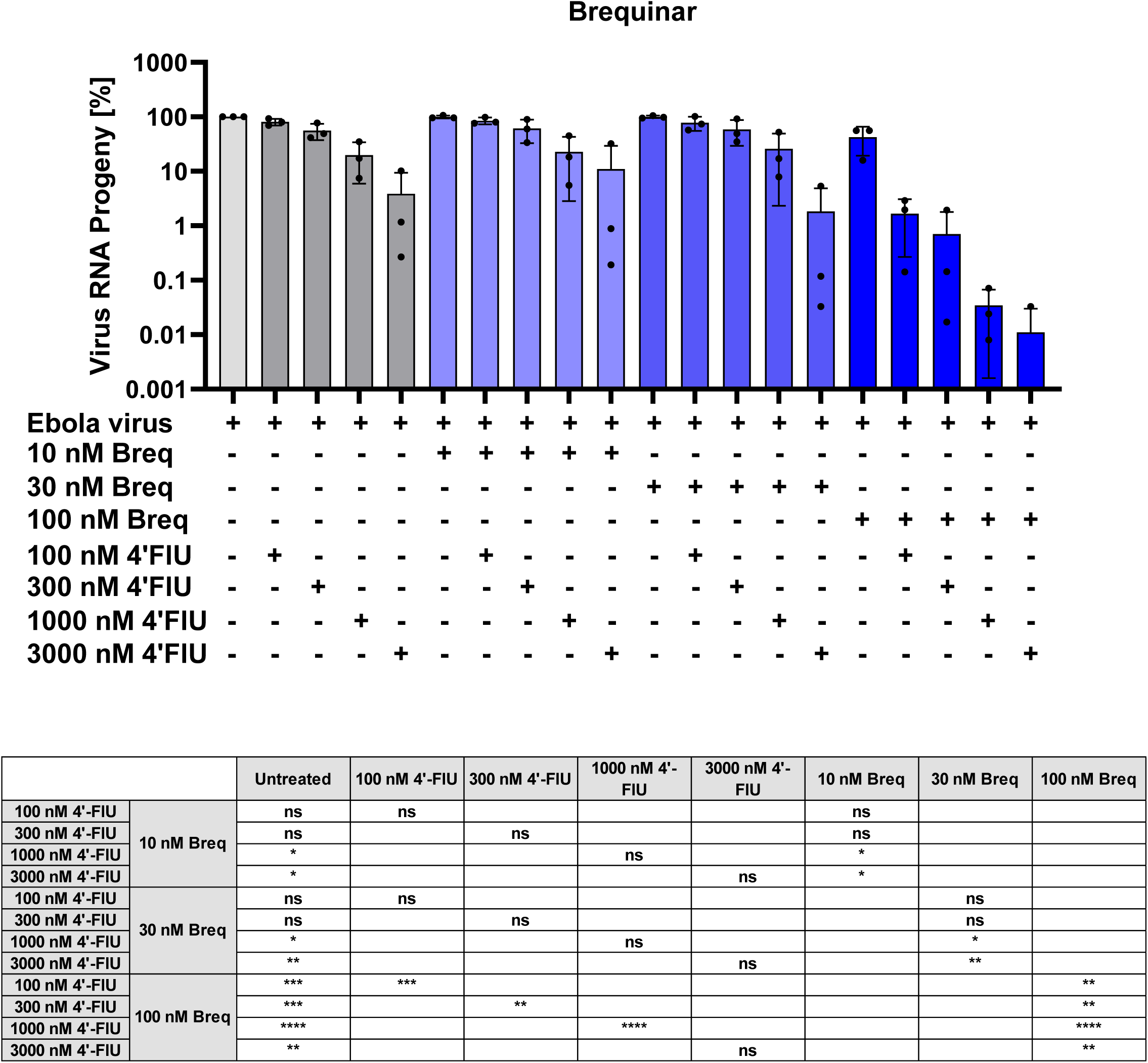
provides details on the quantitative analyses of EBOV infected cells (% infected cells), corresponding to Figure 8B.

## SUPPLEMENTAL TABLE

The Supplemental Table provides the raw data visualized in all Figures that display quantitative data. In the MS Excel file, the sheets are named after the figure panels they correspond to.

## REFERENCES

Agostini, M.L., Pruijssers, A.J., Chappell, J.D., Gribble, J., Lu, X., Andres, E.L., Bluemling, G.R., Lockwood, M.A., Sheahan, T.P., Sims, A.C., Natchus, M.G., Saindane, M., Kolykhalov, A.A., Painter, G.R., Baric, R.S. & Denison, M.R., 2019. Small-Molecule Antiviral β-d-N(4)-Hydroxycytidine Inhibits a Proofreading-Intact Coronavirus with a High Genetic Barrier to Resistance. Journal of virology 93. 10.1128/jvi.01348-19.

Aksu, M., Kumar, P., Guttler, T., Taxer, W., Gregor, K., Mußil, B., Rymarenko, O., Stegmann, K.M., Dickmanns, A., Gerber, S., Reineking, W., Schulz, C., Henneck, T., Mohamed, A., Pohlmann, G., Ramazanoglu, M., Mese, K., Groß, U., Ben-Yedidia, T., Ovadia, O., Fischer, D.W., Kamensky, M., Reichman, A., Baumgartner, W., von Kockritz-Blickwede, M., Dobbelstein, M. & Gorlich, D., 2024. Nanobodies to multiple spike variants and inhalation of nanobody-containing aerosols neutralize SARS-CoV-2 in cell culture and hamsters. Antiviral research 221, 105778. 10.1016/j.antiviral.2023.105778.

Almeida-Pinto, F., Pinto, R. & Rocha, J., 2024. Navigating the Complex Landscape of Ebola Infection Treatment: A Review of Emerging Pharmacological Approaches. Infectious diseases and therapy 13, 21–55. 10.1007/s40121-023-00913-y.

Amati, F., Tonutti, A., Huston, J. & Dela Cruz, C.S., 2023. Glucocorticoid Therapy in COVID-19. Seminars in respiratory and critical care medicine 44, 100–117. 10.1055/s-0042-1759778.

Amaya, M., Cheng, H., Borisevich, V., Navaratnarajah, C.K., Cattaneo, R., Cooper, L., Moore, T.W., Gaisina, I.N., Geisbert, T.W., Rong, L. & Broder, C.C., 2021. A recombinant Cedar virus based high-throughput screening assay for henipavirus antiviral discovery. Antiviral research 193, 105084. 10.1016/j.antiviral.2021.105084.

Armando, F., Beythien, G., Kaiser, F.K., Allnoch, L., Heydemann, L., Rosiak, M., Becker, S., Gonzalez-Hernandez, M., Lamers, M.M., Haagmans, B.L., Guilfoyle, K., van Amerongen, G., Ciurkiewicz, M., Osterhaus, A. & Baumgartner, W., 2022. SARS-CoV-2 Omicron variant causes mild pathology in the upper and lower respiratory tract of hamsters. Nature communications 13, 3519. 10.1038/s41467-022-31200-y.

Bankhead, P., Loughrey, M.B., Fernandez, J.A., Dombrowski, Y., McArt, D.G., Dunne, P.D., McQuaid, S., Gray, R.T., Murray, L.J., Coleman, H.G., James, J.A., Salto-Tellez, M. & Hamilton, P.W., 2017. QuPath: Open source software for digital pathology image analysis. Scientific reports 7, 16878. 10.1038/s41598-017-17204-5.

Blaurock, C., Breithaupt, A., Weber, S., Wylezich, C., Keller, M., Mohl, B.P., Gorlich, D., Groschup, M.H., Sadeghi, B., Hoper, D., Mettenleiter, T.C. & Balkema-Buschmann, A., 2022. Compellingly high SARS-CoV-2 susceptibility of Golden Syrian hamsters suggests multiple zoonotic infections of pet hamsters during the COVID-19 pandemic. Scientific reports 12, 15069. 10.1038/s41598-022-19222-4.

Buchholz, U.J., Finke, S. & Conzelmann, K.K., 1999. Generation of bovine respiratory syncytial virus (BRSV) from cDNA: BRSV NS2 is not essential for virus replication in tissue culture, and the human RSV leader region acts as a functional BRSV genome promoter. Journal of virology 73, 251–259. 10.1128/jvi.73.1.251-259.1999.

Burrough, E.R., Magstadt, D.R., Petersen, B., Timmermans, S.J., Gauger, P.C., Zhang, J., Siepker, C., Mainenti, M., Li, G., Thompson, A.C., Gorden, P.J., Plummer, P.J. & Main, R., 2024. Highly Pathogenic Avian Influenza A(H5N1) Clade 2.3.4.4b Virus Infection in Domestic Dairy Cattle and Cats, United States, 2024. Emerging infectious diseases 30, 1335–1343. 10.3201/eid3007.240508.

Calle-Hernandez, D.M., Hoyos-Salazar, V. & Bonilla-Aldana, D.K., 2023. Prevalence of the H5N8 influenza virus in birds: Systematic review with meta-analysis. Travel medicine and infectious disease 51, 102490. 10.1016/j.tmaid.2022.102490.

Chen, S.F., Ruben, R.L. & Dexter, D.L., 1986. Mechanism of action of the novel anticancer agent 6-fluoro-2-(2’-fluoro-1,1’-biphenyl-4-yl)-3-methyl-4-quinolinecarbo xylic acid sodium salt (NSC 368390): inhibition of de novo pyrimidine nucleotide biosynthesis. Cancer research 46, 5014–5019.

Christian, S., Merz, C., Evans, L., Gradl, S., Seidel, H., Friberg, A., Eheim, A., Lejeune, P., Brzezinka, K., Zimmermann, K., Ferrara, S., Meyer, H., Lesche, R., Stoeckigt, D., Bauser, M., Haegebarth, A., Sykes, D.B., Scadden, D.T., Losman, J.A. & Janzer, A., 2019. The novel dihydroorotate dehydrogenase (DHODH) inhibitor BAY 2402234 triggers differentiation and is effective in the treatment of myeloid malignancies. Leukemia 33, 2403–2415. 10.1038/s41375-019-0461-5.

Corman, V.M., Landt, O., Kaiser, M., Molenkamp, R., Meijer, A., Chu, D.K., Bleicker, T., Brunink, S., Schneider, J., Schmidt, M.L., Mulders, D.G., Haagmans, B.L., van der Veer, B., van den Brink, S., Wijsman, L., Goderski, G., Romette, J.L., Ellis, J., Zambon, M., Peiris, M., Goossens, H., Reusken, C., Koopmans, M.P. & Drosten, C., 2020. Detection of 2019 novel coronavirus (2019-nCoV) by real-time RT-PCR. Euro surveillance : bulletin Europeen sur les maladies transmissibles = European communicable disease bulletin 25. 10.2807/1560-7917.es.2020.25.3.2000045.

De Clercq, E., 2004. Antiviral drugs in current clinical use. Journal of clinical virology : the official publication of the Pan American Society for Clinical Virology 30, 115–133. 10.1016/j.jcv.2004.02.009.

De Clercq, E., 2023. The development of BVDU: An odyssey. Antiviral chemistry & chemotherapy 31, 20402066231152971. 10.1177/20402066231152971.

de Mariz, E.M.L.S., 2023. The synergy between nucleotide biosynthesis inhibitors and antiviral nucleosides: New opportunities against viral infections? Archiv der Pharmazie 356, e2200217. 10.1002/ardp.202200217.

Durrwald, R., Wedde, M., Biere, B., Oh, D.Y., Heßler-Klee, M., Geidel, C., Volmer, R., Hauri, A.M., Gerst, K., Thurmer, A., Appelt, S., Reiche, J., Duwe, S., Buda, S., Wolff, T. & Haas, W., 2020. Zoonotic infection with swine A/H1(av)N1 influenza virus in a child, Germany, June 2020. Euro surveillance : bulletin Europeen sur les maladies transmissibles = European communicable disease bulletin 25. 10.2807/1560-7917.es.2020.25.42.2001638.

Garcia-Blanco, M.A., Ooi, E.E. & Sessions, O.M., 2022. RNA Viruses, Pandemics and Anticipatory Preparedness. Viruses 14. 10.3390/v14102176.

Geraghty, R.J., Aliota, M.T. & Bonnac, L.F., 2021. Broad-Spectrum Antiviral Strategies and Nucleoside Analogues. Viruses 13. 10.3390/v13040667.

Gong, M., Yang, Y., Huang, Y., Gan, T., Wu, Y., Gao, H., Li, Q., Nie, J., Huang, W., Wang, Y., Zhang, R., Zhong, J., Deng, F., Rao, Y. & Ding, Q., 2021. Novel quinolone derivatives targeting human dihydroorotate dehydrogenase suppress Ebola virus infection in vitro. Antiviral research 194, 105161. 10.1016/j.antiviral.2021.105161.

Gordon, C.J., Tchesnokov, E.P., Schinazi, R.F. & Gotte, M., 2021. Molnupiravir promotes SARS-CoV-2 mutagenesis via the RNA template. The Journal of biological chemistry 297, 100770. 10.1016/j.jbc.2021.100770.

Graaf, A., Petric, P.P., Sehl-Ewert, J., Henritzi, D., Breithaupt, A., King, J., Pohlmann, A., Deutskens, F., Beer, M., Schwemmle, M. & Harder, T., 2022. Cold-passaged isolates and bat-swine influenza a chimeric viruses as modified live-attenuated vaccines against influenza a viruses in pigs. Vaccine 40, 6255–6270. 10.1016/j.vaccine.2022.09.013.

Greene, S., Watanabe, K., Braatz-Trulson, J. & Lou, L., 1995. Inhibition of dihydroorotate dehydrogenase by the immunosuppressive agent leflunomide. Biochemical pharmacology 50, 861–867. 10.1016/0006-2952(95)00255-x.

Hahn, F., Wangen, C., Hage, S., Peter, A.S., Dobler, G., Hurst, B., Julander, J., Fuchs, J., Ruzsics, Z., Oberla, K., Jack, H.M., Ptak, R., Muehler, A., Groppel, M., Vitt, D., Peelen, E., Kohlhof, H. & Marschall, M., 2020. IMU-838, a Developmental DHODH Inhibitor in Phase II for Autoimmune Disease, Shows Anti-SARS-CoV-2 and Broad-Spectrum Antiviral Efficacy In Vitro. Viruses 12. 10.3390/v12121394.

Harcourt, J.L., Caidi, H., Anderson, L.J. & Haynes, L.M., 2011. Evaluation of the Calu-3 cell line as a model of in vitro respiratory syncytial virus infection. Journal of virological methods 174, 144–149. 10.1016/j.jviromet.2011.03.027.

Hassan, K.E., Ahrens, A.K., Ali, A., El-Kady, M.F., Hafez, H.M., Mettenleiter, T.C., Beer, M. & Harder, T., 2022. Improved Subtyping of Avian Influenza Viruses Using an RT-qPCR-Based Low Density Array: ’Riems Influenza a Typing Array’, Version 2 (RITA-2). Viruses 14. 10.3390/v14020415.

Henritzi, D., Petric, P.P., Lewis, N.S., Graaf, A., Pessia, A., Starick, E., Breithaupt, A., Strebelow, G., Luttermann, C., Parker, L.M.K., Schroder, C., Hammerschmidt, B., Herrler, G., Beilage, E.G., Stadlbauer, D., Simon, V., Krammer, F., Wacheck, S., Pesch, S., Schwemmle, M., Beer, M. & Harder, T.C., 2020. Surveillance of European Domestic Pig Populations Identifies an Emerging Reservoir of Potentially Zoonotic Swine Influenza A Viruses. Cell host & microbe 28, 614–627.e616. 10.1016/j.chom.2020.07.006.

Hillary, V.E. & Ceasar, S.A., 2023. An update on COVID-19: SARS-CoV-2 variants, antiviral drugs, and vaccines. Heliyon 9, e13952. 10.1016/j.heliyon.2023.e13952.

Hume, A.J., Heiden, B., Olejnik, J., Suder, E.L., Ross, S., Scoon, W.A., Bullitt, E., Ericsson, M., White, M.R., Turcinovic, J., Thao, T.T.N., Hekman, R.M., Kaserman, J.E., Huang, J., Alysandratos, K.D., Toth, G.E., Jakab, F., Kotton, D.N., Wilson, A.A., Emili, A., Thiel, V., Connor, J.H., Kemenesi, G., Cifuentes, D. & Muhlberger, E., 2022. Recombinant Lloviu virus as a tool to study viral replication and host responses. PLoS pathogens 18, e1010268. 10.1371/journal.ppat.1010268.

Ianevski, A., Giri, A.K. & Aittokallio, T., 2020. SynergyFinder 2.0: visual analytics of multi-drug combination synergies. Nucleic acids research 48, W488–W493. 10.1093/nar/gkaa216.

Jacob, S.T., Crozier, I., Fischer, W.A., 2nd, Hewlett, A., Kraft, C.S., Vega, M.A., Soka, M.J., Wahl, V., Griffiths, A., Bollinger, L. & Kuhn, J.H., 2020. Ebola virus disease. Nature reviews. Disease primers 6, 13. 10.1038/s41572-020-0147-3.

Kabinger, F., Stiller, C., Schmitzova, J., Dienemann, C., Kokic, G., Hillen, H.S., Hobartner, C. & Cramer, P., 2021. Mechanism of molnupiravir-induced SARS-CoV-2 mutagenesis. Nature structural & molecular biology 28, 740–746. 10.1038/s41594-021-00651-0.

Karber, G., 1931. Beitrag zur kollektiven Behandlung pharmakologischer Reihenversuche. Naunyn-Schmiedebergs Archiv fur experimentelle Pathologie und Pharmakologie 162, 480–483. 10.1007/BF01863914.

Klotz, L., Eschborn, M., Lindner, M., Liebmann, M., Herold, M., Janoschka, C., Torres Garrido, B., Schulte-Mecklenbeck, A., Gross, C.C., Breuer, J., Hundehege, P., Posevitz, V., Pignolet, B., Nebel, G., Glander, S., Freise, N., Austermann, J., Wirth, T., Campbell, G.R., Schneider-Hohendorf, T., Eveslage, M., Brassat, D., Schwab, N., Loser, K., Roth, J., Busch, K.B., Stoll, M., Mahad, D.J., Meuth, S.G., Turner, T., Bar-Or, A. & Wiendl, H., 2019. Teriflunomide treatment for multiple sclerosis modulates T cell mitochondrial respiration with affinity-dependent effects. Science translational medicine 11. 10.1126/scitranslmed.aao5563.

Krammer, F., Smith, G.J.D., Fouchier, R.A.M., Peiris, M., Kedzierska, K., Doherty, P.C., Palese, P., Shaw, M.L., Treanor, J., Webster, R.G. & Garcfa-Sastre, A., 2018. Influenza. Nature reviews. Disease primers 4, 3. 10.1038/s41572-018-0002-y.

Kumar, P., Zhang, X., Shaha, R., Kschischo, M. & Dobbelstein, M., 2024. Identification of antibody-resistant SARS-CoV-2 mutants via N4-Hydroxycytidine mutagenesis. Antiviral research 231, 106006. 10.1016/j.antiviral.2024.106006.

Laing, E.D., Amaya, M., Navaratnarajah, C.K., Feng, Y.R., Cattaneo, R., Wang, L.F. & Broder, C.C., 2018. Rescue and characterization of recombinant cedar virus, a non-pathogenic Henipavirus species. Virology journal 15, 56. 10.1186/s12985-018-0964-0.

Lawitz, E., Jacobson, I.M., Nelson, D.R., Zeuzem, S., Sulkowski, M.S., Esteban, R., Brainard, D., McNally, J., Symonds, W.T., McHutchison, J.G., Dieterich, D. & Gane, E., 2015. Development of sofosbuvir for the treatment of hepatitis C virus infection. Annals of the New York Academy of Sciences 1358, 56–67. 10.1111/nyas.12832.

Leban, J. & Vitt, D., 2011. Human dihydroorotate dehydrogenase inhibitors, a novel approach for the treatment of autoimmune and inflammatory diseases. Arzneimittel-Forschung 61, 66–72. 10.1055/s-0031-1296169.

Li, H., Kim, J.V. & Pickering, B.S., 2023. Henipavirus zoonosis: outbreaks, animal hosts and potential new emergence. Frontiers in microbiology 14, 1167085. 10.3389/fmicb.2023.1167085.

Li, J., Takeda, M., Imahatakenaka, M. & Ikeda, M., 2024. Identification of dihydroorotate dehydrogenase inhibitor, vidofludimus, as a potent and novel inhibitor for influenza virus. Journal of medical virology 96, e29372. 10.1002/jmv.29372.

Lieber, C.M., Aggarwal, M., Yoon, J.J., Cox, R.M., Kang, H.J., Sourimant, J., Toots, M., Johnson, S.K., Jones, C.A., Sticher, Z.M., Kolykhalov, A.A., Saindane, M.T., Tompkins, S.M., Planz, O., Painter, G.R., Natchus, M.G., Sakamoto, K. & Plemper, R.K., 2023. 4’-Fluorouridine mitigates lethal infection with pandemic human and highly pathogenic avian influenza viruses. PLoS pathogens 19, e1011342. 10.1371/journal.ppat.1011342.

Lieber, C.M., Kang, H.J., Aggarwal, M., Lieberman, N.A., Sobolik, E.B., Yoon, J.J., Natchus, M.G., Cox, R.M., Greninger, A.L. & Plemper, R.K., 2024. Influenza A virus resistance to 4’-fluorouridine coincides with viral attenuation in vitro and in vivo. PLoS pathogens 20, e1011993. 10.1371/journal.ppat.1011993.

Lieber, C.M. & Plemper, R.K., 2022. 4’-Fluorouridine Is a Broad-Spectrum Orally Available First-Line Antiviral That May Improve Pandemic Preparedness. DNA and cell biology 41, 699–704. 10.1089/dna.2022.0312.

Lo, M.K., Amblard, F., Flint, M., Chatterjee, P., Kasthuri, M., Li, C., Russell, O., Verma, K., Bassit, L., Schinazi, R.F., Nichol, S.T. & Spiropoulou, C.F., 2020. Potent in vitro activity of β-D-4’-chloromethyl-2’-deoxy-2’-fluorocytidine against Nipah virus. Antiviral research 175, 104712. 10.1016/j.antiviral.2020.104712.

Lo, M.K., Jordan, P.C., Stevens, S., Tam, Y., Deval, J., Nichol, S.T. & Spiropoulou, C.F., 2018. Susceptibility of paramyxoviruses and filoviruses to inhibition by 2’-monofluoro-and 2’-difluoro-4’-azidocytidine analogs. Antiviral research 153, 101–113. 10.1016/j.antiviral.2018.03.009.

Luthra, P., Naidoo, J., Pietzsch, C.A., De, S., Khadka, S., Anantpadma, M., Williams, C.G., Edwards, M.R., Davey, R.A., Bukreyev, A., Ready, J.M. & Basler, C.F., 2018. Inhibiting pyrimidine biosynthesis impairs Ebola virus replication through depletion of nucleoside pools and activation of innate immune responses. Antiviral research 158, 288–302. 10.1016/j.antiviral.2018.08.012.

Marsh, G.A., de Jong, C., Barr, J.A., Tachedjian, M., Smith, C., Middleton, D., Yu, M., Todd, S., Foord, A.J., Haring, V., Payne, J., Robinson, R., Broz, I., Crameri, G., Field, H.E. & Wang, L.F., 2012. Cedar virus: a novel Henipavirus isolated from Australian bats. PLoS pathogens 8, e1002836. 10.1371/journal.ppat.1002836.

Martin, S., Chiramel, A.I., Schmidt, M.L., Chen, Y.C., Whitt, N., Watt, A., Dunham, E.C., Shifflett, K., Traeger, S., Leske, A., Buehler, E., Martellaro, C., Brandt, J., Wendt, L., Muller, A., Peitsch, S., Best, S.M., Stech, J., Finke, S., Romer-Oberdorfer, A., Groseth, A., Feldmann, H. & Hoenen, T., 2018. A genome-wide siRNA screen identifies a druggable host pathway essential for the Ebola virus life cycle. Genome medicine 10, 58. 10.1186/s13073-018-0570-1.

Meganck, R.M. & Baric, R.S., 2021. Developing therapeutic approaches for twenty-first-century emerging infectious viral diseases. Nature medicine 27, 401–410. 10.1038/s41591-021-01282-0.

Mourad, A., Thibault, D., Holland, T.L., Yang, S., Young, A.R., Arnold Egloff, S.A. & Thomas, L.E., 2023. Dexamethasone for Inpatients With COVID-19 in a National Cohort. JAMA network open 6, e238516. 10.1001/jamanetworkopen.2023.8516.

Mungall, B.A., Middleton, D., Crameri, G., Bingham, J., Halpin, K., Russell, G., Green, D., McEachern, J., Pritchard, L.I., Eaton, B.T., Wang, L.F., Bossart, K.N. & Broder, C.C., 2006. Feline model of acute nipah virus infection and protection with a soluble glycoprotein-based subunit vaccine. Journal of virology 80, 12293–12302. 10.1128/jvi.01619-06.

Niwa, H., Yamamura, K. & Miyazaki, J., 1991. Efficient selection for high-expression transfectants with a novel eukaryotic vector. Gene 108, 193–199. 10.1016/0378-1119(91)90434-d.

Parys, A., Vandoorn, E., King, J., Graaf, A., Pohlmann, A., Beer, M., Harder, T. & Van Reeth, K., 2021. Human Infection with Eurasian Avian-Like Swine Influenza A(H1N1) Virus, the Netherlands, September 2019. Emerging infectious diseases 27, 939–943. 10.3201/eid2703.201863.

Patel, K., Kirkpatrick, C.M., Nieforth, K.A., Chanda, S., Zhang, Q., McClure, M., Fry, J., Symons, J.A., Blatt, L.M., Beigelman, L., DeVincenzo, J.P., Huntjens, D.R. & Smith, P.F., 2019. Respiratory syncytial virus-A dynamics and the effects of lumicitabine, a nucleoside viral replication inhibitor, in experimentally infected humans. The Journal of antimicrobial chemotherapy 74, 442–452. 10.1093/jac/dky415.

Ramakrishnan, M.A., 2016. Determination of 50% endpoint titer using a simple formula. World journal of virology 5, 85–86. 10.5501/wjv.v5.i2.85.

Schnell, M.J., Mebatsion, T. & Conzelmann, K.K., 1994. Infectious rabies viruses from cloned cDNA. The EMBO journal 13, 4195–4203. 10.1002/j.1460-2075.1994.tb06739.x.

Schultz, D.C., Johnson, R.M., Ayyanathan, K., Miller, J., Whig, K., Kamalia, B., Dittmar, M., Weston, S., Hammond, H.L., Dillen, C., Ardanuy, J., Taylor, L., Lee, J.S., Li, M., Lee, E., Shoffler, C., Petucci, C., Constant, S., Ferrer, M., Thaiss, C.A., Frieman, M.B. & Cherry, S., 2022. Pyrimidine inhibitors synergize with nucleoside analogues to block SARS-CoV-2. Nature 604, 134–140. 10.1038/s41586-022-04482-x.

Sheahan, T.P., Sims, A.C., Zhou, S., Graham, R.L., Pruijssers, A.J., Agostini, M.L., Leist, S.R., Schafer, A., Dinnon, K.H., 3rd, Stevens, L.J., Chappell, J.D., Lu, X., Hughes, T.M., George, A.S., Hill, C.S., Montgomery, S.A., Brown, A.J., Bluemling, G.R., Natchus, M.G., Saindane, M., Kolykhalov, A.A., Painter, G., Harcourt, J., Tamin, A., Thornburg, N.J., Swanstrom, R., Denison, M.R. & Baric, R.S., 2020. An orally bioavailable broad-spectrum antiviral inhibits SARS-CoV-2 in human airway epithelial cell cultures and multiple coronaviruses in mice. Science translational medicine 12. 10.1126/scitranslmed.abb5883.

Sibille, G., Luganini, A., Sainas, S., Boschi, D., Lolli, M.L. & Gribaudo, G., 2022. The Novel hDHODH Inhibitor MEDS433 Prevents Influenza Virus Replication by Blocking Pyrimidine Biosynthesis. Viruses 14. 10.3390/v14102281.

Simon, G., Larsen, L.E., Durrwald, R., Foni, E., Harder, T., Van Reeth, K., Markowska-Daniel, I., Reid, S.M., Dan, A., Maldonado, J., Huovilainen, A., Billinis, C., Davidson, I., Aguero, M., Vila, T., Herve, S., Breum, S., Chiapponi, C., Urbaniak, K., Kyriakis, C.S., Brown, I.H. & Loeffen, W., 2014. European surveillance network for influenza in pigs: surveillance programs, diagnostic tools and Swine influenza virus subtypes identified in 14 European countries from 2010 to 2013. PloS one 9, e115815. 10.1371/journal.pone.0115815.

Sourimant, J., Lieber, C.M., Aggarwal, M., Cox, R.M., Wolf, J.D., Yoon, J.J., Toots, M., Ye, C., Sticher, Z., Kolykhalov, A.A., Martinez-Sobrido, L., Bluemling, G.R., Natchus, M.G., Painter, G.R. & Plemper, R.K., 2022. 4’-Fluorouridine is an oral antiviral that blocks respiratory syncytial virus and SARS-CoV-2 replication. Science (New York, N.Y.) 375, 161–167. 10.1126/science.abj5508.

Spackman, E., Senne, D.A., Bulaga, L.L., Myers, T.J., Perdue, M.L., Garber, L.P., Lohman, K., Daum, L.T. & Suarez, D.L., 2003. Development of real-time RT-PCR for the detection of avian influenza virus. Avian diseases 47, 1079–1082. 10.1637/0005-2086-47.s3.1079.

Stegmann, K.M., Dickmanns, A., Gerber, S., Nikolova, V., Klemke, L., Manzini, V., Schlosser, D., Bierwirth, C., Freund, J., Sitte, M., Lugert, R., Salinas, G., Meister, T.L., Pfaender, S., Gorlich, D., Wollnik, B., Groß, U. & Dobbelstein, M., 2021. The folate antagonist methotrexate diminishes replication of the coronavirus SARS-CoV-2 and enhances the antiviral efficacy of remdesivir in cell culture models. Virus research 302, 198469. 10.1016/j.virusres.2021.198469.

Stegmann, K.M., Dickmanns, A., Heinen, N., Blaurock, C., Karrasch, T., Breithaupt, A., Klopfleisch, R., Uhlig, N., Eberlein, V., Issmail, L., Herrmann, S.T., Schreieck, A., Peelen, E., Kohlhof, H., Sadeghi, B., Riek, A., Speakman, J.R., Groß, U., Gorlich, D., Vitt, D., Muller, T., Grunwald, T., Pfaender, S., Balkema-Buschmann, A. & Dobbelstein, M., 2022. Inhibitors of dihydroorotate dehydrogenase cooperate with molnupiravir and N4-hydroxycytidine to suppress SARS-CoV-2 replication. iScience 25, 104293. 10.1016/j.isci.2022.104293.

Sweileh, W.M., 2017. Global research trends of World Health Organization’s top eight emerging pathogens. Globalization and health 13, 9. 10.1186/s12992-017-0233-9.

Toots, M., Yoon, J.J., Cox, R.M., Hart, M., Sticher, Z.M., Makhsous, N., Plesker, R., Barrena, A.H., Reddy, P.G., Mitchell, D.G., Shean, R.C., Bluemling, G.R., Kolykhalov, A.A., Greninger, A.L., Natchus, M.G., Painter, G.R. & Plemper, R.K., 2019. Characterization of orally efficacious influenza drug with high resistance barrier in ferrets and human airway epithelia. Science translational medicine 11. 10.1126/scitranslmed.aax5866.

Wahl, A., Gralinski, L.E., Johnson, C.E., Yao, W., Kovarova, M., Dinnon, K.H., 3rd, Liu, H., Madden, V.J., Krzystek, H.M., De, C., White, K.K., Gully, K., Schafer, A., Zaman, T., Leist, S.R., Grant, P.O., Bluemling, G.R., Kolykhalov, A.A., Natchus, M.G., Askin, F.B., Painter, G., Browne, E.P., Jones, C.D., Pickles, R.J., Baric, R.S. & Garcia, J.V., 2021. SARS-CoV-2 infection is effectively treated and prevented by EIDD-2801. Nature 591, 451–457. 10.1038/s41586-021-03312-w.

Westover, J.B., Jung, K.H., Alkan, C., Boardman, K.M., Van Wettere, A.J., Martens, C., Rojas, I., Hicks, P., Thomas, A.J., Saindane, M.T., Bluemling, G.R., Mao, S., Kolykhalov, A.A., Natchus, M.G., Bates, P., Painter, G.R., Ikegami, T. & Gowen, B.B., 2024. Modeling Heartland virus disease in mice and therapeutic intervention with 4’-fluorouridine. Journal of virology 98, e0013224. 10.1128/jvi.00132-24.

Xiong, R., Zhang, L., Li, S., Sun, Y., Ding, M., Wang, Y., Zhao, Y., Wu, Y., Shang, W., Jiang, X., Shan, J., Shen, Z., Tong, Y., Xu, L., Chen, Y., Liu, Y., Zou, G., Lavillete, D., Zhao, Z., Wang, R., Zhu, L., Xiao, G., Lan, K., Li, H. & Xu, K., 2020. Novel and potent inhibitors targeting DHODH are broad-spectrum antivirals against RNA viruses including newly-emerged coronavirus SARS-CoV-2. Protein & cell 11, 723–739. 10.1007/s13238-020-00768-w.

Yamamoto, T., Koyama, H., Kurajoh, M., Shoji, T., Tsutsumi, Z. & Moriwaki, Y., 2011. Biochemistry of uridine in plasma. Clinica chimica acta; international journal of clinical chemistry 412, 1712–1724. 10.1016/j.cca.2011.06.006.

Yarchoan, R., Klecker, R.W., Weinhold, K.J., Markham, P.D., Lyerly, H.K., Durack, D.T., Gelmann, E., Lehrman, S.N., Blum, R.M., Barry, D.W. &, et al., 1986. Administration of 3’-azido-3’-deoxythymidine, an inhibitor of HTLV-III/LAV replication, to patients with AIDS or AIDS-related complex. Lancet (London, England) 1, 575–580. 10.1016/s0140-6736(86)92808-4.

Zheng, Y., Li, S., Song, K., Ye, J., Li, W., Zhong, Y., Feng, Z., Liang, S., Cai, Z. & Xu, K., 2022. A Broad Antiviral Strategy: Inhibitors of Human DHODH Pave the Way for Host-Targeting Antivirals against Emerging and Re-Emerging Viruses. Viruses 14. 10.3390/v14050928.

Zibat, A., Zhang, X., Dickmanns, A., Stegmann, K.M., Dobbelstein, A.W., Alachram, H., Soliwoda, R., Salinas, G., Groß, U., Gorlich, D., Kschischo, M., Wollnik, B. & Dobbelstein, M., 2023. N4-hydroxycytidine, the active compound of Molnupiravir, promotes SARS-CoV-2 mutagenesis and escape from a neutralizing nanobody. iScience 26, 107786. 10.1016/j.isci.2023.107786.

